# Cluefish: mining the dark matter of transcriptional data series with over-representation analysis enhanced by aggregated biological prior knowledge

**DOI:** 10.1101/2024.12.18.627334

**Authors:** Ellis Franklin, Elise Billoir, Philippe Veber, Jérémie Ohanessian, Marie Laure Delignette-Muller, Sophie Martine Prud’homme

## Abstract

Interpreting transcriptomic data presents significant challenges, particularly in non-targeted approaches. While modern functional enrichment methods are well-suited for experimental designs involving two conditions, they are less applicable to data series. In this context, we developed Cluefish, a free and open-source, semi-automated R workflow designed for untargeted, comprehensive biological interpretation of transcriptomic data series. Cluefish applies over-representation analysis on pre-clustered protein-protein interaction networks, using clusters as anchors to identify smaller, more specific biological functions. Innovative features, including cluster merging and recovery of isolated genes through shared biological contexts, enable a more complete exploration of the data. We applied Cluefish to an in-house dataset with zebrafish exposed to a dose-gradient of dibutyl phthalate, and to two published toxicology datasets featuring different organisms. Combined with DRomics, a tool for dose-response analysis—Cluefish identified gene clusters deregulated at low doses and linked to biological functions overlooked by the standard approach. Notably, it revealed that retinoid signalling disruption may be the most sensitive pathway affected by dibutyl phthalate during zebrafish development, potentially leading to morphological changes. The Cluefish workflow aims to provide valuable clues for biological hypothesis generation and experimental validation. It is freely available at https://github.com/ellfran-7/cluefish.

**GRAPHICAL ABSTRACT:** 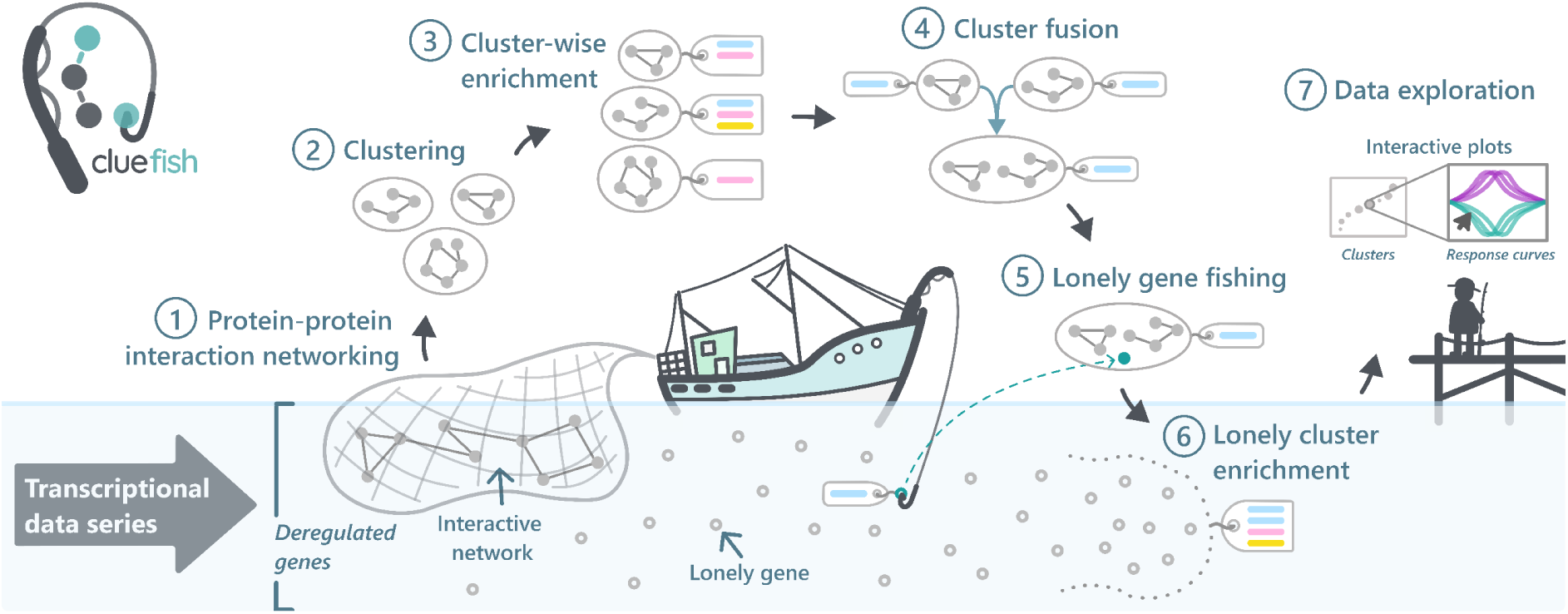

## INTRODUCTION

Advances in omics approaches has made the comprehensive measurement of DNA, RNA, proteins and metabolites in biological samples a standard practice (1). This has led to a massive increase in generated data, providing a goldmine for uncovering novel biological functions, genotype-phenotype relationships, and elucidating molecular mechanisms (2). However, the sheer volume presents significant challenges in both analysis and interpretation (3, 4).

In the context of transcriptomics, analyses yield extensive transcript lists that require impractical manual literature reviews. To address this, functional enrichment analysis, also known as pathway enrichment analysis, has become the standard approach. This method condenses large gene lists into more manageable and interpretable sets of biological functions or pathways (5). Functional enrichment relies on predefined gene sets representing molecular functions, biological processes or pathways, as outlined by databases like the Gene Ontology (GO) (6), pathway databases such as the Kyoto Encyclopedia of Genes and Genomes (KEGG) (7) and Wikipathways (WP) (8, 9), or experimentally derived gene sets available in the MSigDB (10, 11).

Functional enrichment methods have evolved over four generations. The first generation consists of Over-Representation Analysis (ORA) methods, which test whether a predefined gene set contains a disproportionately large number of deregulated genes compared to a uniform random sample of genes among all the genes detected in the experiment. In this case, the functional gene set is considered as enriched. The second generation includes Functional Class Scoring (FCS) methods, which evaluate whether genes within a set are concentrated at the top or bottom of a ranked list of all the genes from the experiment, based on expression changes such as fold-change (11). FCS improves on ORA by considering coordinated changes in functionally related gene sets, though it still treats genes as independent (12). The third generation introduced Pathway Topology (PT)-Based methods, which leverage pathway topology information from databases such as KEGG and Reactome (13). These methods utilise knowledge about gene positions, roles and interactions within pathways (14, 15). While PT-based methods improve on ORA and FCS by incorporating gene dependencies and interactions, they still view pathways as isolated entities and thus overlook pathway crosstalk (16). The most recent and fourth generation, Network Topology (NT)-based approaches, goes a step further by systematically considering pathway crosstalk within a network (17). NT-based methods utilise biological interaction networks, such as those based on protein-protein interactions (PPI) from databases like FunCoup and STRING (18, 19). These approaches examine changes in network structure and correlations to provide a more integrated understanding of pathway interactions, accounting for crosstalk and overlap between pathways— elements neglected by previous generations (20). However, despite their potential, NT-based methods are limited. This is because biological networks are often incomplete, with many missing or unreliable interactions, and these methods tend to focus solely on the network itself, which restricts the effectiveness and completeness of the analyses (17). As a result, earlier generations of functional enrichment methods still remain largely used in practice.

While these earlier methods are commonly applied to transcriptomic data with two or three conditions, dealing with data involving many ordered conditions—referred to as “data series”—poses substantial complexity, even before functional enrichment. Data series capture measurements collected over successive points in time, space, or under varying conditions (e.g., different doses of a contaminant). In dose-response (DR) designs, which explore the relationship between exposure and effect, differential gene expression (DEG) analysis, although often used, is not well-suited. DEG relies on the comparison of the response at each dose to the control (or sometimes on pairwise comparisons across all conditions), multiplying the number of tests, applying each to a subset of the data, and generating multiple fold-change values for each transcript. In contrast, DR omics analysis replaces DEG by leveraging the entire DR relationship for each transcript (21). Tools, like the DRomics R package (22), identify significantly deregulated transcripts and model their DR relationships. These models are used to characterise the response (whether monotonic or biphasic) and to calculate a benchmark dose (BMD) for each transcript. A BMD is a dose or concentration that produces a predetermined change in the response rate of an adverse effect, reflecting the sensitivity of a transcript to a given stressor (22). However, FCS, and PT-based enrichment, which rely on fold-change values, are not adapted for functional enrichment in a DR context. (12). In contrast, ORA is currently the only suitable method available to complement DR modelling, as it exclusively uses deregulated gene lists.

Alternative approaches exist for analysing data series, such as weighted gene co-expression network analysis (WGCNA) (23), which clusters genes based on expression correlation patterns across conditions. Other methods like maSigPro (24), RNFuzzyApp (25) and TRAP (26), tailored for time-series RNA-seq data, offer comprehensive analytical frameworks that include, among other utilities, identification of genes with significant temporal expression changes and clustering of expression patterns.

Nevertheless, to derive biological meaning from these identified gene sets and clusters, functional enrichment methods, predominantly ORA, remain necessary downstream steps, whether or not they are included in the workflow (27–31). However, ORA comes with several other limitations. The enrichment results, often derived from Fisher’s Exact Tests, tends to highlight broad biological functions. Indeed, when using the entire list of deregulated genes, the large test denominator makes it more challenging to achieve the statistical significance needed to identify smaller or lesser-known functions. This tendency is intrinsically tied to the granularity of gene sets in certain databases (e.g., GO). Gene sets may be overly specific, while others are too general, potentially compromising the relevance of the analysis (32, 33). Additionally, the enrichment results represent only a fragmented portion of the deregulated gene list, as the interpretation is based solely on the biological functions and the genes participating in their enrichment. Consequently, relying solely on ORA, as done in the standard approach, is not sufficient. Additional methods are needed in combination to address the shortcomings of the standard approach.

Our main goal was to create a workflow that allows to include a greater part of the deregulated gene list in the interpretation, capturing a more complete picture of the transcriptomic data. We also aimed to improve functional enrichment detection sensitivity to identify more specific and smaller biological functions while minimising redundancy. Furthermore, we wanted to ensure the workflow is applicable to a wide range of both model and non-model organisms.

To meet these goals, we constructed Cluefish (Clustering, Enrichment and Fishing), a semi-automated workflow designed for comprehensive and untargeted exploration of transcriptomic data series. We developed Cluefish using a DR transcriptomic dataset from zebrafish embryos exposed to varying doses of dibutyl phthalate (DBP), a plasticiser found in human biological samples (34) and environmental surface waters (35). This main dataset allowed us to fully test and apply the workflow, conducting an in-depth biological interpretation. To demonstrate Cluefish’s versatility across different organisms with varying annotation richness in knowledgebases, we extended our validation to two secondary externally published DR transcriptomic datasets: Sprague Dawley rat (*Rattus norvegicus*) liver exposed to a dose-gradient of perfluorooctanoic acid (PFOA) (36), and Poplar (*Populus canadensis*) roots exposed to a dose-gradient of phenanthrene (PHE) (37). Cluefish is open-source and available under the CeCILL license, offering a powerful tool for improving biological interpretation and aiding in the development of targeted interventions.

## MATERIAL AND METHODS

### Zebrafish dataset

#### Fish

Zebrafish (*Danio rerio*) embryos (AB strain) were obtained from the Luxembourg Centre for Systems Biomedicine (LCSB) aquatic platform (Luxembourg University, Belval, Luxembourg). They were collected within 1 hour after spawn, placed in standardised fish water at 26°C (NF EN ISO 7346-2 (1998), pre-oxygenated by bubbling for 12h) and transferred to the LIEC laboratory facilities (Metz, France).

#### DBP solutions preparation and DBP quantification

Commercial DBP (purity 99%, CAS no. 84-74-2) was purchased from Sigma-Aldrich (St. Louis, MO, USA). Solvent-free DBP stock solution was made in fish water at an effective concentration of 5.4 mg/L. Stock solution and fish water effective DBP concentration quantification were performed 1 day before the beginning of exposure (3 technical replicates) on an Ultimate 3000 Rs chromatography system with UV detection at 230 nm (DAD) (Dionex, CA, USA).

Quantification was performed on an injected volume of 10 µL, using an Acquity UPLC HSS T3 2.5 µm, 2.1×100 mm column (Waters, Germany) as stationary phase, and 60% acetonitrile and 40% H2O as mobile phase, under a flow of 8 ml/min at 25°C. Fish water DBP quantification were below the limit of detection (LOD) of 3 µg/L. The stock solutions were stored in the dark at 26°C for the all experiment and were stable for at least the duration of exposure under these conditions (Supplementary Table S1). Tests solutions of DBP were made daily in fish water by serial dilution of the DBP stock solution in oxygenated fish water during the whole experiment to reach a concentration gradient [0 – 5 – 10 – 50 – 100 µg/L] built from environmental concentrations (median to high) up to the NOEC of DBP for chronic exposure in fish (100 µg/L) according to the European CHemicals Agency (ECHA). To avoid any external contamination of test solutions by phthalates or other plasticisers, only glassware was used for fish water, stock and test DBP solutions preparation, handling and storage.

#### Zebrafish embryos exposure and sample collection

30 zebrafish embryos (256 cells stage (38)) were randomly distributed in glass beakers with 60 mL of test solution, with 5 replicate beakers of 30 embryos for each exposure condition. Embryos were kept in a climate-controlled chamber (Cooled incubator MIR-554-PE, PHC Europe B.V.) at 26 (±0.5) °C, under 14:10 light/dark photoperiod (20 µE/m²/s). Exposure was semi-dynamic, with 1/3 of exposure solution volume being replaced every day by freshly prepared test solutions. After 120h exposure to DBP, pools of 10 embryos (‘protruding mouth’ stage (38)) were collected as one sample, flash-frozen in liquid nitrogen, and stored at −80 °C before RNA extraction. Remaining embryos were fixed in 4% paraformaldehyde (PFA) in PBS at 4 °C for 1 week and washed several times with PBS before morphological assessment.

#### Transcriptome sequencing

Total RNA of zebrafish embryos (pools of 10 embryos, 5 replicates per condition) was extracted using the RNeasy mini extraction kit (Qiagen). After quality and integrity assessment of extracted RNA by NanoDrop (Thermo Fisher Scientific, Waltham, MA) and by the RNA Nano 6000 Assay Kit of the Bioanalyzer 2100 system (Agilent Technologies, CA, USA), the 3 higher quality RNA samples per condition (A260/A280 > 1.7, A260/A230 > 2, RIN >8.8) were selected for high throughput mRNA sequencing. mRNA libraries were prepared after poly-A enrichment from 250 ng total RNA using NEBNext® Poly(A) mRNA Magnetic Isolation Module (New England Biolabs) followed by NEBNext® Ultra^TM^II Directional RNA library Prep kit (New England Biolabs). Library size was checked by the DNA 1000 Assay Kit of the Bioanalyzer 2100 system.

Libraries were sequenced on a NextSeq500 Illumina sequencer with 1x 75 bp single reads, providing 40 to 57 million reads for each sample. Library preparation, quality check and sequencing were performed by Helixio, France.

#### mRNAseq data processing

Raw mRNAseq data processing was performed on the Galaxy french server provided by the ‘Institut Français de Bioinformatique’ (IFB). Raw reads were filtered according to size (>55 bp) and quality parameters (mean Phred score >20 on 5’ and 3’ bases and on a 4 bp sliding window) using Trimmomatic (39), leading to keep 87 to 92% of total reads according to the sample. Cleaned reads were mapped to the zebrafish reference transcriptome GCRz11 and quantified using Salmon quant (v. 1.5.1) (40). Each transcript was annotated with the Ensembl transcript stable identifier (ID; e.g. ENSDART00000121728.3).

#### Embryos morphological traits

Lateral images of the fixed embryos were taken using a digital microscope (VMX-7000, KEYENCE, France). Body length and left eye area were measured using the “VHX-6000_950F Measurement Data Tabulation Tool” image processing software (KEYENCE, France).

### External datasets

To further evaluate the generalisability of Cluefish, datasets of two external transcriptomic studies were included. These studies were selected based on the inclusion of dose-response experimental designs and the contrasting richness of organism gene annotation available in public knowledgebases between them.

From Gwinn *et al.* (36), we selected the dataset exploring male Sprague Dawley rat (*Rattus norvegicus*) liver tissue subjected to perfluorooctanoic acid (PFOA), as their analysis reported a high number of deregulated transcripts and many enriched biological functions. The experiment involved exposing male rats, aged 8-10 weeks, to a concentration gradient of PFOA [0 – 0.156 – 0.3125 – 0.625 – 1.25 – 2.5 – 5 – 10 – 20 mg/kg] for up to 5 days. Each condition included four biological replicates. RNA was extracted from liver tissue and analysed via TempO-Seq. As described in the paper (36), filtered reads were aligned to the rat S1500+ probe sequences. Each transcript was labelled with a unique combination of NCBI gene symbol and probe IDs specific to TempO-Seq assays (e.g. NFIL3_9304). The count matrix was retrieved from the Gene Expression Omnibus (GEO) under the accession number GSE147072.

From Gréau *et al.* 2024 (37), we selected the dataset exploring root tissue exposed to phenanthrene (PHE) as their analysis reported the highest number of deregulated transcripts. Poplar plants (*Populus canadensis)* were exposed for four weeks to a phenanthrene (PHE) concentration gradient [0 – 100 – 200 – 400 – 700 – 1000 – 1500 – 2000 mg kg^-1^]. Each condition included eight biological replicates. RNA was extracted from root tissue and analysed via RNAseq. As described in the paper (37), filtered reads were mapped and counted on the *Populus trichocarpa* genome (V4, Phytozome). Each transcript was annotated with the Ensemble gene stable ID (e.g. Potri.001G000400). The count matrix was retrieved from GEO under the accession number GSE263776.

The three datasets, the main dataset and two external ones, will hereafter be referred to as the zebrafish, rat liver, and poplar root datasets.

### Dose-response analysis

The DRomics R package was used to identify significantly deregulated transcripts and model Dose-Response (DR) curves. The default parameters in DRomics were applied to all three datasets (41). Transcript count datasets were imported, normalised, and transformed into a logarithmic scale using the rlog method within DRomics (42).

Significantly deregulated transcripts were selected using quadratic trend tests with a False Discovery Rate (FDR) of 0.05 (43). DR models (linear, Hill, exponential, Gauss-probit, or log-Gauss-probit) were fitted using non-linear regression and selected based on the lowest second-order Akaike information criterion (44). The modelled DR curves were characterised by their trends (bell-shaped, decreasing, increasing, and U-shaped) and yielded a benchmark dose (BMD-1SD) (22), calculated from a benchmark response (BMR-1SD) using the following equation: *BMR* 1 *SD*= *y* 0 *± SD*, where y0 is the theoretical level at the control given by the DR model and SD is the residual standard deviation of the modelled DR curve (22, 45). The 95% confidence intervals on the BMD-1SD values were computed by non-parametric bootstrap (5000 iterations). Modelled DR curves were discarded if the point or the interval estimates of the BMD could not be successfully calculated. For the zebrafish embryo morphological traits, body length and left eye area were likewise analysed using the DRomics workflow for continuous anchoring (individual) data.

### The Cluefish workflow

Cluefish is composed of 11 main data analysis steps, followed by optional data visualisation steps (Figure 1). The workflow was applied to the in-house zebrafish dataset and the two external published datasets. Default Cluefish parameters and detailed steps for the zebrafish case study are described below, whereas details on the choices made for the two external datasets are provided in the Supplementary Text S1. Each step is as follows:

**Figure 1.**
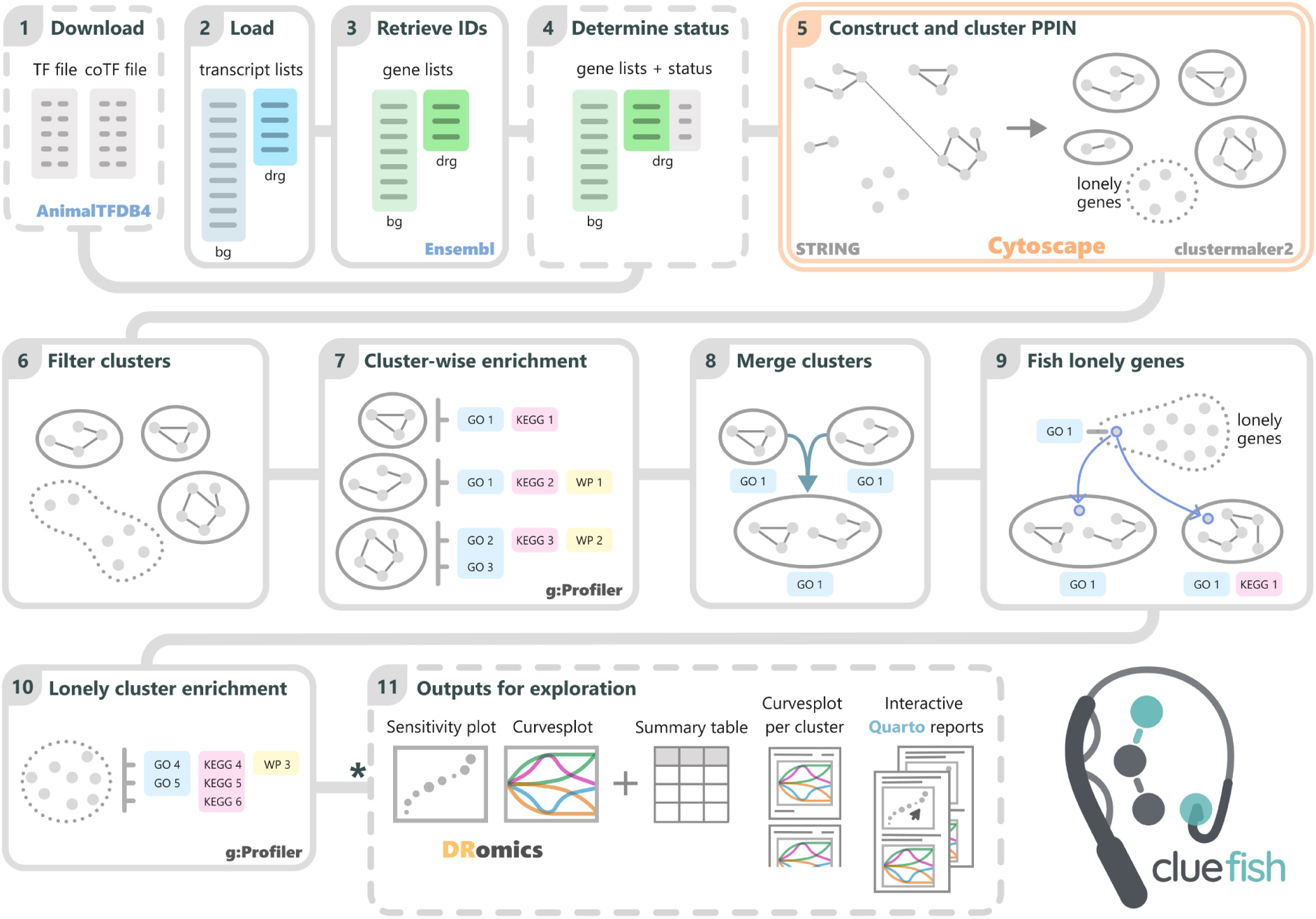
Schematic of Cluefish workflow. Steps performed in R are shown in single-line frames, with dashed frames for optional steps, while the step carried out in Cytoscape is shown within a double lined frame. In step 5, deregulated genes, which code for proteins in STRING, are represented as nodes. Their interactions are illustrated by lines. Clusters are represented by full circles, while isolated genes are categorised as *lonely genes*, forming the *lonely cluster*, represented by a dashed circle. The schematic guides the user through the various steps but does not explicitly depict the output-to-input relationships, as some steps require multiple previous outputs. The full detailed workflow is provided in the vignette (https://ellfran-7.github.io/cluefish/articles/cluefish.html) \**The final step can only be performed if the list of deregulated transcripts loaded in Step 2 is derived from the outputs of DRomics*. Abbreviations: bg: background, drg: deregulated.

#### Step 1: Download transcription (co-)factor annotations

Transcription factors (TFs) and transcriptional co-factors (CoTFs) form multi-protein complexes that mediate cellular signalling and facilitate gene transcription (46, 47). This combinatorial function establishes a crucial link between signalling pathways and gene regulation (48). To incorporate this regulatory information, lists of TF and CoTF genes for the specified organism are obtained from the Animal Transcription Factor Database 4.0 (AnimalTFDB4) (49). If these files are not available for a particular organism, this step can be skipped. The zebrafish TF and CoTF gene lists were successfully downloaded.

#### Step 2: Load background and deregulated transcript lists

The background transcript list includes all transcript IDs detected in the experiment, whereas the deregulated transcript list contains only the IDs of significantly deregulated transcripts. The deregulated transcript list was derived from the DRomics pipeline in our case studies.

#### Step 3: Retrieve gene identifiers

Transcript IDs are converted to gene IDs and other pertinent annotations by linking with the Ensembl database via the biomaRt R package (50, 51). To ensure compatibility with databases and tools used later in Cluefish, specifically the STRING database and g:Profiler (52), the results must include IDs supporting the organism in both platforms. In most cases, the Ensembl gene ID meets this requirement for well-referenced organisms, significantly reducing the need for repeated ID conversions throughout the workflow. In this step, all detected transcript IDs are mapped to their corresponding Ensembl gene IDs, forming the foundation for the deregulated gene list. In our case, the zebrafish transcript IDs were converted to their Ensembl stable gene ID and external gene name, corresponding to readable gene names.

#### Step 4: Determine deregulated gene regulatory status

The next step involves determining the regulatory status of the deregulated genes, using the TF and CoTF annotation files acquired earlier. If these files are not available for a particular organism, this step can be skipped. Since the files were successfully downloaded for zebrafish, it was possible to assess the regulatory status of the deregulated genes.

#### Step 5: Construct and retrieve the clustered protein-protein interaction (PPI) network

The preceding output is then manually imported into the Cytoscape application (53). Using the StringApp plugin (54), the deregulated transcripts’ STRING-compatible IDs are entered into the Protein Query search bar. This query process is customisable, offering inclusion of specific interaction types, setting limits for interaction confidence scores, and supplementary interactions. The default value in STRING is set at 0.4 for the confidence score limit to construct the PPI network, representing the probability that the interaction really exists given the available evidence. In our zebrafish case study, a confidence score of 0.9 was utilised to only consider highly confident interactions, and we included all interaction types. The resulting PPI network was constructed from both known and predicted protein-protein associations from the STRING database. To identify clusters, the Markov Chain Clustering (MCL) algorithm is applied using the clustermaker2 extension (55, 56). The granularity of clusters is controlled by the inflation parameter, with lower values (1–2) yielding fewer, larger clusters, and higher values (5–6) resulting in more, smaller clusters. Very large clusters may lack coherent biological context, while very small clusters may not represent a biological complex or may only be involved in specific cases. In our zebrafish case study, this parameter was set at 4 which is the default value in clustermaker2. This setting balances the creation of few large clusters and many small clusters. The node table, containing clustered elements, is manually exported as a CSV file.

#### Step 6: Filter clusters

Returning to the R environment, clusters are filtered based on their gene set sizes using what is called the lower cluster size filter, under which clusters are removed. By default, in Cluefish, this filter is set at 4. The choice of lower cluster size limit may depend on the study, organism, and earlier decisions during PPI network creation and clustering. Higher size limits result in fewer large clusters, whereas smaller limits retain both large and small clusters, removing only very small clusters. In our zebrafish case study, the default value was used.

#### Step 7: Perform cluster-wise functional enrichment

To characterise each cluster, ORA can be conducted using the gprofiler2 R package (57), which interfaces with g:Profiler. This tool was chosen firstly for its extensive support of both model and non-model organisms, totaling 984 and the ability to use custom annotation data. The main tool of interest in g:Profiler is g:GOst, which can perform ORA on multiple gene lists and search against a wide range of functional databases, including GO, KEGG and Wikipathways (WP). Specifically, Fisher’s one-tailed tests can be applied to measure the likelihood that the overlap between a query gene set and biological function gene sets is due to chance. Another important aspect of g:Profiler is its access to previous versions of the software and data, ensuring reproducible analyses. Furthermore, it offers a filtering method for highlighting GO driver terms by considering the underlying topology of annotations, as described in Kolberg et al. 2023 (52). By default, in Cluefish, the deregulated gene list is concurrently analysed against GO Biological Processes (GO:BP), KEGG and WP pathways, using the background gene list comprising all annotated protein-coding genes from the experiment. The FDR multiple testing correction method is applied, with a significance limit set at 0.05. To address the inherent complexity and redundancy within GO term hierarchies, only highlighted GO:BP terms are retained. In our zebrafish case study, the default settings were chosen. In addition to these, we implemented two new filters.

The first one, called the biological function size filter, are the minimum and maximum gene set sizes of biological functions, i.e., the number of genes associated with each biological function found within the background gene list. By default, in Cluefish, lower and upper biological function size limits are set at 5 and 500 respectively. Very large gene sets, which are dependent on the organism, are often associated with broad biological processes. These can introduce noise and obscure more specific functions. Conversely, very small gene sets hold little statistical support, as minor changs in gene membership can substantially affect FDR values, leading to unstable enrichment. Additionally, very small gene sets can be overly specific and may not capture the full biological context of a pathway. These lower and upper limits can be adjusted based on the study, the organism, and the type of transcriptomic data (e.g., whole transcriptome or tissue-specific). In our zebrafish case study, the default values were used.

The second filter, called the enrichment gene count filter, is the minimum number of genes participating in the enrichment of a biological function. By default, in Cluefish, the enrichment gene count filter is set at 3 in order to avoid identifying biological functions enriched by too few genes, particularly in smaller clusters. Such clusters can lead to a higher rate of false positives, and enrichments based on very few genes may not provide substantial biological insight into the cluster itself. This limit should be proportionate to the cluster sizes, particularly the smallest clusters. In our zebrafish case study, the default value was used.

#### Step 8: Merge clusters

After characterising the clusters, the next step is to identify and merge clusters that share the same enriched biological functions from at least one specified database (e.g. GO, KEGG etc.) in the ORA. By default, in Cluefish, the merging process is conducted separately for each chosen database, in descending order of the total number of unique genes present in both the database and the background list.

This step combines clusters whose proteins were insufficiently interactive in STRING to form a single cluster but are shown through functional enrichment to participate in the same biological processes. Merged clusters thus form larger, contextually unified groups representing distinct biological functions. In our zebrafish case study, the default settings were chosen, with clusters first merged based on shared GO Biological Process (GO:BP) functions first, followed by KEGG, and finally WP.

#### Step 9: Fish lonely genes

Genes not assigned to any cluster are referred to as « *lonely genes* ». These genes are not typically explored because they do not contribute to functional enrichment, and thus, they do not help in characterising the data. Genes become *lonely* in one of two cases: either they are initially not sufficiently interactive with another protein complex to participate in or form a cluster, or they were initially part of a cluster but failed to pass the cluster size filtering step, thus falling into the *lonely cluster*. To achieve a more comprehensive analysis and expand clusters, *lonely genes* are incorporated into existing clusters based on shared functional annotations. Specifically, if a biological function annotation of a *lonely gene* matches a biological function enriched in a cluster, the *lonely gene* is integrated into the corresponding cluster. The *friendliness* metric represents the number of clusters a *lonely gene* can be incorporated into.

#### Step 10: Perform functional enrichment of the lonely cluster genes

The *lonely cluster*, now consisting of the remaining *lonely genes*, can undergo a simple functional enrichment analysis to gain further insight into the biological context of the genes not associated to any existing clusters. The customisations used in the earlier cluster-wise enrichment analysis are fully applicable here as well. Therefore, the parameters were set to match those chosen in the previous cluster-wise functional enrichment analysis for the zebrafish dataset.

#### Step 11: Generate outputs for exploration

To fully explore the Cluefish results, DRomics results are required in order to include Dose Response (DR) modelling metrics. While DR analysis is not required for previous steps of Cluefish, it becomes essential for generating the following outputs. A summary table, which captures key details from each step and tool used in the Cluefish workflow, can be generated and exported to CSV for manual exploration. Additionally, plots of fitted DR curves for each cluster’s gene set can be generated and exported to PDF, offering a visual exploration of cluster content. Three report templates created using Quarto (Allaire, J.J., Teague, C., Scheidegger, C., Xie, Y., & Dervieux, C., 2024. Quarto. Zenodo. https://doi.org/10.5281/zenodo.5960048) are incorporated into the Cluefish compendium, which may be useful to users. Examples can be found in Supplementary Files S3, S4 and S5. These templates summarise Cluefish results interactively, integrate them with DRomics findings, and enhance data usability for comprehensive biological interpretation. Although these reports can be generated automatically, they may require adjustments to suit various types of cases.

### The standard workflow

To illustrate Cluefish’s capacity to improve biological interpretation of DR transcriptomic data, we compared it with the standard approach. This involved applying a simple functional enrichment analysis to the entire deregulated gene list, using the same parameters as those chosen for the cluster-wise enrichment step in Cluefish. For the zebrafish and rat liver datasets, three databases were used: GO:BP, KEGG and WP. For the poplar root dataset, only GO:BP and KEGG were used, as WP lacked sufficient coverage. Both approaches were compared in terms of number of enriched biological functions and number of deregulated genes involved in them.

### Implementation

For gene ID retrieval and functional enrichment analyses across all three datasets, we used Ensembl genome assemblies (release 111): GRCz11 (GCA_000002035.4) for zebrafish, mRatBN7.2 (GCA_015227675.2) for rat liver, and Pop_tri_v4 (GCA_000002775.4) for poplar root. Details of the versions of tools and databases used for each dataset are provided in Supplementary Table S2. Cluefish is primarily implemented in the R statistical computing language (58), with a short manual processing step halfway in the Cytoscape application. Dependencies in R, including binary dependencies, are automatically bundled and installed using renv (v. 1.0.7) (59). Cluefish is structured as a research compendium, which is an organised collection of files and directories encompassing all necessary data, code, and documentation required to reproduce the research (60). Git is utilised for version control (v. 2.45.2.windows.1.). The workflow requires two main data inputs: the background and deregulated transcript lists. If these inputs come from DRomics results, the entire Cluefish workflow can be used. However, if the inputs are from a different selection method, Cluefish can be applied up to Step 10. This is because the outputs for visual exploration depend on DRomics figures (e.g., DR curves) and the associated computed metrics. Cluefish is freely available at https://github.com/ellfran-7/cluefish with an associated vignette serving as a guide through the Cluefish workflow with some additional functionalities available, which has not been included in the workflow description for the sake of simplicity (https://ellfran-7.github.io/cluefish/articles/cluefish.html).

## RESULTS

### Dose-response analysis

A total of 41396, 2654, and 34699 transcripts were detected in at least one sample across the zebrafish, rat liver, and poplar root datasets, respectively. One control sample from the zebrafish dataset was excluded from the analysis due to its divergence in the principal component analysis (PCA) plot, indicating it was an outlier (Supplementary Figure S1). Thus, the analysis proceeded with the remaining 14 samples from the original 15, resulting in 39890 transcripts forming the background transcript list for the zebrafish dataset. The main outputs of the dose-response analysis with DRomics are summarised in Table 1 and Table 2 for all three datasets. Across all datasets, most deregulated features exhibited monotonic trends along the dose-gradient, with 93%, 79%, and 60% showing either increasing or decreasing trends for the zebrafish, rat liver, and poplar root datasets, respectively, and a minority of bell or U-shaped trends.

**Table 1.** Number of transcripts across the three datasets (zebrafish, rat liver, and poplar root) showing the total experimental background, deregulated transcripts identified by DRomics, and deregulated transcripts with computed benchmark dose (BMD) and confidence intervals (CI).

|  | Zebrafish | Rat liver | Poplar root |
| --- | --- | --- | --- |
| Total background transcripts | 39890 | 2654 | 34699 |
| Deregulated transcripts | 2449 | 873 | 6463 |
| With BMD and CI | 2433 | 843 | 6352 |

**Table 2.** Distribution of transcript trend types across the three datasets (zebrafish, rat liver, and poplar root). The table presents the number and percentage of transcripts exhibiting monotonic (increasing or decreasing) and biphasic (bell-shaped or U-shaped) trends among deregulated transcripts with computed benchmark dose (BMD) and confidence intervals (CI).

|  | Zebrafish | Rat liver | Poplar root |
| --- | --- | --- | --- |
| Increasing (inc) | 1132 (46%) | 314 (37%) | 2156 (34%) |
| Decreasing (dec) | 1145 (47%) | 353 (42%) | 1683 (26%) |
| Bell-shaped (bell) | 64 (3%) | 71 (8%) | 1375 (22%) |
| U-shaped (U) | 92 (4%) | 105 (13%) | 1138 (18%) |

### Comparison of both approaches

Before performing functional enrichment analysis, transcript IDs were converted to Ensembl gene IDs for the zebrafish and rat liver datasets, as described in Step 3. The poplar root dataset was already annotated with Ensembl gene IDs. In the zebrafish dataset, the background and deregulated transcript lists (with computed BMD and CI) were converted to 27729 and 2365 unique Ensembl stable gene IDs, respectively. For the rat liver dataset, these lists were mapped to 2367 and 756 unique Ensembl stable gene IDs, respectively.

Simple functional enrichment analysis revealed that the deregulated gene lists significantly enriched multiple biological functions across the GO:BP and KEGG databases in all datasets, and in the WikiPathways (WP) database for the zebrafish and rat liver datasets only.

For the zebrafish dataset, the standard approach identified enrichement in 12 GO:BP, 6 KEGG and two WP biological functions. A total of 320 genes contributed to these enrichments, while 2045 genes did not (Supplementary Table S3). Cluefish identified all 6 KEGG and 2 WP pathways found through the standard approach and additionally detected 45 KEGG and 8 WP pathways (Figure 2A). In contrast, only 5 GO:BP terms were shared between both approaches. Cluefish identified 36 unique GO:BP terms, whereas the standard approach identified seven. These seven GO:BP functions were higher-level parent terms, representing more general and broader biological concepts. They were redundant with the lower-level child terms found by Cluefish, although they did not share the same GO term names. Cluefish led to the consideration of an additional 733 genes into enriched biological functions, accounting for 30% of the deregulated gene list (Figure 2B). Of the 320 genes found by the standard approach, 293 were also recovered by Cluefish, leaving 27 unique to the standard approach.

**Figure 2.**
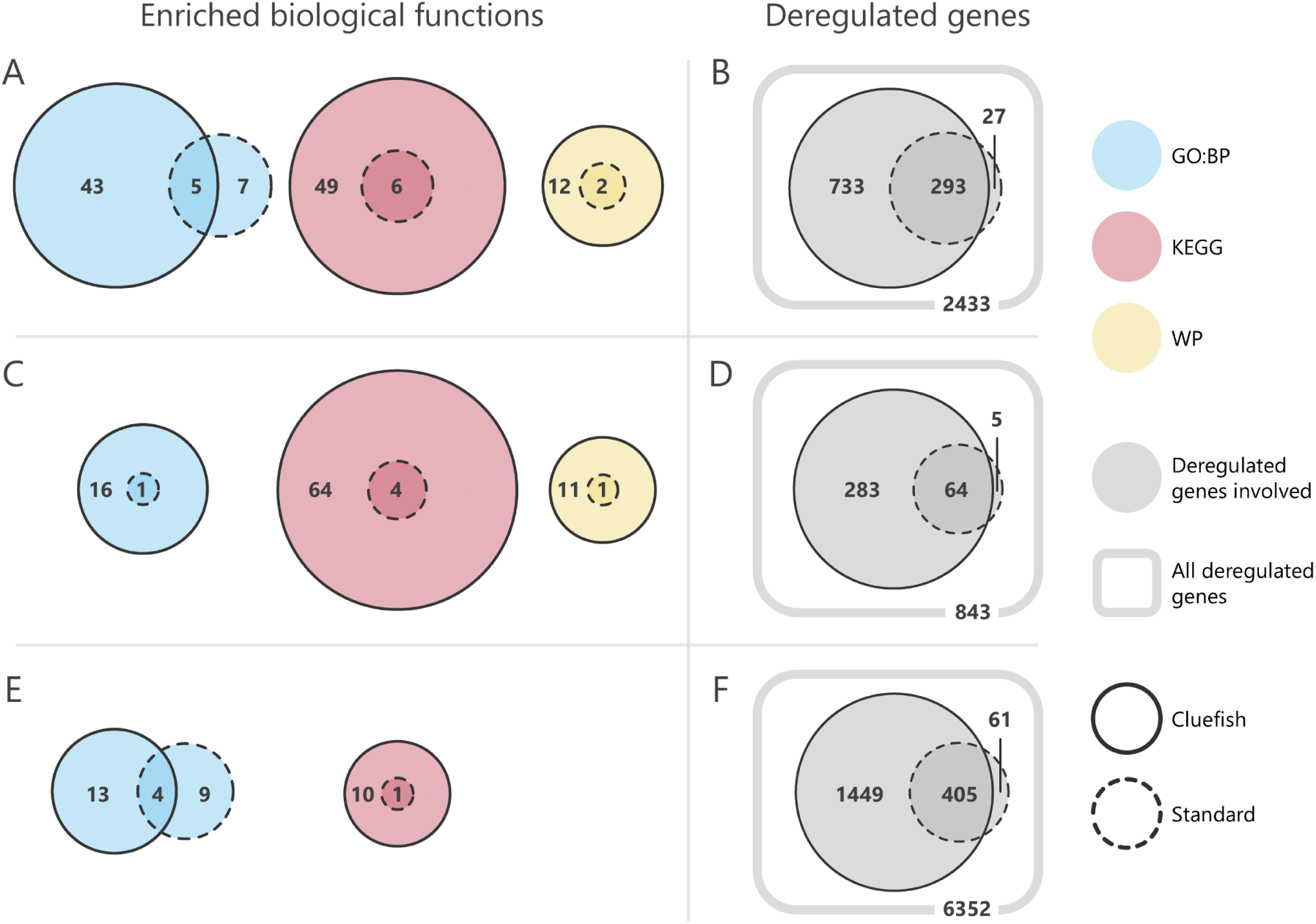
Significantly enriched biological functions (left panel) and involved deregulated genes (right panel) for zebrafish (A, B), rat liver (C, D), and poplar root (E, F) datasets. Left panels (A, C, E) illustrate the overlap of enriched functions by database and approach. Right panels (B, D, F) display the overlap of deregulated genes involved in these enriched functions, representing the set of genes considered for biological interpretation for both approaches. Rounded squares indicate the total number of deregulated genes with computed benchmark dose (BMD) and confidence intervals (CI)).

For the rat liver dataset, the standard approach identified enrichment in one GO:BP, four KEGG, and one WP biological function, involving 69 genes, whereas 687 genes did not participate in any of these enrichments (Supplementary Table S4). Cluefish recovered all functions detected through the standard approach, while also adding 16 GO:BP, 64 KEGG and seven WP functions (Figure 2C). Cluefish incorporated an additional 282 genes into enriched biological functions, accounting for 37% of the deregulated gene list (Figure 2D). Of the 69 genes participating in enrichment found by the standard approach, 64 were also recovered by Cluefish, leaving 5 genes unique to the standard approach. Full details on outputs and details of key steps throughout the Cluefish application of the rat liver dataset can be found in Supplementary Tables S5 to S10.

For the poplar root dataset, the standard approach identified enrichment in 13 GO:BP and 1 KEGG pathway, involving 466 genes, while 5886 were not associated with any enriched function (Supplementary Table S11). Cluefish recovered the same KEGG pathways detected by the standard approach and identified 9 additional ones (Figure 2E). Only 4 GO:BP terms were shared between both approaches. Cluefish identified 13 unique GO:BP terms, whereas the standard approach identified 9 unique GO:BP terms. Of the 9 GO:BP terms uniquely identified by the standard approach, some were redundant with terms found by Cluefish, represented under different names and at different levels of the GO hierarchy. For example, while both approaches identified the “translation” (GO:0006412) term, the standard approach additionally identified the “Ribosome” (KEGG:03010) and “ribosome assembly” (GO:0042255) term. These biological terms substantially overlapped with “translation”. Specifically, 93 out of 101 deregulated genes (92%) annotated for “Ribosome” and 26 out of 31 deregulated genes (84%) annotated for “ribosome assembly” were also among the 198 genes annotated for “translation” (Supplementary Table S12). Cluefish led to the consideration of an additional 1778 genes into enriched functions, accounting for 28% of the deregulated gene list (Figure 2F). Of the 466 genes considered by the standard approach, 403 were also considered by Cluefish, leaving 63 genes uniquely considered by the standard approach. Full details on outputs and aspects of key steps throughout the Cluefish application of the poplar root dataset can be found in Supplementary Tables S5, S6, S13, S14, S15 and S16.

### Exploration of the Cluefish application to the zebrafish dataset

#### Overview of the results

From the AnimalTFDB4 database, 3328 Ensembl genes corresponding to known zebrafish TFs and CoTFs were retrieved. Among the 2365 deregulated genes, 286 were identified as coding for these transcriptional regulators. After constructing the PPI network and performing MCL clustering, 204 clusters were formed (Supplementary Table S5). Of these, only 53 clusters passed the size filter, encompassing 399 genes. Cluster-wise ORA identified 41 driver GO:BP, 51 KEGG and 10 WP enriched functions. See the Supplementary Table S6 for the cluster-wise ORA results both before and after filter, and the Supplementary Table S17 for the initial ORA results prior to filtering. Several clusters were merged based on shared biological function, reducing the total number of clusters to 44. Following *lonely fishing*, the number of genes in the newly expanded clusters increased to 904 (37% of the deregulated gene list). The majority of fished *lonely genes* had a *friendliness* value of one, two, or three, with the highest value reaching eight. The final number of clusters was 45 (44 expanded clusters + *Lonely cluster*) (Supplementary Table S18). The size of the *lonely cluster* was greatly reduced, containing 1461 genes. The functional enrichment of the *lonely cluster* identified 6 driver GO:BP, 4 KEGG and 2 WP biological functions (Supplementary Table S19). These functions did not overlap with any from the cluster-wise enrichment. With this added information, over 1026 deregulated genes enriched at-least one biological function, corresponding to 43% of the deregulated gene list. The sensitivity of the 44 expanded clusters can be illustrated using summarised BMD-1SD values for the transcripts within each cluster (Figure 3A). The first quartile represents the value below which 25% of the BMD of the transcripts fall below. According to this summary, the most sensitive clusters #43, #37, #10 and #15 with BMD-1SD first quantile values of approximately 1.95, 2.19, 2.23 and 2.62 µg/L, respectively. In contrast, clusters #51, #27, #35 and #38 were the least sensitive, with first quartile values of 33.6, 31.7, 31.2 and 29.8 µg/L, respectively (Supplementary Table S20).

**Figure 3.**
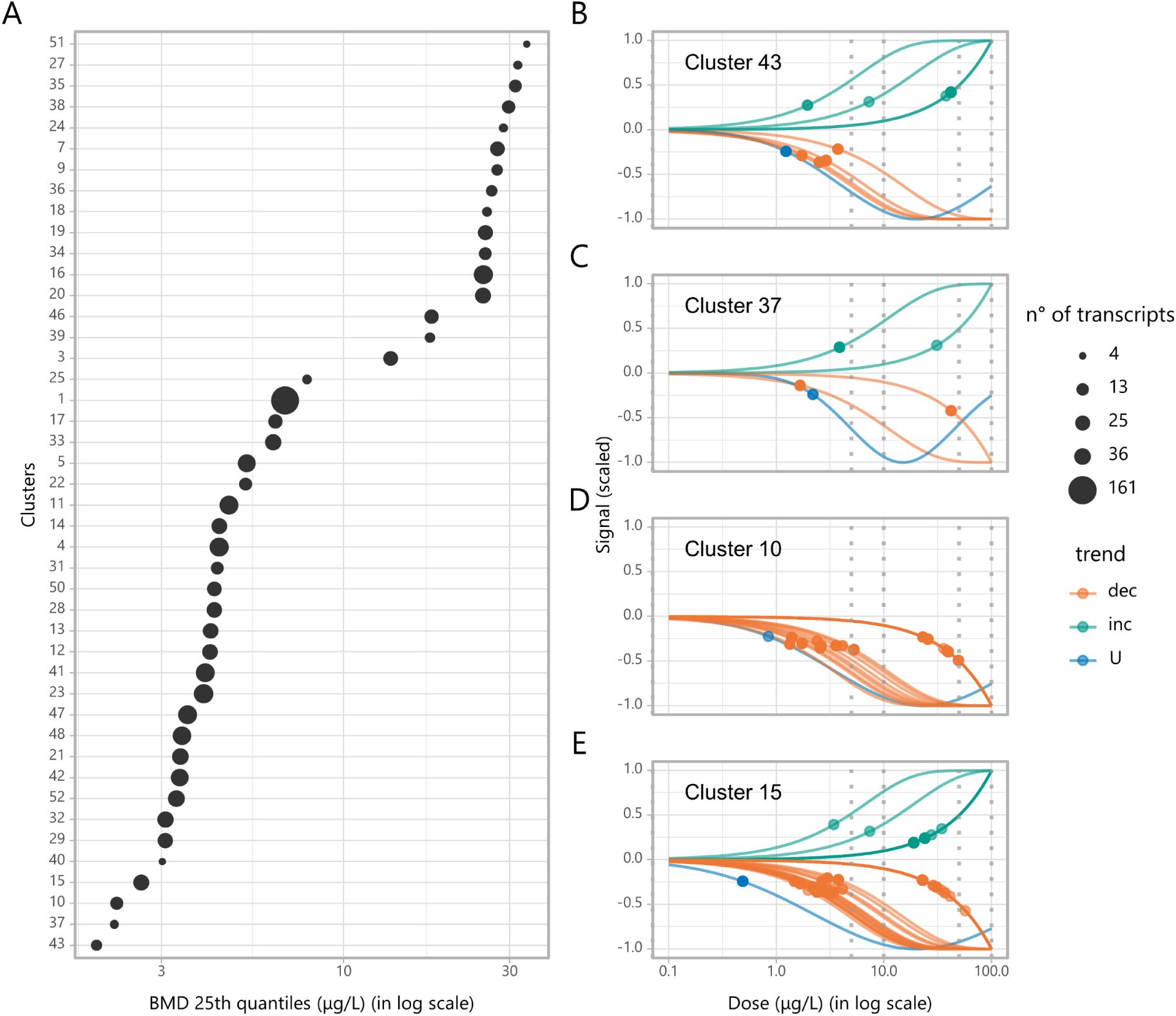
Results of the Cluefish workflow applied to the zebrafish dataset. (A) Summarised BMD 25th quantile values for each cluster, with points ordered from most to least sensitive and sized according to the number of transcripts. (B, C, D, E) Dose-response fitted curves and computed BMD points for transcripts in (B) cluster 43, (C) cluster 37, (D) cluster 10 and (E) cluster 15, with colors indicating the trend for each curve shape: dec (decreasing), inc (increasing), and U (U-shaped). Grey dotted vertical lines represent the experimental doses.

#### In-depth exploration of findings

In ecotoxicology, the primary focus is on examining the impact of low doses of contaminants in the environment. Prioritising the exploration of the most sensitive clusters is a reasonable approach in this context.

Since the cluster #37 was not characterised, it was excluded from subsequent analysis (Figure 3C). We chose to focus on clusters #43, #10, and #15, as they exhibited the lowest 25th BMD quantile values. Cluster #43 comprised 9 distinct transcripts and genes, with some showing increasing and some showing decreasing trends (Figure 3B). Most of the transcripts BMDs ranged between 1.23 and 7.27 µg/L. This cluster solely enriched the “Retinol metabolism” (KEGG:00830) KEGG pathway, with no GO:BP or WP function enrichments. Cluster #10 was composed of 16 distinct transcripts, with the majority showing decreasing trends, also with similar shaped DR curves (Figure 3D). Ten BMD values ranged between 0.84 and 5.29 µg/L, while the remaining six ranged between 23.1 and 49.4 µg/L. This cluster enriched sphingolipid metabolism functions in GO:BP and KEGG, specifically “sphingolipid metabolic process” (GO:0006665) and “Sphingolipid metabolism” (KEGG:00600), respectively. Cluster #15 comprised 31 distinct transcripts, mostly showing decreasing trends, also with similar DR curve shapes (Figure 3E). In this cluster, 18 BMD values ranged between 0.49 and 7.37 µg/L, while the remaining 13 BMD values ranged between 19.0 and 57.1 µg/L. This cluster enriched one GO:BP (lipid biosynthetic process - GO:0008610), one KEGG (Steroid biosynthesis - KEGG:00100), and one WP (Cholesterol biosynthesis - WP:WP1387) biological function. None of the 3 clusters contained known transcriptional regulators.

#### Investigations towards biological interpretation

Discovering the Retinol metabolism pathway as not only deregulated but also highly sensitive prompted us to investigate the involvement of Hox genes, which are among the primary canonical targets of retinoic acids (61–63). All the Hox genes were located within the *lonely cluster*, where we identified 13 Hox gene transcripts, all showing increasing trends (Figure 4A). Eleven of these transcripts had BMD values between 2.51 and 6.80 µg/L and very similar DR curves, while the remaining two had values of 28.9 and 40.6 µg/L (Supplementary Table S21). All of these genes act as transcriptional regulators.

**Figure 4.**
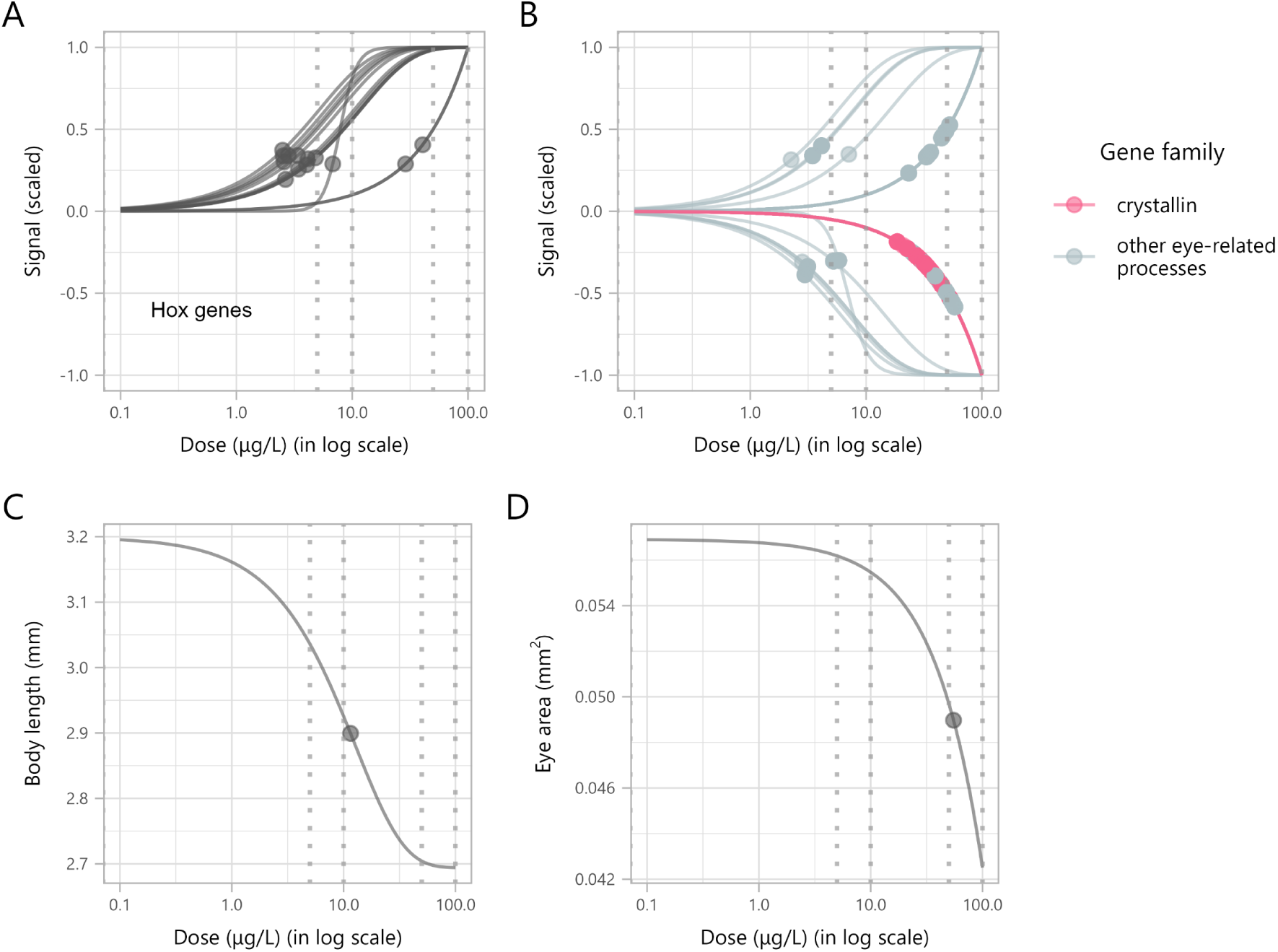
Dose-response curves and computed benchmark dose (BMD) points for deregulated zebrafish transcripts and individual endpoints. (A) Hox gene transcripts. (B) Transcripts associated with eye development and function, with transcripts’ genes from the crystallin family highlighted in pink and other eye-related genes in light grey. (C) Body length measurements. (D) Eye surface area measurements. Grey dotted vertical lines represent the experimental doses.

Further exploration of the *lonely cluster* revealed that the biological functions enriched by the highest number of deregulated genes were “lens development in camera-type eye” (GO:0002088) and “visual perception” (GO:0007601), each enriched by 26 genes. In total, 58 deregulated genes were related to eye development or function (Figure 4B).

Among these, 27 crystallin gene transcripts were found, all following decreasing trends, with BMD values ranging from 18.6 to 52.6 µg/L (Supplementary Table S22). Our transcriptomic findings led us to investigate individual endpoints. At the individual level, zebrafish embryos exhibited a dose-dependent reduction in body length and eye surface area, with BMD values of 11.5 and 55.1 µg/L, respectively (Figure 4C-D). See Supplementary Table S23 for the raw left eye area and body length measurements.

## DISCUSSION

Building the road towards the biological interpretation of transcriptomic data series is a challenging task. To address this, we developed Cluefish, a free and open-source, semi-automated R workflow that offers a comprehensive and untargeted exploration of such data. When used alongside DRomics, Cluefish offers a more complete analysis of dose-response (DR) transcriptomic data. This workflow aims to address the limitations of the standard approach towards biological interpretation by leveraging the power of ORA on pre-clustered networks. These clusters act as anchors for ORA, improving enrichment detection sensitivity. This allows us to identify smaller, more specific biological processes while simultaneously forming exploratory gene groups.

Cluefish integrates valuable information from various biological function/pathway databases (e.g., GO, KEGG, WP), the AnimalTFDB4 database of transcription factors and co-factors for 183 animal genomes, and the STRING database of known and predicted PPIs for 12535 organisms. Despite drawing from multiple complementary databases, these databases collectively provide a broad coverage of organisms, ensuring that Cluefish is applicable to a wide range of both model and non-model organisms. Innovative features such as cluster merging and the fishing of *lonely genes* based on shared biological context ensure a more comprehensive analysis.

The standard approach identified a limited, overly broad set of enriched functions for the three studied datasets. This limitation stems from the statistical method and the coarse granularity of functional gene sets. As a result, both the qualitative and quantitative interpretation of the data suffer. Additionally, focusing solely on enriched functions is risky, as it may overlook valuable insights from deregulated genes that do not contribute to functional enrichment. Indeed, with the standard approach, a substantial proportion of deregulated genes failed to contribute to any functional enrichment: over 84% in zebrafish, 81% in rat liver, and 93% in poplar root datasets. By analysing biologically homogeneous gene subsets, Cluefish minimises the influence of unrelated genes and ensures a more focused analysis. Moreover, by reintegrating *lonely genes* back into these clusters through shared biological context, Cluefish enhances the biological coherence of each cluster and increases the number of genes available for biological interpretation.

Some genes, however, remain lonely and persist in the lonely cluster. These genes should not be dismissed as noise, as their exclusion from PPI-based clusters likely reflects either (1) limited interactions or annotation data for the organism in public databases, or (2) their involvement in non-physical associations, such as transcriptional regulation or functional similarity not captured through physical interaction networks. By explicitly characterising the biological functions represented in the lonely cluster, Cluefish ensures that no potentially valuable data is overlooked.

The effectiveness of the Cluefish workflow is illustrated in our zebrafish case study by the identification of specific clusters, such as clusters #43, #10, and #15, which contain transcripts with high sensitivity values. Importantly, the biological functions tied to these clusters, including retinol and lipid metabolism, were missed by the standard approach, as they were not identified as enriched. Cluster #43 includes genes involved in retinol (vitamine A) metabolism. Our findings suggest a mechanistic hypothesis centred on the disruption of retinoic acid (RA) signalling. Disrupted retinol metabolism may lead to altered levels of biologically active retinoids, such as 9-*cis* RA, all-trans RA (ATRA), and 11-*cis* Retinal, causing significant changes in retinoid signalling in zebrafish embryos. This hypothesis is supported by the deregulation of several ATRA target genes abundance. Hox genes, particularly those located in the anterior part of the embryo, are canonical targets of RA signalling (64, 65). Among the 48 Hox genes in the zebrafish genome (66), we detected 47 in our dataset, with 13 upregulated and homogenous in DR curve shape, all located in the anterior positions 1 to 9 of the Hox clusters. Other canonical ATRA target genes are *cyp26a1*, *cyp26b1* and *cyp26c1*. Cyp26 enzymes are key regulators of ATRA levels control in cells, by transforming ATRA into inactive metabolites. Cyp26 are thus responsible for the prevention of activation of RA receptors by RA in specific cells (65). Cyp26 coding transcripts are not identified as deregulated in our dataset (Supplementary Figure S2A-C). However, *cyp26a1* was selected as significantly deregulated along the dose-gradient but no DR model was successfully fitted to the DR curve of this transcript. Visual inspection of expression data for *cyp26a1* suggests however a bell-shape response of transcript abundance. Alltogether, these responses in abundance of transcripts of canonical ATRA response genes suggest an increased ATRA signalling at the lowest DBP exposure concentrations.

RA plays a central role in early patterning and definition of the spinal cord during zebrafish development (67). Disruption of this pathway has been linked to teratogenic effects, including spinal deformities such as bent spine and malformed tail, highlighting its crucial role in embryogenesis (68, 69). The dose-dependent reduction in body length observed in zebrafish embryos further strengthens this connection. In addition, the crucial role of RA in ocular development (70) can be linked to the identification of 59 deregulated genes related to eye development. The downregulation of crystallin genes —responsible for maintaining lens transparency and refractive power (71, 72) —points to potential defects in lens function. The dose-dependent reduction in eye surface area aligns with this, suggesting compromised eye development linked to disrupted retinoid signalling.

The sensitive downregulation of genes involved in lipid metabolism and transport (clusters #10 and #15) could also be attributed to the deregulation of retinol metabolism under DBP exposure. 9-*cis* RA serves as a ligand for the retinoid X receptors (RXR), that acts as co-receptors for several nuclear receptors involved in lipid metabolism genes regulation (73). This may otherwise be due to the sensitive downregulation of one of the genes coding for RXRs, *rxraa* (Supplementary Figure S3). While it is currently unclear whether rxr genes expression is under direct regulation of retinoids, both this *rxraa* downregulation at the lowest DBP concentration and retinoid abundance deregulation could directly or indirectly participate to the downregulation of lipid metabolism and transport genes.

This hypothesis of DBP mode of action based on retinoid signalling disruption is consistent with findings from previous studies (69, 74, 75). This has been previously suggested through screening in mouse stem cell lines (75) and observed in transcriptomic data from adult male fish testes (76) as well as in rodent models (77). However, most studies on DBP focus on its interaction with the PPAR signalling pathway (78–80). By applying the Cluefish workflow, our DR transcriptomic data suggests that retinoid signalling, and function disruption may be the most sensitive pathway affected by DBP during zebrafish development, potentially leading to significant morphological changes. RA signalling, while still understudied in the context of endocrine disruption, has been increasingly recognised as vulnerable to pollutants in both toxicological and ecotoxicological settings (81, 82). Given the critical role of RA signalling in embryonic development, which is conserved across both vertebrates and invertebrates, the potential impact of DBP on this pathway deserves closer attention (83). These findings illustrate how Cluefish can reveal connections that would have been missed by the standard approach. By investigating clusters, we identified links between them and guided us to delve deeper into the *lonely cluster*, where valuable insights resided, and target individual endpoints. Cluefish enabled us to formulate hypotheses supported by multiple concordant elements that not only make biological sense but also hold a major scientific interest. These insights primarily arise from exploring three of the four most sensitive clusters. While a more comprehensive analysis could uncover additional roads for biological interpretation, a complete biological interpretation is beyond the scope of this paper.

An important consideration in biological interpretation approaches for transcriptomic and other omics data is the organism under study. Gene annotation richness varies substantially across species, typically reflecting whether the organism is an established model system with extensive reference databases or a less-studied species with limited annotation. A critical aspect of validating Cluefish’s capacity to enhance biological interpretation was testing its performance across diverse organisms. To assess Cluefish’s versatility and generalisability, we applied it to two externally published dose-response transcriptomic datasets: rat liver exposed to perfluorooctanoic acid (PFOA) and poplar roots exposed to phenanthrene (PHE).

In the rat liver dataset, cluster #3 emerged as the most sensitive to PFOA exposure. This cluster was characterised by enrichment in cellular response to lipid (GO:0071396), superoxide metabolic process (GO:0006801), positive regulation of tumor necrosis factor production (GO:0032760), and activation of innate immune response (GO:0002218). These findings suggest that even the lowest doses of PFOA may trigger inflammation, oxidative stress, and lipid metabolism disruption. While the original study did not include detailed biological interpretation, our results align with existing literature on PFOA toxicity mechanisms (84–88).

For the poplar root dataset, the standard approach identified several redundant biological functions with substantial gene overlap. In contrast, Cluefish detected more specific processes within similar branches of the GO term hierarchy. In the original study, authors reported enrichment in oxidative stress response, cellular redox homeostasis, defense response, and biotic stimuli response pathways. Cluefish successfully identified these same, or similar, functions but also provided additional resolution. Notably, the authors hypothesised that PHE induces ethylene signalling pathway responses, yet this biological function was not identified as enriched in their analysis nor the standard approach in this study, illustrating the challenges of developing complex hypotheses without prior knowledge of potential mechanisms.

Cluefish, however, revealed biological functions directly supporting this hypothesis, including ethylene-activated signalling pathway (GO:0006979), hormone-mediated signalling pathway (GO:0009755), and response to toxic substance (GO:0097237). This demonstrates how Cluefish can reduce the need for extensive manual mining of data functional annotation and allow for a more automatic exploration independent of researcher prior knowledge.

The preliminary insights gained from these external datasets demonstrate Cluefish’s potential, though a more comprehensive analysis would be necessary to plan targeted experimental validations and fully elucidate the molecular mechanisms underlying toxicant effects.

The Cluefish workflow involves several filters, and selecting appropriate values can be challenging. One key parameter is the lower cluster size filter, which excludes clusters based on gene set size. Users should evaluate the total number of clusters, the number of genes contributing to enrichment, and the number of lonely genes successfully fished. The goal is to find a balance between these metrics and identify the point at which the lonely gene fishing process begins to plateau. Other filters depend on factors such as the organism (from poorly to well-annotated genomes), the type of transcriptomic data (e.g., whole transcriptome vs. tissue-specific), and choices made in previous steps. We recommend exploring a range of parameter settings to better understand their effects and to fine-tune the workflow for each specific dataset.

Clustering is essential in RNA-seq data analysis, grouping genes or samples by shared traits like expression patterns or network connections (89, 90). A widely used approach is weighted correlation network analysis (WGCNA) (91), which identifies co-regulated genes by constructing co-expression networks from their expression profiles across samples (92). The underlying hypothesis behind WGCNA is that coregulated genes are functionally related. However, due to the complexity and multi-faceted interconnectivity of organismal pathways, this can result in clusters of co-regulated but functionally unrelated genes. In our approach, DRomics first identifies a list of deregulated transcripts. Then, using STRING within Cytoscape, we group genes based on known and predicted interactions, directly forming clusters of functionally relevant genes without relying on gene expression patterns. This approach avoids clustering functionally unrelated genes and allows us to characterise the clusters and their genes using the results of DR modelling, within a non-targeted approach. PPI network construction and clustering are often applied post-enrichment to visualise relationships between gene sets, using tools like GSCluster (93) and NeVOmics (94).

Other tools like EnrichmentMap (95) and ClueGO (96), both extensions of Cytoscape, help reduce redundancy in biological functions via PPI networks by accounting for both the overlap and interactions between gene products. Recently, clustering PPI networks before enrichment has gained traction, as seen in tools like PathFindR (97) and ANUBIX (98). This approach improves enrichment sensitivity, uncovering hidden biological mechanisms (99). Though both tools share the pre-clustering concept with Cluefish, their application to lesser-referenced organisms is limited. ANUBIX uses constrained randomisation to evaluate subnetwork relevance, making it optimal for highly comprehensive PPI networks. PathFindR, primarily designed for human datasets, is applicable to non-human organisms but enrichment is limited to KEGG, Reactome (100) and MSigDB databases, with the latter two being restricted to very few model organisms. Cluefish, by contrast, seeks to overcome these limitations by incorporating a functional enrichment tool that supports a broader range of organisms. Nevertheless, challenges still remain for lesser-referenced species.

While Cluefish improves traditional ORA, it has some limitations. Cluefish relies on STRING’s PPI data, which can be limiting for transcriptional regulators. For instance, Hox genes, despite being functionally linked, often cluster separately due to a lack of direct interactions (101). This also affects other transcriptional regulators, many of which fall into the *lonely cluster*. Additionally, Cluefish remains semi-automated due to a manual Cytoscape step. Full automation could be achieved by integrating this step into R using the RCy3 package (102), but querying large gene lists and clustering complex networks is still computationally challenging. We plan to implement full automation as soon as it becomes feasible.

Despite these limitations, enhancing the methods and databases used in Cluefish could be highly beneficial. Firstly, a more comprehensive set of databases would greatly increase the workflow’s versatility and applicability. For instance, expanding PPI data to include lesser-studied organisms would broaden its scope, while extending TFlinks (103) —a database of transcription factor-target gene relationships—to cover a wider range of model organisms would significantly enhance biological interpretation in a mechanistic context. Secondly, a major challenge is the complexity and redundancy of biological functions, particularly in GO terms (104, 105). To reduce redundancy, we applied Alexa et al.’s (2006) method (106), which selects representative terms using the hierarchical structure of the GO tree. This seems to be a more effective approach than semantic similarity methods based solely on gene overlap (106). KEGG pathways also face redundancy issues, and while some methods address this (97, 107), simplifying biological functions remains challenging. Finally, ORA assumes that both genes and pathways are independent, meaning that neither topology nor pathway overlap is considered, which limits its scope. Newer enrichment methods (e.g., topology-based) show promise but are not yet compatible with DR modelling metrics, which Cluefish uses. These methods also require highly complete datasets, limiting their application to well-referenced organisms like humans and mice. To enhance Cluefish’s functionality, tools that handle metrics beyond expression values, like BMD and trend metrics, are needed.

To the best of our knowledge, Cluefish is the first workflow to integrate DR modelling with the functional enrichment of pre-clustered networks. This comprehensive approach allows for the creation of biologically meaningful clusters, characterised by sensitivity and trend derived from DR analysis. With innovative features such as cluster merging and *lonely gene* fishing, Cluefish provides users with a powerful tool for a more thorough interpretation of transcriptomic datasets. By offering clearer insights, it facilitates the generation of biological hypotheses and guiding experimental validation.

To further establish Cluefish’s advantages over standard methods and demonstrate its versatility, future work should apply the workflow to additional DR datasets. Testing across more organisms, spanning from poorly to well-annotated genomes, and varying numbers of deregulated transcripts will allow for a deeper evaluation of its robustness and broader applicability.

While Cluefish employs DR modelling to identify deregulated transcripts and visually explore the results, its core functionality extends beyond this specific tool or analysis type. Cluefish can process a background list of transcripts along with a sub-list, typically consisting of deregulated transcripts. These deregulated transcripts can be derived from a wide range of experimental designs, from the simplest to the more complex ones, such as time or spatial data series. Therefore, it would be interesting to explore the application of Cluefish to other types of data series.

## Supporting information

Supplementary File S3 - Report Ex. 1

Supplementary File S4 - Report Ex. 2

Supplementary File S5 - Report Ex. 3

## DATA AVAILABILITY

The Cluefish project is available in its developmental version on the Github repository (https://github.com/ellfran-7/cluefish). The specific version used to produce the results in this paper is available in the Software Heritage archive at https://archive.softwareheritage.org/browse/origin/directory/?origin_url, https://github.com/ellfran-7/cluefish, and can be accessed with the following Software Heritage persistent identifier (SWHID): swh:1:dir:555002e59cc81f71df7fe9a16712483a7e59c77b. A detailed step-by-step guide of the workflow is available in the form of a vignette at https://ellfran-7.github.io/cluefish/articles/cluefish.html. The main zebrafish transcriptomic supporting this study is publicly available on NCBI Gene Expression Omnibus (GEO) at https://www.ncbi.nlm.nih.gov/geo/, and can be accessed with GSE283957. Individual zebrafish morphometric data (left eye area and body length measurements) are provided as supplementary material (Supplementary Table S23). Additionally, two external transcriptomic datasets were utilised: the rat liver dataset from Gwinn *et al.* (accession GSE147072) and the poplar root dataset from Gréau *et al.* (accession GSE263776), both of which are also publicly available through the GEO database.

## SUPPLEMENTARY DATA

Supplementary Data are available at NAR online.

## AUTHOR CONTRIBUTIONS

Ellis Franklin: Conceptualisation, Data curation, Formal analysis, Methodology, Software, Writing—original draft, Writing—review & editing. Elise Billoir: Conceptualisation, Methodology, Supervision, Validation, Writing—review & editing. Philippe Veber: Conceptualisation, Methodology, Writing—review & editing. Jérémie Ohanessian: Investigation, Validation, Writing—review & editing. Marie Laure Delignette-Muller: Conceptualisation, Methodology, Supervision, Validation, Writing— review & editing. Sophie M. Prud’homme: Conceptualisation, Data curation, Investigation, Methodology, Supervision, Validation, Writing—original draft, Writing— review & editing.

## ACKNOWLEDGEMENTS

This work was partly done in the “Pôle de compétences en biologie environnementale” and the “Pôle de compétences en chimie analytique environnementale”, ANATELo, LIEC laboratory, UMR 7360 CNRS – Université de Lorraine. We would like to express our thanks to Maria Lorena Cordero Maldonado for providing us with the zebrafish embryos, Carole Cossu-Leguille for her assistance with zebrafish larvae sampling, to Bénédicte Sohm for performing RNA extraction and quality checks, and to Claire Caillard for preparing the samples for morphological measurements. Their invaluable contributions significantly enhanced the quality of this research. The raw mRNAseq data processing was performed on the Core Cluster of the Institut Français de Bioinformatique (IFB) (ANR-11-INBS-0013).

