## Supplementary File 1 - Text and Figures for "Cluefish: mining the dark matter of transcriptional data series with over-representation analysis enhanced by aggregated biological prior knowledge"

**Text S1. Parameter choices for Cluefish applied to the external datasets**

**Rat liver dataset**

For the rat liver (*Rattus norvegicus*) dataset, we downloaded the TF and CoTF gene lists (Step 1). We extracted gene names from the dataset's original custom identifiers, then converted these to their corresponding Ensembl gene IDs (*Step 3*). We maintained the default confidence score (**0.9**) in STRING for the protein-protein interaction network and the inflation parameter (**4**) for MCL clustering, similar to the zebrafish dataset, due to the high-quality annotation and interaction data available for this species (*Step 5*). The lower cluster size filter was kept at **4** to maintain consistency in cluster definition (*Step 6*). The main adjustments were made to the functional enrichment parameters in *Step 7* to account for the tissue-specific nature of this dataset. As by default, we tested against GO:BP, KEGG and WP pathways. As we were working with liver-specific expression data rather than whole-organism transcriptomics, we applied a more restrictive biological function size filter (lower and upper limits of **10** and **130** genes, respectively) to focus on liver-relevant processes and exclude overly broad or tissue-irrelevant functions. The enrichment gene count filter was increased to **4** to align with the higher lower-limit of biological function size and to avoid enrichments driven by only one or two genes. The friendliness parameter was set to **2** to prevent overly promiscuous genes (such as central metabolic regulators) from being incorporated into multiple clusters, which could dilute cluster specificity (*Step 9*).

**Poplar root dataset**

For this dataset concerning *Populus trichocarpa* (to which the studied *Populus canadensis* was mapped), several parameter adjustments were necessary due to its status as a less-referenced organism. No TF or CoTF gene lists were available, therefor we skipped the regulatory status annotation step (*Step 1*). We converted the initial Ensembl gene IDs to UniProtKB/TrEMBL IDs, required in this case for the protein-correspondance in the STRING database (*Step 3*). To compensate for the more limited data on protein interaction in this non-model organism, we reduced the STRING confidence score to **0.6**, allowing more predicted interactions to be included in the network. We also adjusted the MCL inflation parameter to **2** (versus the default 4) to create fewer but larger clusters, preventing excessive fragmentation of the network into many small clusters with low confidence (*Step 5*). The lower cluster size filter was maintained at **4** (*Step 6*). For functional enrichment analysis, we tested against GO:BP and KEGG only, as the WP database lacked sufficient coverage for this organism. We retained a broad range for the biological function size filter (**5**-**800** genes) to accommodate the whole-transcriptome background and the less comprehensive functional annotation of poplar genes (*Step 7*). The enrichment gene count filter was reduced to **2**, allowing enrichment to be detected even with fewer genes per function, which is particularly important for smaller clusters in this less-annotated species. As with the rat dataset, the friendliness parameter was set to **2** to maintain cluster specificity while fishing lonely (unassigned) genes (*Step 9*).

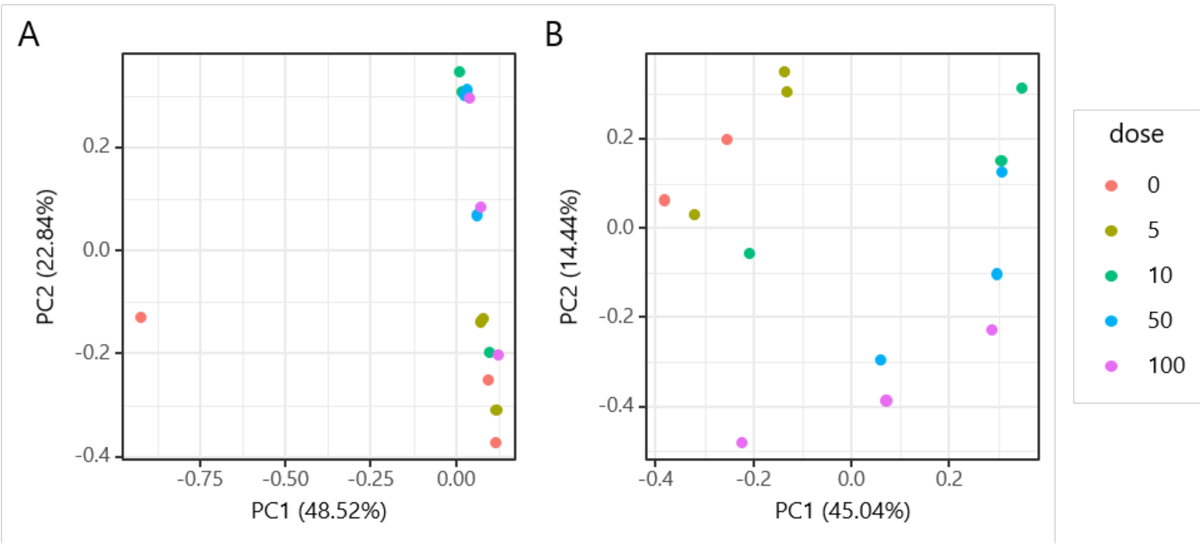

**Figure S1. Principal Component Analysis (PCA) plots showing the data before (A) and after (B) removal of one control sample (replicate).**

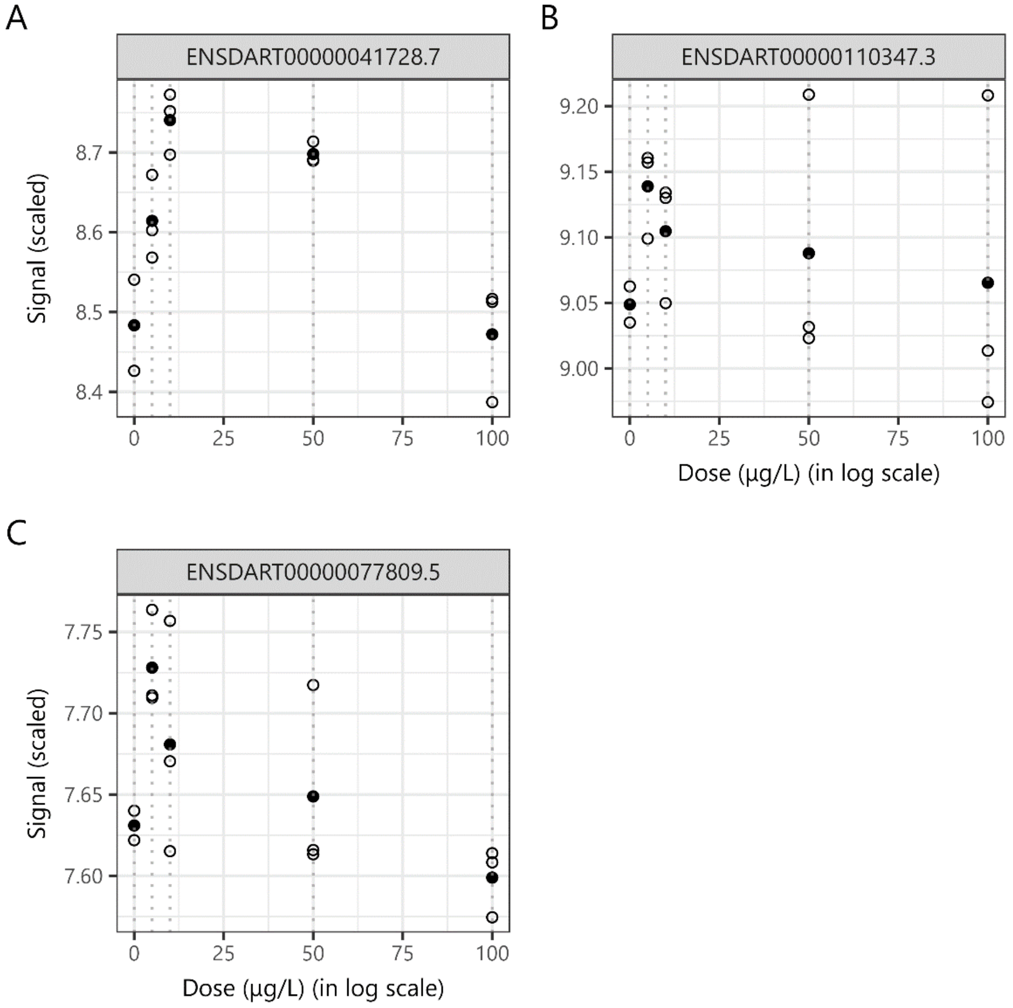

**Figure S2. Plots of the raw dose-response data for the transcripts of three Cyp26 genes: (A) cyp26a1 (ENSDART00000041728.7), (B) cyp26b1 (ENSDART00000110347.3), and (C) cyp26c1 (ENSDART00000077809.5).** Grey dashed vertical lines represent the experimental doses.

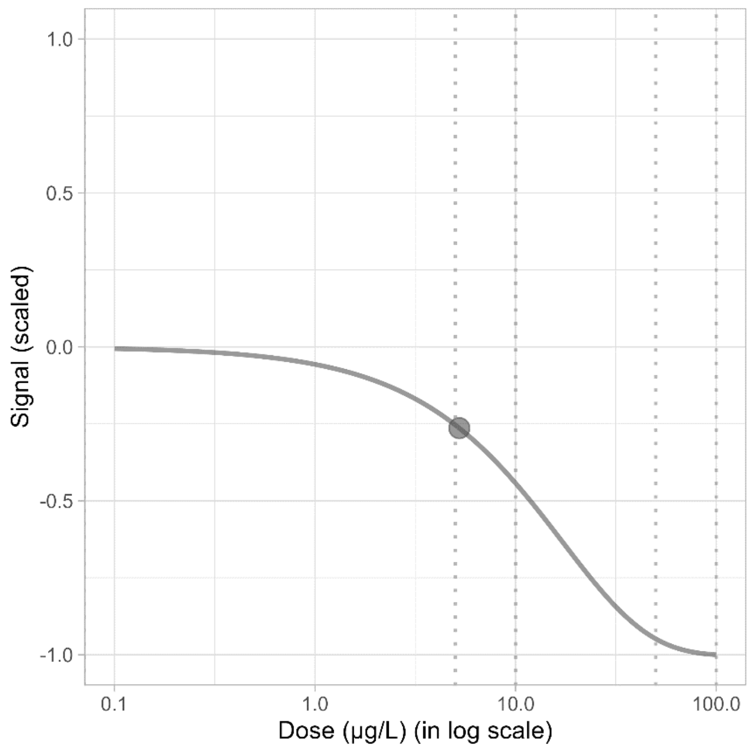

**Figure S3. Dose-response fitted curve and computed BMD point for the rxraa gene transcript (ENSDART00000080481.6).** Grey dashed vertical lines represent the experimental doses.
