## Supplementary File S3 - Report Ex. 1 for "Cluefish: mining the dark matter of transcriptional data series with over-representation analysis enhanced by aggregated biological prior knowledge"

### Report comparison of standard and cluefish workflow

AUTHOR
Ellis Franklin

PUBLISHED
2024-08-07

#### 1 Introduction

The cluefish workflow seeks to enhance the biological interpretation of transcriptomic dose-response modeling data, particularly following the DRomics analysis. Its primary goal is to comprehensively characterize the data while ensuring an unbiased and non-targeted approach. By doing so, we aim to not only maximize the depth of the analysis but also maintain objectivity throughout.

To evaluate the efficacy of this workflow, we can contrast its outcomes with those of a standard workflow, which typically involves functional enrichment analysis on the single list of deregulated transcript genes. This comparative analysis will provide insights into the strengths and potential advantages of cluefish.

#### 2 Comparing initial enriched term content

Here we exclusively compare the enriched terms identified by the two distinct workflows: the standard workflow, which uses the entire gene list as query for the functional enrichment, and the cluefish workflow, which uses all clusters as individual queries. This comparison is conducted separately for each data source.

> **Important**
> This comparison solely focuses on the enriched terms and without any filtering performed. Thus, the results for the cluefish workflow are derived specifically from the **clustrenrich$gostres** output of the **clustrenrich()** function.

##### Standard workflow results

Show 5 entries

Search:

| | term_name | term_id | term_size | query_size | intersection_size | p_value | e |
|---|---|---|---|---|---|---|---|
| 1 | peptide biosynthetic process | GO:0043043 | 355 | 1509 | 113 | 5.555665588728927e-28 | |
| 2 | translation | GO:0006412 | 351 | 1509 | 111 | 1.870374801556167e-27 | |
| 3 | amide biosynthetic process | GO:0043604 | 408 | 1509 | 121 | 2.2841852128196e-27 | |

| | term_name | term_id | term_size | query_size | intersection_size | p_value | e |
|---|---|---|---|---|---|---|---|
| 4 | peptide metabolic process | GO:0006518 | 408 | 1509 | 119 | 2.916188044323696e-26 | |
| 5 | amide metabolic process | GO:0043603 | 529 | 1509 | 133 | 9.255574026569427e-23 | |

Showing 1 to 5 of 95 entries    Previous  1  2  3  4  5  …  19  Next

Table 1: Standard workflow functional enrichment results

#### cluefish workflow results

Show 5 entries    Search:

| | query | term_name | term_id | term_size | query_size | intersection_size | p_ |
|---|---|---|---|---|---|---|---|
| 1 | 1 | translation | GO:0006412 | 351 | 31 | 22 | 8.5805442[illegible] |
| 2 | 1 | ribonucleoprotein complex biogenesis | GO:0022613 | 175 | 31 | 8 | 2.18651280[illegible] |
| 3 | 1 | SRP-dependent cotranslational protein targeting to membrane | GO:0006614 | 14 | 31 | 2 | 0.002636779 |
| 4 | 1 | embryo development ending in birth or egg hatching | GO:0009792 | 180 | 31 | 4 | 0.00309650 |
| 5 | 1 | myeloid cell differentiation | GO:0030099 | 89 | 31 | 3 | 0.004890742 |

Showing 1 to 5 of 289 entries    Previous  1  2  3  4  5  …  58  Next

Table 2: cluefish workflow functional enrichment results

#### Venn diagrams between both workflow per source

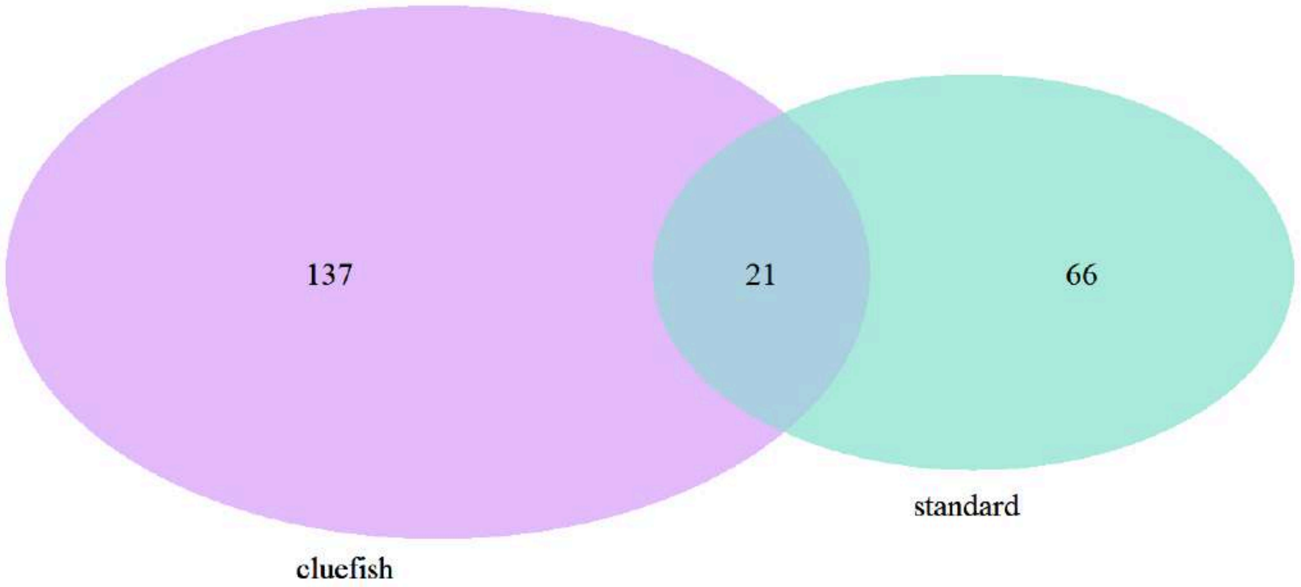


Figure 1: Venn diagram of highlighted enriched GO terms between the standard and cluefish workflow

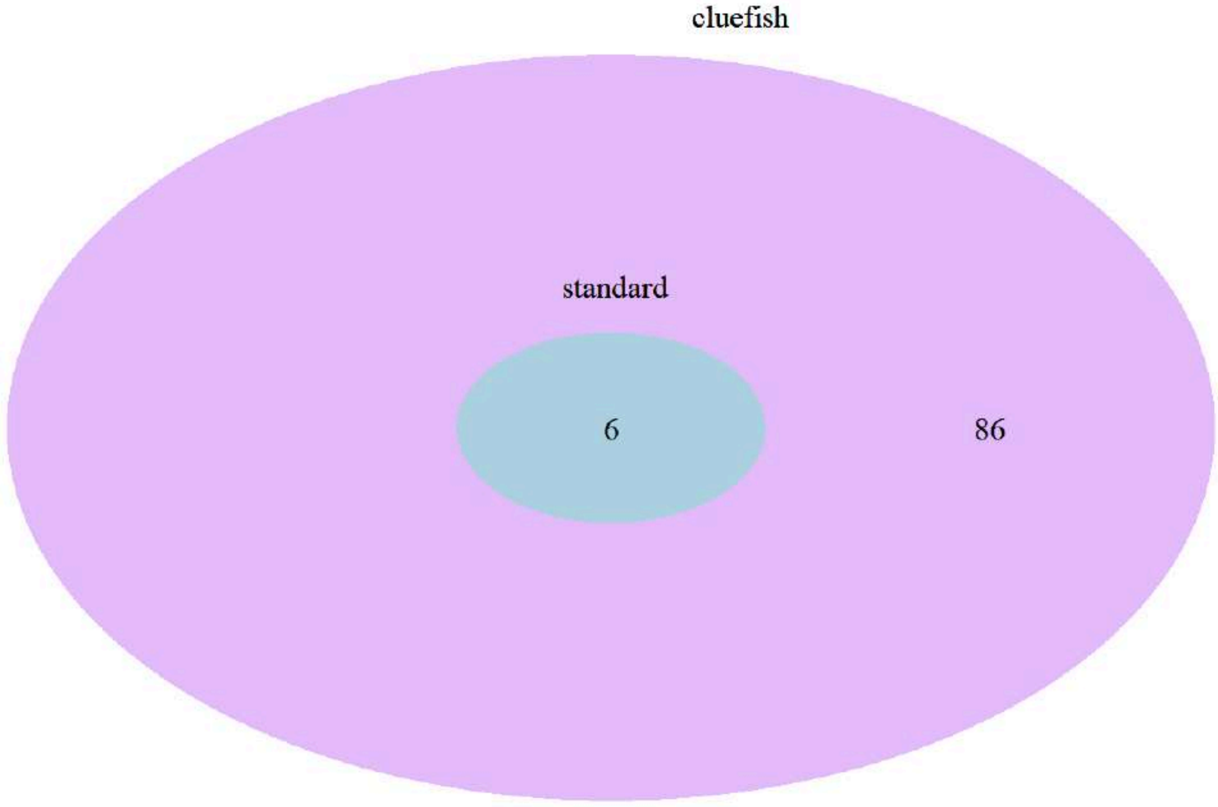


Figure 2: Venn diagram of enriched KEGG pathways between the standard and cluefish workflow

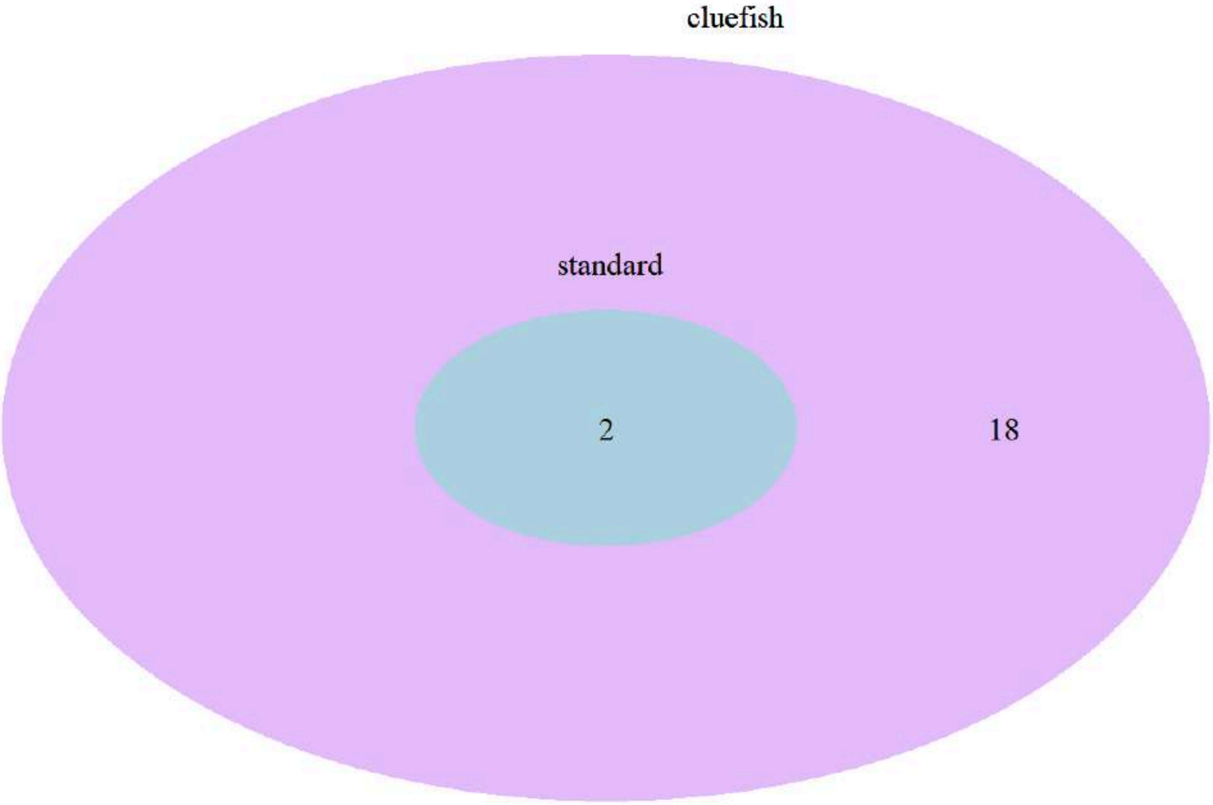


Figure 3: Venn diagram of enriched Wikipathways between the standard and cluefish workflow

### 3 Comparing enriched term content after filters in the cluefish workflow

With filters applied to the enriched terms within following steps of the standard and cluefish workflow, we can re-examine the discrepancies between enriched terms considered between both workflow.

#### Standard workflow results

Show 5 entries Search:

| | term_name | term_id | term_size | query_size | intersection_size | p_value |
|---|---|---|---|---|---|---|
| 1 | peptide biosynthetic process | GO:0043043 | 355 | 1509 | 113 | 5.555665588728927e-28 |
| 2 | lens development in camera-type eye | GO:0002088 | 70 | 1509 | 28 | 4.10849912135549e-9 |
| 3 | protein folding | GO:0006457 | 126 | 1509 | 31 | 0.0001116792401074228 |
| 4 | DNA replication | GO:0006260 | 92 | 1509 | 22 | 0.003734667164216148 |
| 5 | intracellular protein transmembrane transport | GO:0065002 | 14 | 1509 | 7 | 0.00928929320209972 |

Showing 1 to 5 of 20 entries

Previous 1 2 3 4 Next

Table 3: Standard workflow functional enrichment results

#### cluefish workflow results

Show 5 entries Search:

| | query | term_name | term_id | term_size | query_size | intersection_size | p_ |
|---|---|---|---|---|---|---|---|
| 1 | 1 | Cytoplasmic ribosomal proteins | WP:WP324 | 60 | 19 | 18 | 1.42532120 |
| 2 | 1 | embryo development ending in birth or egg hatching | GO:0009792 | 180 | 31 | 4 | 0.00309650 |
| 3 | 1 | myeloid cell differentiation | GO:0030099 | 89 | 31 | 3 | 0.004890742 |
| 4 | 1 | ribonucleoprotein complex biogenesis | GO:0022613 | 175 | 31 | 8 | 2.18651280 |
| 5 | 1 | Ribosome | KEGG:03010 | 102 | 21 | 19 | 6.38552598 |

Showing 1 to 5 of 133 entries Previous 1 2 3 4 5 … 27 Next

Table 4: cluefish workflow functional enrichment results

#### Venn diagrams between both workflows per source after filters in the cluefish workflow

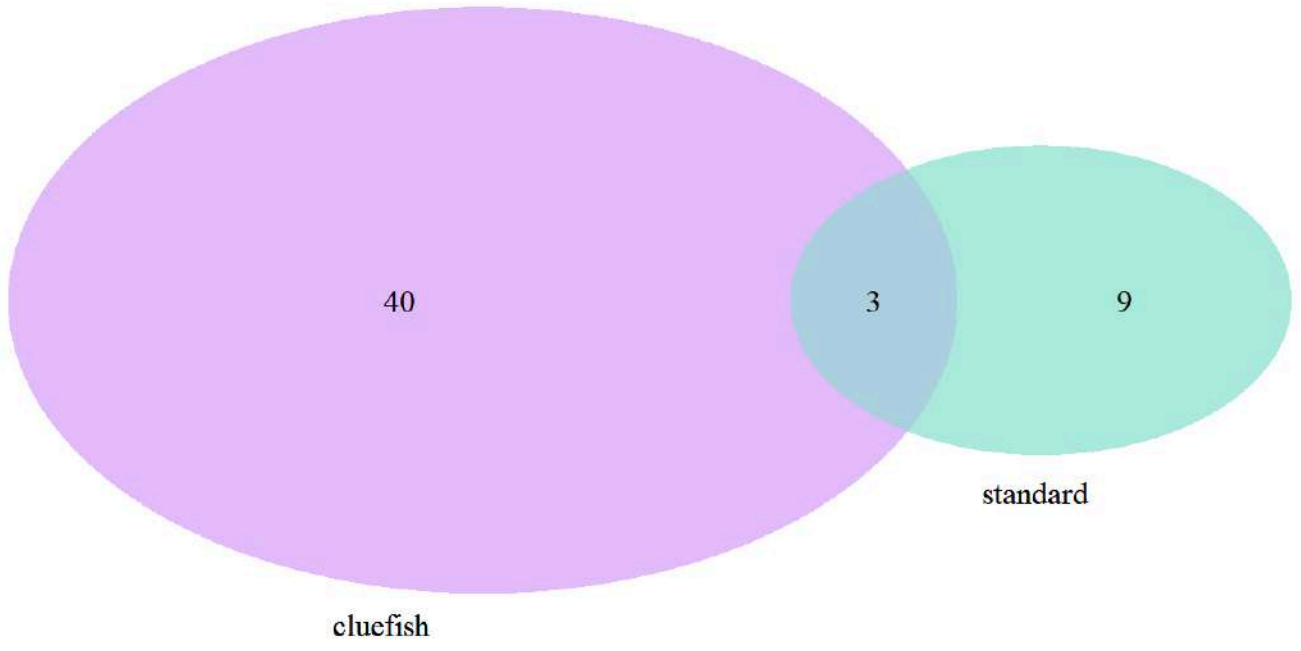


Figure 4: Venn diagram of highlighted enriched GO temrs in the standard and filtered highlighted enriched GO terms in the cluefish workflow

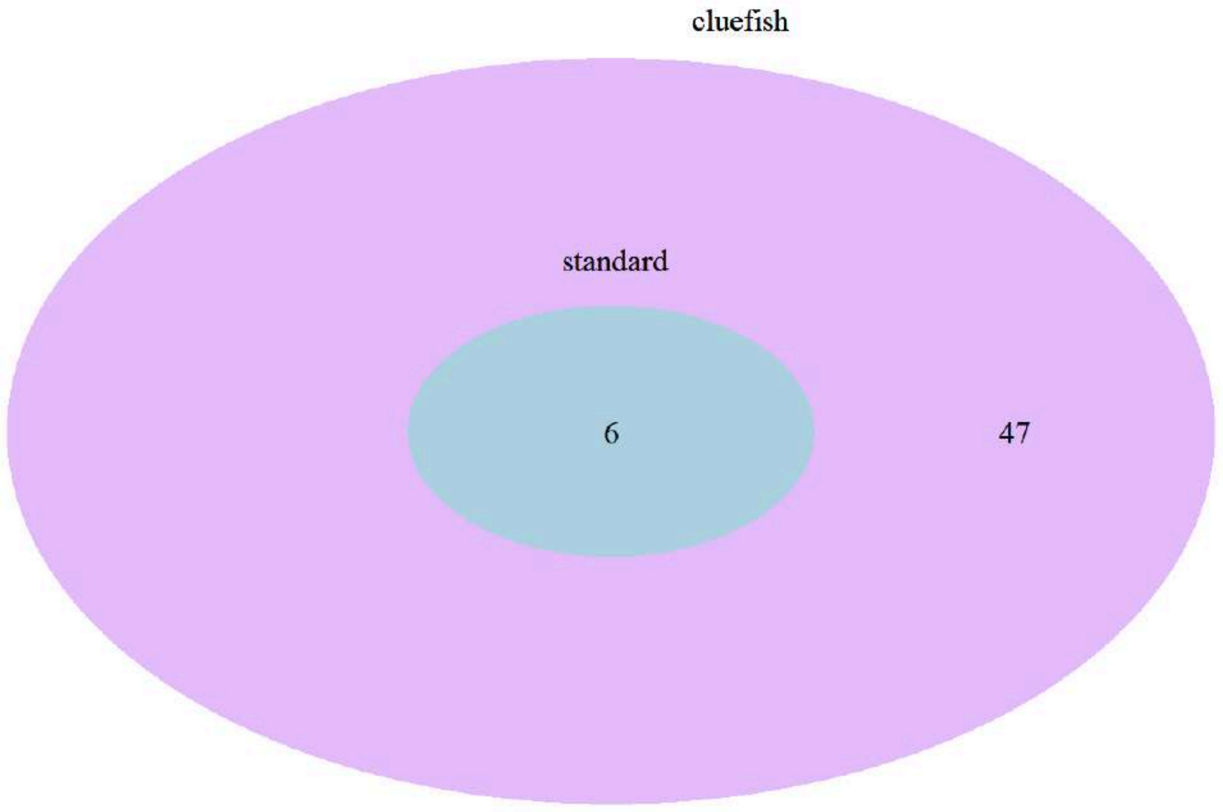


Figure 5: Venn diagram of enriched KEGG pathways in the standard and filtered enriched KEGG pathways in the cluefish workflow

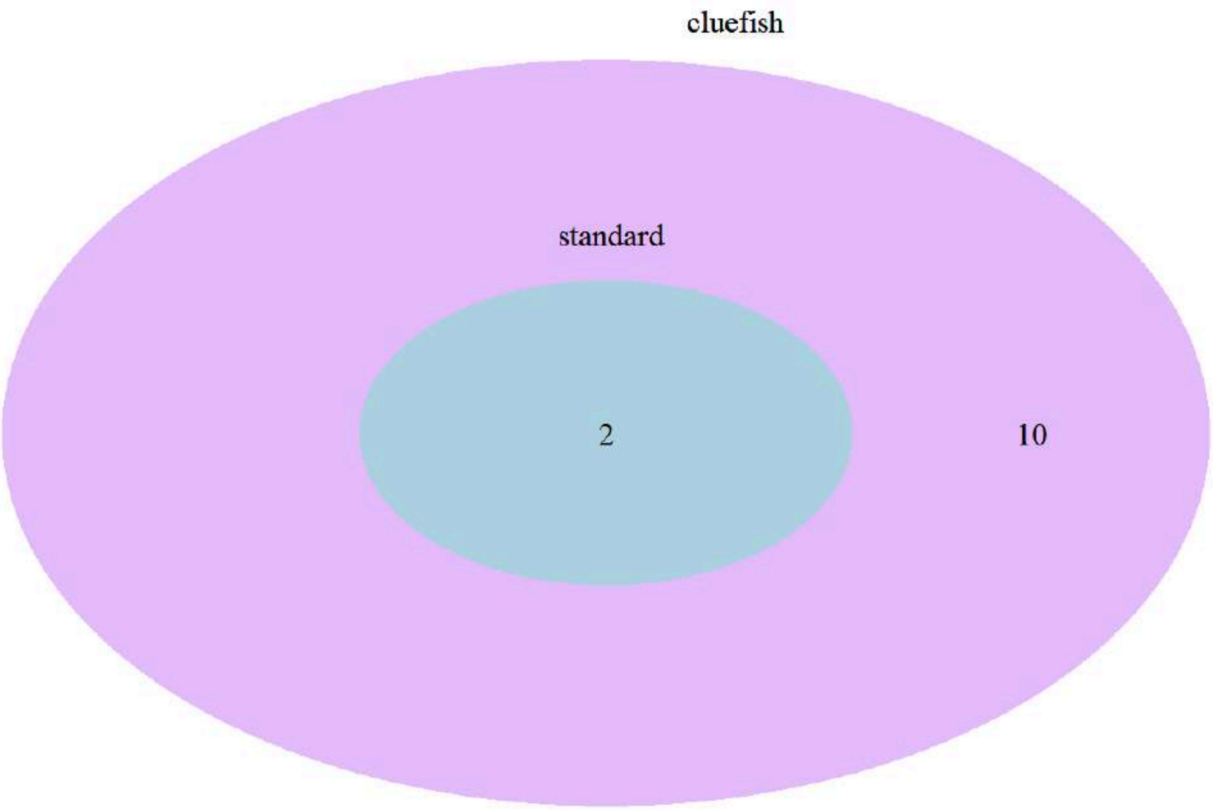


Figure 6: Venn diagram of enriched Wikipathways in the standard and filtered enriched Wikipathways in the cluefish workflow

#### 4 A short summary

The extent of what we can gather, or *what are we working with*, is based on the following question: *how many transcript are considered in the interpretation phase?* And *what is left to be overlooked?*

| Metric | Standard | cluefish |
|---|---|---|
| Number of transcripts considered | 332 | 940 |
| Number of lonely transcripts | 2101 | 1493 |

Table 5: Summary of the comparison between both workflow

However, within the cluefish workflow, the lonely cluster consisting of all the lonely content (transcripts/genes) remains open for exploration. Within the **“lonely_results_report.qmd”**, an additional functional enrichment analysis is conducted using the lonely gene list as the query. This approach enables us to delve into what might be overlooked and whether there is any biological significance within this group.
