## Supplementary File S4 - Report Ex. 2 for "Cluefish: mining the dark matter of transcriptional data series with over-representation analysis enhanced by aggregated biological prior knowledge"

### Report of the lonely cluster results

AUTHOR
Ellis Franklin

PUBLISHED
2024-08-07

#### 1 Introduction

This document is a comprehensive report, meticulously encapsulating the pivotal results derived from the DRomics analysis while detailing the workflow. It succinctly summarizes the impacts and outputs of each significant step within the pipeline, supported by visual representations like plots depicting the summary of BMD values per cluster, aiding in prioritization. Moreover, additional specific visualizations for each cluster, such as plots of fitted curves, empirical cumulative distribution function (ECDF) plots of BMD values, and summary tables, enable a more focused exploration.

This document serves as a supplementary report to the workflow_results_report, focusing on the Lonely cluster. The Lonely cluster comprises genes that remain unassociated with any other cluster throughout the workflow process.

The lonely cluster is composed of a total of 1493.

Functional enrichment analysis is conducted to link the cluster with biological processes, facilitating deeper exploration of aspects potentially overlooked in the analysis.

#### 2 The Lonely cluster as an interactive table

Table 1: Summary table of the lonely cluster

Show 10 entries Search:

| | transcript_id | gene_id | gene_name | NewCluster | Friendliness | Term_name | Source |
|---|---|---|---|---|---|---|---|
| 1 | ENSDART00000000069.8 | ENSDARG00000000068 | slc9a3r1a_g2t1 | Lonely | 1 | | |
| 2 | ENSDART00000000070.7 | ENSDARG00000000069 | dap_g2t1 | Lonely | 1 | | |
| 3 | ENSDART00000001691.8 | ENSDARG00000001463 | tdh2 | Lonely | 1 | Glycine, serine and threonine metabolism | KEGG |
| 4 | ENSDART00000001691.8 | ENSDARG00000001463 | tdh2 | Lonely | 1 | Metabolic pathways | KEGG |
| 5 | ENSDART00000002164.9 | ENSDARG00000008433 | unc45b_t2 | Lonely | 1 | | |
| 6 | ENSDART00000002279.8 | ENSDARG00000021442 | cdh11 | Lonely | 1 | | |
| 7 | ENSDART00000002398.7 | ENSDARG00000018264 | trim101 | Lonely | 1 | | |
| 8 | ENSDART00000002469.7 | ENSDARG00000018989 | hspa4b_g2t1 | Lonely | 1 | | |
| 9 | ENSDART00000003133.10 | ENSDARG00000021399 | yipf2_t2 | Lonely | 1 | | |
| 10 | ENSDART00000003170.5 | ENSDARG00000018145 | mid1ip1l | Lonely | 1 | | |

Showing 1 to 10 of 1,667 entries

Previous 1 2 3 4 5 … 167 Next

#### 3 Characterizing the Lonely cluster by functional enrichment

| Source | Query size | Background size |
|---|---|---|
| GO:BP | 763 | 15468 |
| KEGG | 177 | 5714 |
| WP | 35 | 1383 |

Table 2: Number of genes involved in the query and background of the ORA

Show 10 entries | Search:

| | term_name | term_id | source | term_size | intersection_size | query_size | effective_domain_ |
|---|---|---|---|---|---|---|---|
| 1 | lens development in camera-type eye | GO:0002088 | GO:BP | 70 | 26 | 763 | |
| 2 | visual perception | GO:0007601 | GO:BP | 139 | 26 | 763 | |
| 3 | transition metal ion transport | GO:0000041 | GO:BP | 41 | 9 | 763 | |
| 4 | regulation of small GTPase mediated signal transduction | GO:0051056 | GO:BP | 106 | 15 | 763 | |
| 5 | positive regulation of leukocyte migration | GO:0002687 | GO:BP | 15 | 5 | 763 | |
| 6 | regulation of synapse organization | GO:0050807 | GO:BP | 23 | 6 | 763 | |
| 7 | Endocytosis | KEGG:04144 | KEGG | 268 | 22 | 177 | |
| 8 | Lysosome | KEGG:04142 | KEGG | 143 | 14 | 177 | |
| 9 | Polycomb repressive complex | KEGG:03083 | KEGG | 72 | 9 | 177 | |
| 10 | Ferroptosis | KEGG:04216 | KEGG | 46 | 6 | 177 | |

Showing 1 to 10 of 12 entries

Previous 1 2 Next

#### 4 The results as an interactive curvesplot

**Cluster Lonely - 1493 transcripts**

Curvesplot colored by trend | BMDplot colored by trend

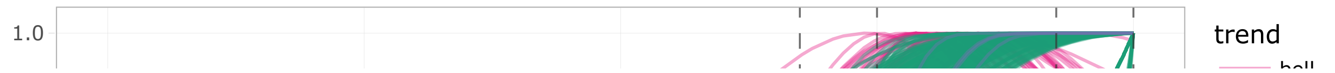

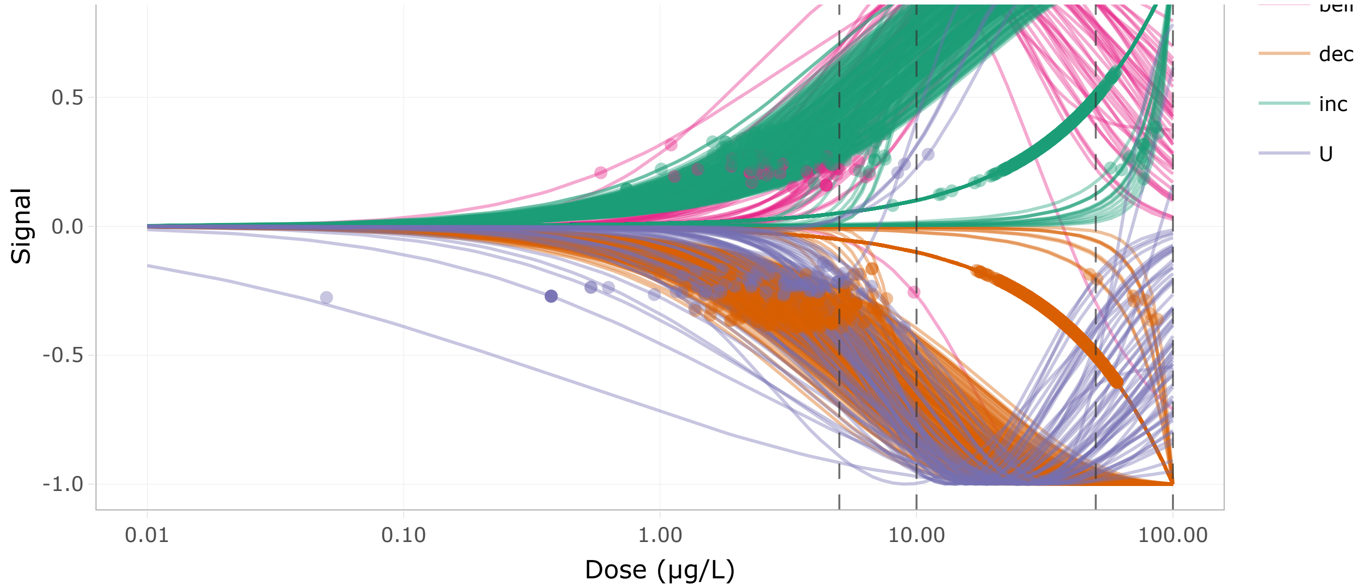


Figure 1: DR curves for the Lonely cluster colored by trend

Curvesplot colored by enriched term

BMDplot colored by enriched term

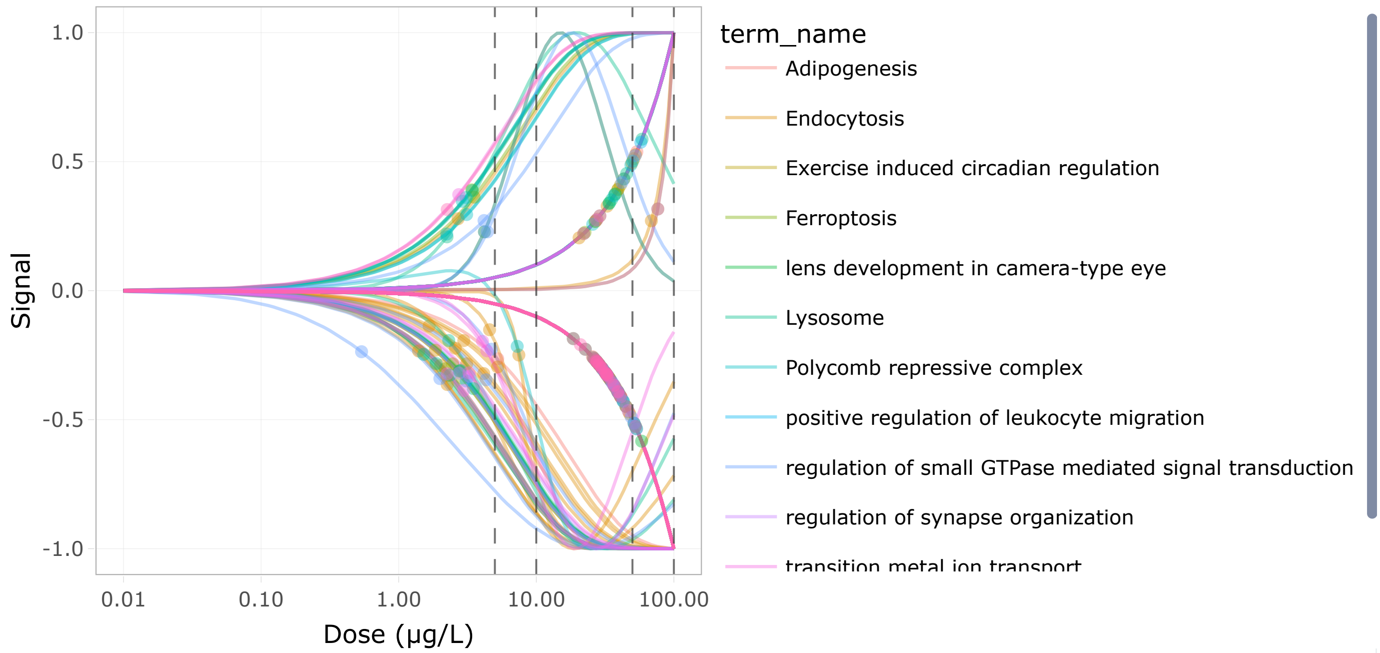


Figure 3: DR curves for all genes associated to the terms enriched by the lonely cluster
