## Supplementary File S5 - Report Ex. 3 for "Cluefish: mining the dark matter of transcriptional data series with over-representation analysis enhanced by aggregated biological prior knowledge"

### Report of the workflow results

AUTHOR
Ellis Franklin

PUBLISHED
2024-08-19

#### 1 Introduction

This document is a comprehensive report, meticulously encapsulating the pivotal results derived from the DRomics analysis while detailing the workflow. It succinctly summarizes the impacts and outputs of each significant step within the pipeline, supported by visual representations like plots depicting the summary of BMD values per cluster, aiding in prioritization. Moreover, additional specific visualizations for each cluster, such as plots of fitted curves, empirical cumulative distribution function (ECDF) plots of BMD values, and summary tables, enable a more focused exploration.

#### 2 Results from the DRomics analysis

The total number of transcripts derived from this experiment : **39890**

| Step | Number of transcripts concerned |
|---|---|
| Selection of deregulated transcripts | 2449 |
| Computation of BMD-1SD values | 2449 |
| Defined confidence interval around BMD | 2433 |

Table 1: Summary table of DRomics analysis

#### 3 Summary of the workflow

##### Identifier retrieval

| Type of identifier | Background count | Deregulated count |
|---|---|---|
| Ensembl transcript | 39890 | 2433 |
| Ensembl gene | 27720 | 2365 |

Table 2: Summary table of indentifier mapping

**Note**

The external_gene_name identifiers, which depict human-readable gene names, have been adjusted to address redundancy between Ensembl transcript or gene identifiers. As a result, the count of external_gene_name IDs matches the number of transcripts: 2433.

##### Cluster Gene Set Size Filtering

Clusters containing at least **4** Ensembl genes are kept for subsequent steps. Genes in the remaining clusters are designated as “lonely genes”, signifying their absence from any cluster.

| Step | Ensembl Gene Count | Cluster Count |
|---|---|---|
| Before clustrfiltr() | 750 | 204 |
| After clustrfiltr() | 399 | 53 |

Table 3: Summary table of cluster and gene count before and after cluster filtering step

#### Cluster Functional Enrichment

Functional enrichment analysis is conducted individually for each cluster and data source. The parameters used in the `clustrenrich()` function to generate this output are as follows:

- background type ( `bg_type` ) = **custom_annotated**
- data sources ( `sources` ) = **GO:BP, KEGG, WP**
- p-value ( `user_theshold` ) = **0.05**
- multiple testing correction method ( `correction_method` ) = **fdr**
- minimum term size ( `min_term_size` ) = **5**
- maximum term size ( `max_term_size` ) = **500**
- choice of keeping only highlighted GO terms ( `only_highlighted_GO` ) = **TRUE**
- Number of genes required to keep enrichment ( `ngenes_enrich_filtr` ) = **3**

| Metric | GO:BP | KEGG | Wikipathways |
|---|---|---|---|
| Background size | 15468 | 5714 | 1383 |

Table 4: Summary of background gene lists involved in functional enrichment for each source

| Metric | Query Ensembl Gene Count | Cluster Count | Total Term Count |
|---|---|---|---|
| Enrichment results before term filtering | 316 | 52 | 218 |
| Enrichment results after term filtering | 286 | 47 | 97 |

Table 5: Summary table of gene, cluster and total term count during the cluster enrichment step

| Metric | GO:BP | KEGG | Wikipathways |
|---|---|---|---|
| All terms associated with deregulated genes | 3450 | 164 | 43 |

| Metric | GO:BP | KEGG | Wikipathways |
|---|---|---|---|
| Enriched terms | 114 | 91 | 18 |
| Filtered enriched terms | 41 | 51 | 10 |

Table 6: Summary table of term count per source and per characteristic

#### Cluster Fusion

Clusters merge when they share identical enriched terms.

| Metric | Cluster count |
|---|---|
| Before fusion | 53 |
| After fusion | 44 |

Table 7: Summary table of the cluster count before and after cluster fusion

| old_clustr | after_GO_fusion | after_KEGG_fusion | after_WP_fusion |
|---|---|---|---|
| 2 | 2 | 1 | 1 |
| 6 | 6 | 1 | 1 |
| 8 | 8 | 8 | 4 |
| 26 | 16 | 16 | 16 |
| 30 | 13 | 13 | 13 |
| 44 | 44 | 44 | 16 |
| 45 | 45 | 10 | 10 |
| 49 | 49 | 23 | 23 |
| 53 | 53 | 14 | 14 |

Table 8: Summary table of cluster fusion process

#### Lonely fishing

Lonely genes are incorporated into clusters if they share a common term with any given cluster. The friendly limit (friendly_limit) was set to **0** .

| Metric | All | Annotated |
|---|---|---|
| Lonely genes before fishing | 1966 | 1334 |
| Lonely genes | 505 | 505 |

| Metric | All | Annotated |
|---|---|---|
| fished | | |
| Lonely genes after fishing | 1461 | 829 |

Table 9: Summary table of all and only biologically annotated genes involved in the lonely fishing

#### 4 The results as an interactive table

Show 10 entries Search:

| | transcript_id | gene_id | gene_name | NewCluster | Friendliness | Term_name | Source |
|---|---|---|---|---|---|---|---|
| | All | All | All | All | All | All | All |
| 1 | ENSDART00000000069.8 | ENSDARG00000000068 | slc9a3r1a_g2t1 | Lonely | 1 | | |
| 2 | ENSDART00000000070.7 | ENSDARG00000000069 | dap_g2t1 | Lonely | 1 | | |
| 3 | ENSDART00000001691.8 | ENSDARG00000001463 | tdh2 | Lonely | 1 | Glycine, serine and threonine metabolism | KEGG |
| 4 | ENSDART00000001691.8 | ENSDARG00000001463 | tdh2 | Lonely | 1 | Metabolic pathways | KEGG |
| 5 | ENSDART00000002053.11 | ENSDARG00000000476 | pms1_t2 | 27 | 1 | DNA metabolic process | GO:BF |
| 6 | ENSDART00000002053.11 | ENSDARG00000000476 | pms1_t2 | 27 | 1 | mismatch repair | GO:BF |
| 7 | ENSDART00000002128.7 | ENSDARG00000009215 | zgc:112437 | 47 | 1 | Cell adhesion molecules | KEGG |
| 8 | ENSDART00000002128.7 | ENSDARG00000009215 | zgc:112437 | 47 | 1 | Tight junction | KEGG |
| 9 | ENSDART00000002164.9 | ENSDARG00000008433 | unc45b_t2 | Lonely | 1 | | |
| 10 | ENSDART00000002279.8 | ENSDARG00000021442 | cdh11 | Lonely | 1 | | |

Showing 1 to 10 of 5,916 entries Previous 1 2 3 4 5 … 592 Next

Table 10: Summary table of the workflow results

#### 5 The results as sensitivity plots

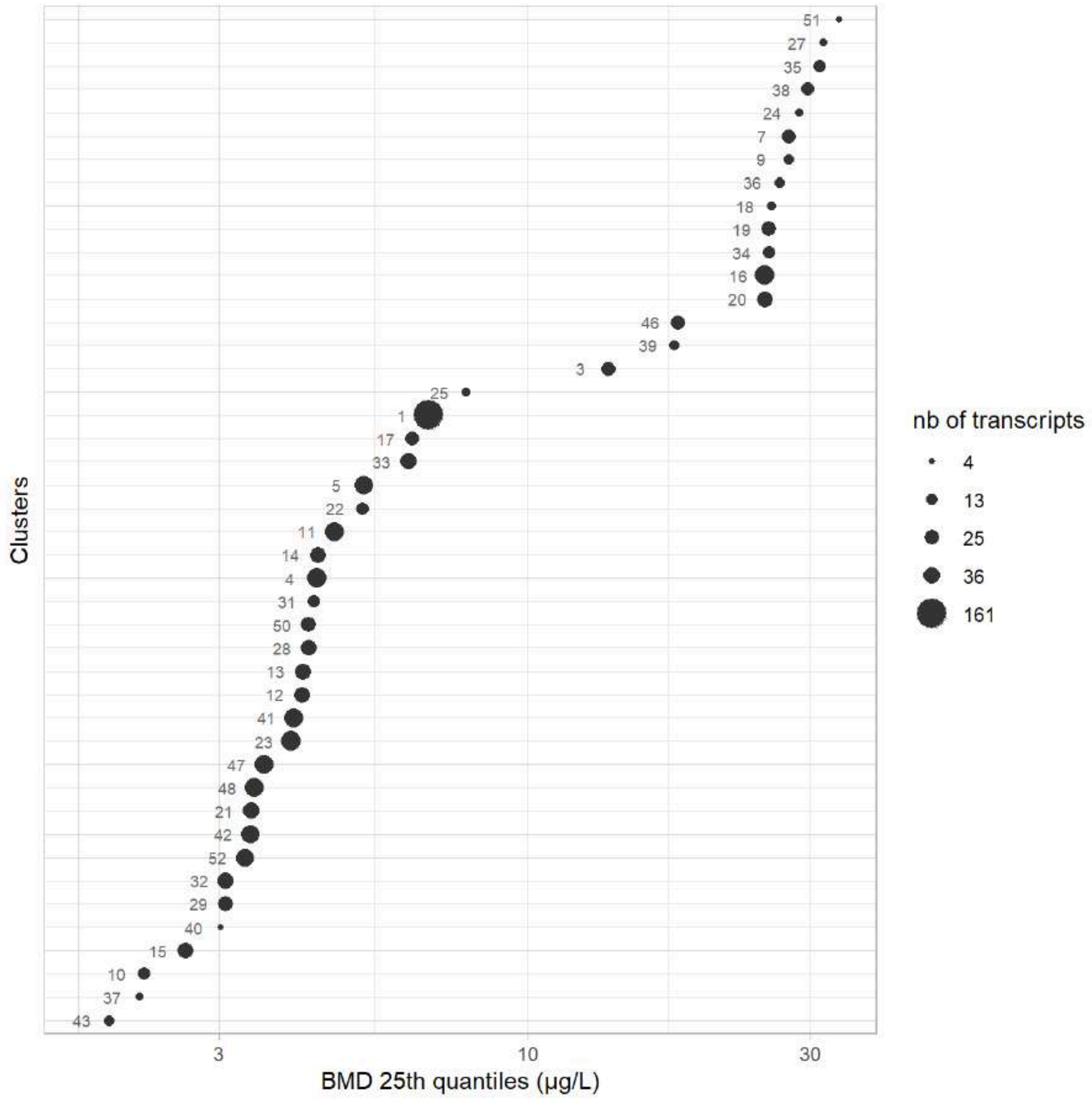


Figure 1: Sensitivity plot of clusters summarised by BMD first quartile

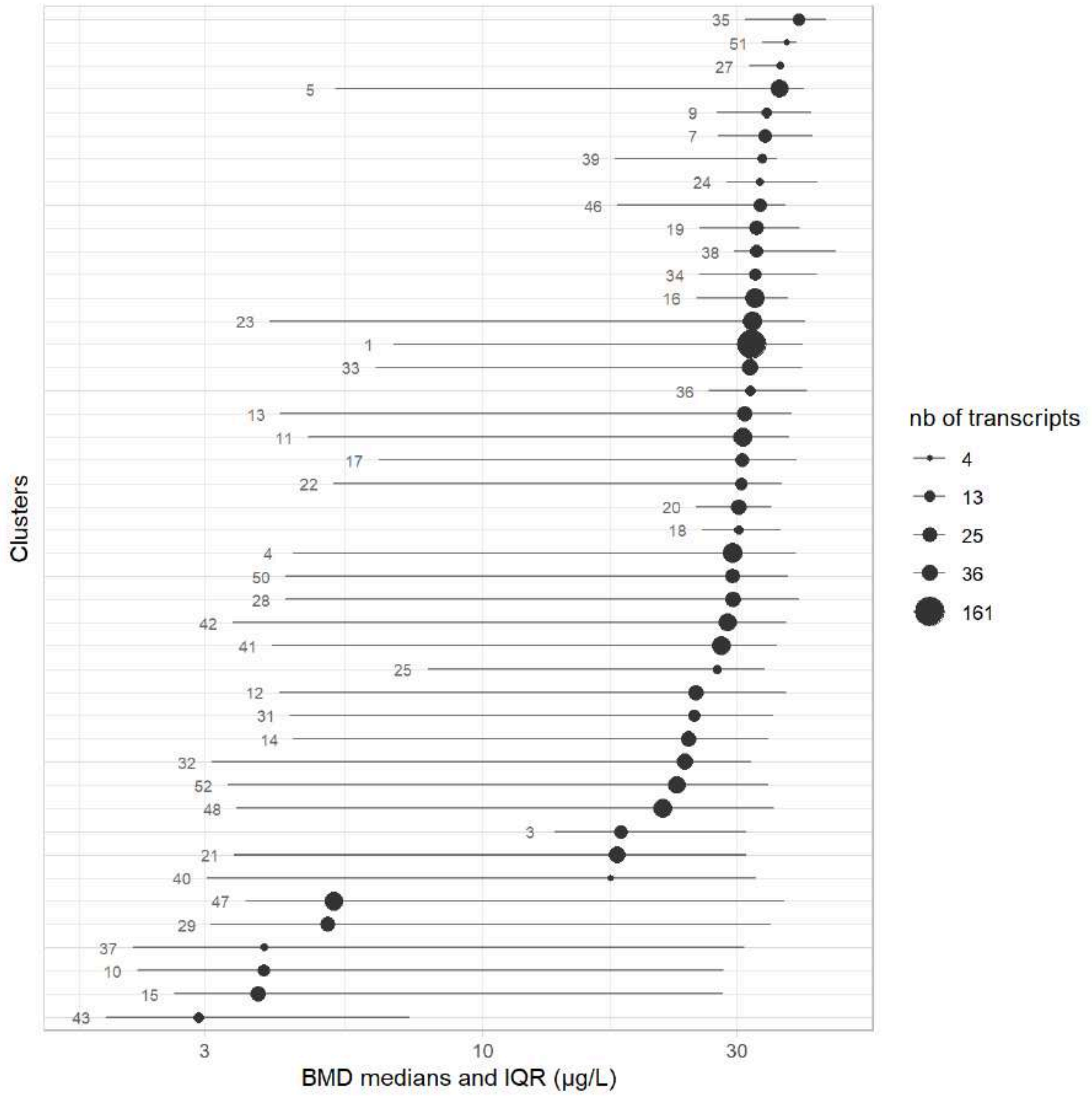


Figure 2: Sensitivity plot of clusters summarised by BMD median with the interquartile range as an interval

#### 6 The final 44 clusters from highest to lowest sensitivity

Each cluster is represented by an interactive :

- a **curvesplot** representing the fitted curves (*with x-axis the dose in log scale and the y-axis the scaled signal*)
- a **BMDplot** representing an ECDF plot with confidence intervals on each BMD value (*with the x-axis the dose in log scale and the y-axis the empirical cumulative distribution*)
- a **table** of the cluster content with some columns of interest:
  - **id**: the gene name associated to the transcript
  - **BMD.zSD**: the computed BMD for the transcript with the color of the cell associated to the BMD value, following the *viridis* color palette.
  - **TF**: the status of the transcripts' gene being either a transcription or co-transcription factor
  - **Trend**: the shape of the dose-response curve fitted
  - **Friendliness**: the total number of clusters that the transcript is associated to

### Cluster 43 - 9 transcripts (BMD q25 = 1.95 µg/L)

**Driver GO terms :**

**KEGG pathways :** Retinol metabolism

**Wikipathways :**

Curvesplot | BMDplot

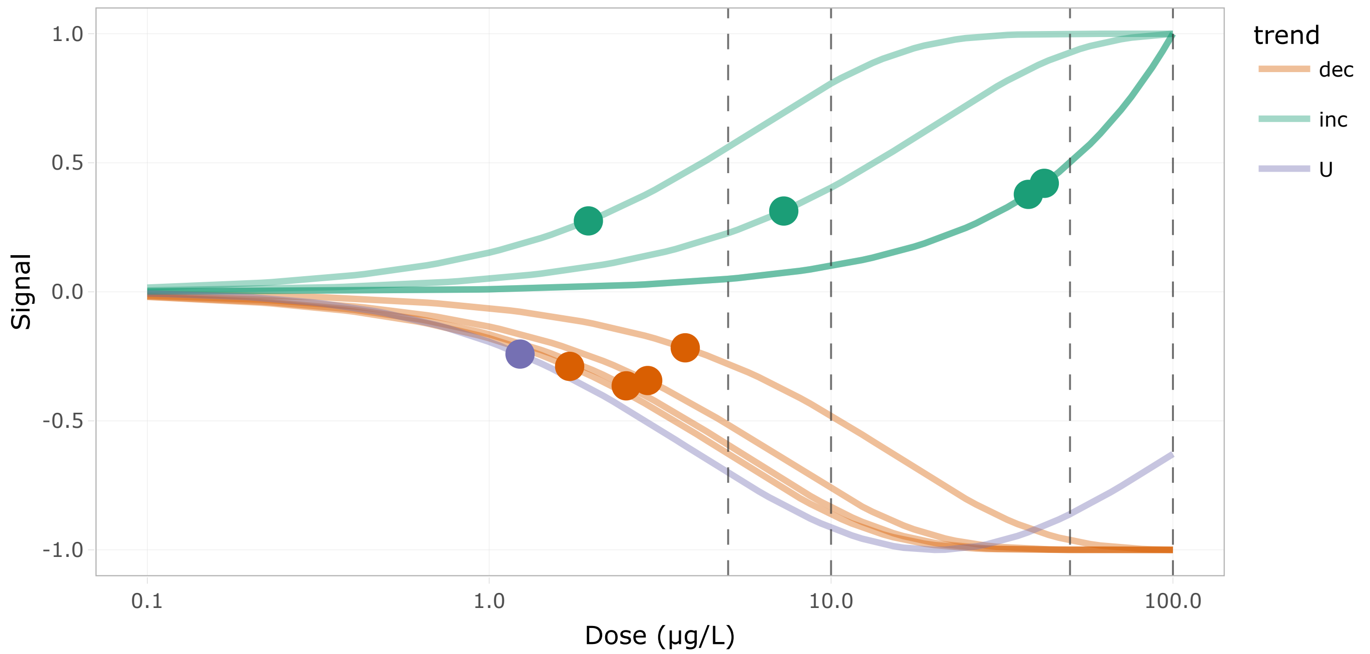


Figure 3: DR curves for the 1st cluster

Show 10 entries

Search:

| | id | BMD.zSD | TF | trend | friendliness |
|---|---|---|---|---|---|
| | All | All | All | All | All |
| 1 | cyp27c1 | 7.273 | false | inc | 1 |
| 2 | cyp3c1_t2 | 42.047 | false | inc | 1 |
| 3 | cyp3c3_g2t1 | 1.231 | false | U | 1 |
| 4 | lrata | 37.765 | false | inc | 1 |
| 5 | lratb.2 | 1.951 | false | inc | 1 |
| 6 | ugt1b1_t2 | 3.745 | false | dec | 1 |
| 7 | ugt2a7_g2t2 | 2.516 | false | dec | 1 |
| 8 | ugt2a7_g3t1 | 1.72 | false | dec | 1 |
| 9 | zgc:77938 | 2.909 | false | dec | 1 |

Showing 1 to 9 of 9 entries

Previous 1 Next

Table 11: Table of the 1st cluster content

#### Cluster 37 - 5 transcripts (BMD q25 = 2.19 µg/L)

**Driver GO terms** :

**KEGG pathways** :

**Wikipathways** :

Curvesplot | BMDplot

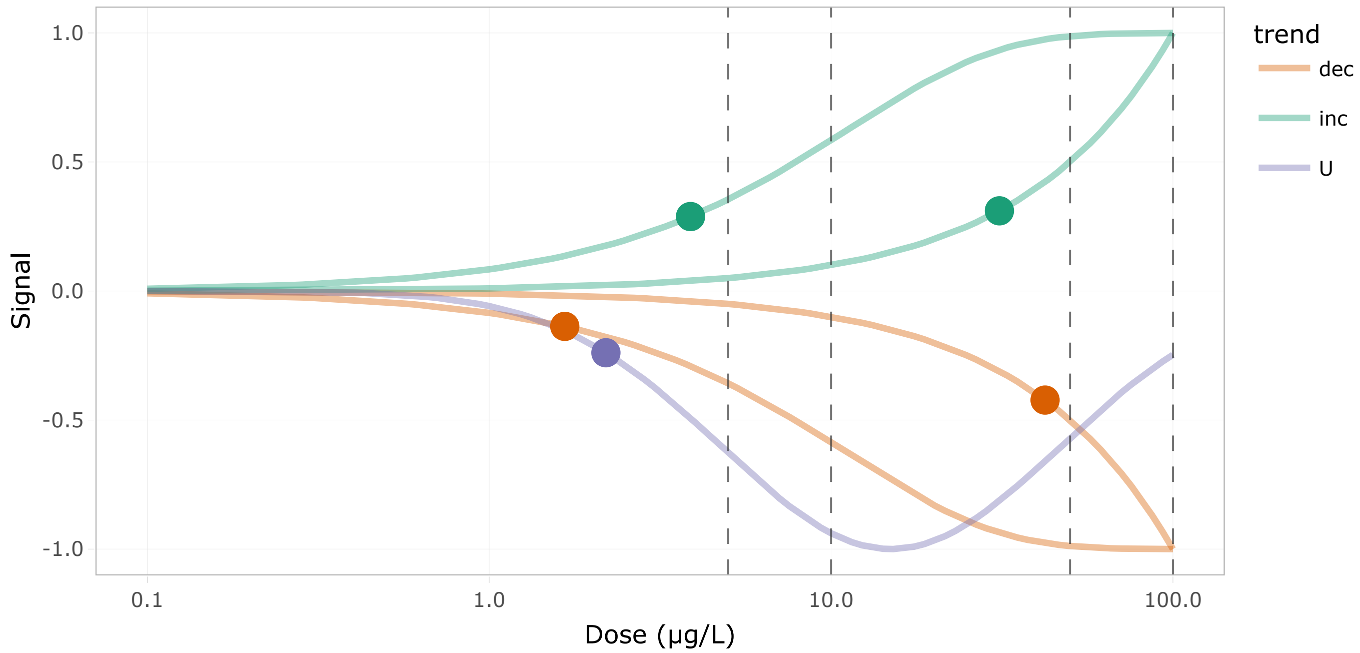


Figure 5: DR curves for the 2nd cluster

Show 10 entries

Search:

| | id | BMD.zSD | TF | trend | friendliness |
|---|---|---|---|---|---|
| | All | All | All | All | All |
| 1 | atg12_t2 | 31.073 | false | inc | 1 |
| 2 | atg3 | 2.195 | false | U | 1 |
| 3 | pi4kaa | 42.264 | false | dec | 1 |
| 4 | pik3c3_t2 | 3.881 | false | inc | 1 |
| 5 | rab5ab | 1.664 | false | dec | 1 |

Showing 1 to 5 of 5 entries

Previous 1 Next

Table 12: Table of the 2nd cluster content

#### Cluster 10 - 16 transcripts (BMD q25 = 2.23 µg/L)

**Driver GO terms** : sphingolipid metabolic process

**KEGG pathways** : Sphingolipid metabolism

**Wikipathways :**

Curvesplot BMDplot

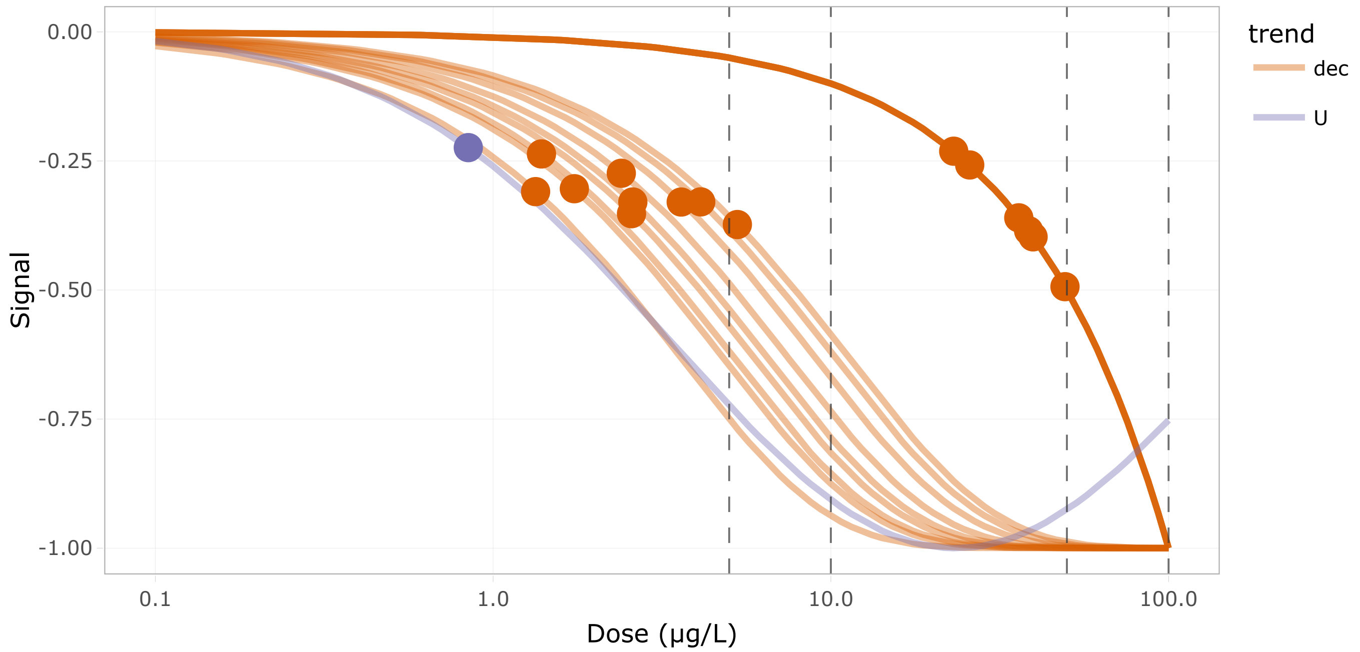


Figure 7: DR curves for the 3rd cluster

Show 10 entries Search:

| | id | BMD.zSD | TF | trend | friendliness |
|---|---|---|---|---|---|
| | All | All | All | All | All |
| 1 | SPHK1 | 36.011 | false | dec | 1 |
| 2 | asah2 | 1.336 | false | dec | 1 |
| 3 | b4galt6 | 38.515 | false | dec | 1 |
| 4 | cers2a_t2 | 25.775 | false | dec | 1 |
| 5 | cers3a_t2 | 1.74 | false | dec | 1 |
| 6 | cers5 | 2.592 | false | dec | 2 |
| 7 | enpp7.1_g3t1 | 0.844 | false | U | 1 |
| 8 | gba2_t2 | 1.39 | false | dec | 1 |
| 9 | hexb_t3 | 39.699 | false | dec | 3 |
| 10 | kdsr_t2 | 2.396 | false | dec | 1 |

Showing 1 to 10 of 16 entries Previous 1 2 Next

Table 13: Table of the 3rd cluster content

#### Cluster 15 - 31 transcripts (BMD q25 = 2.62 µg/L)

**Driver GO terms** : lipid biosynthetic process

**KEGG pathways** : Steroid biosynthesis

**Wikipathways** : Cholesterol biosynthesis

Curvesplot BMDplot

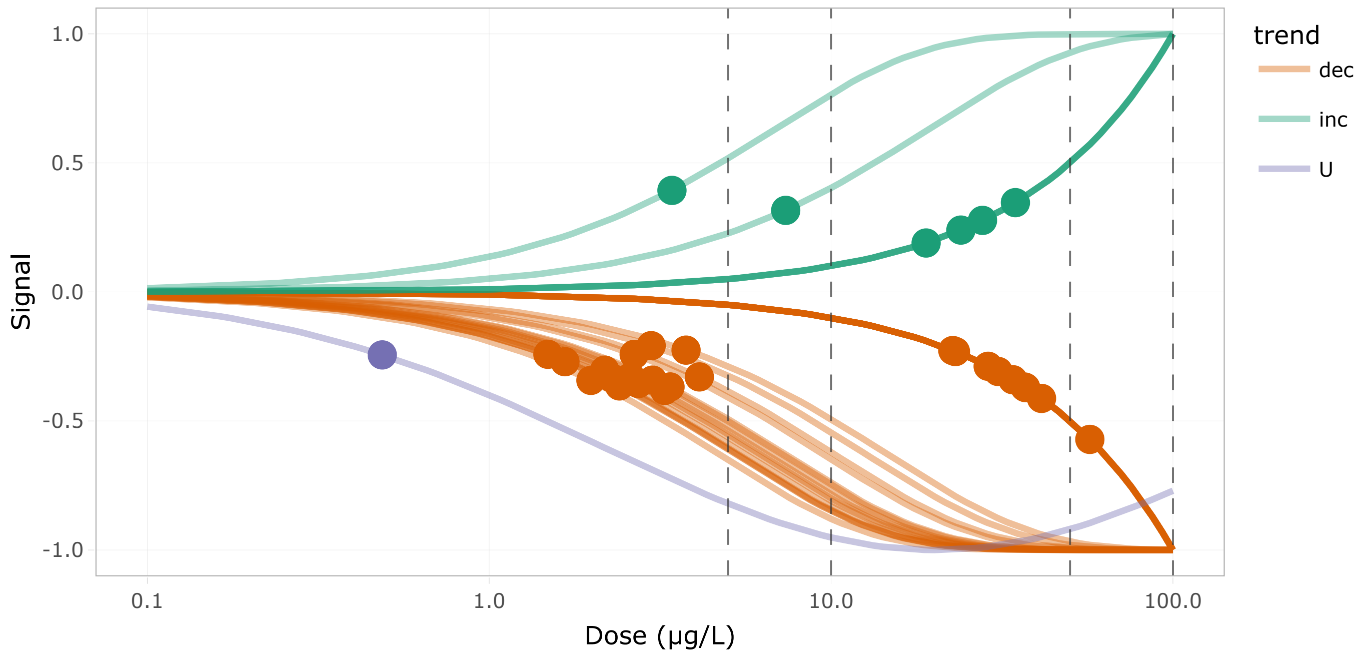


Figure 9: DR curves for the 4th cluster

Show 10 entries Search:

| | id | BMD.zSD | TF | trend | friendliness |
|---|---|---|---|---|---|
| | All | All | All | All | All |
| 1 | ERG28 | 41.262 | false | dec | 1 |
| 2 | PYURF_t1 | 7.369 | false | inc | 1 |
| 3 | acsl1b | 2.177 | false | dec | 3 |
| 4 | alox12 | 3.381 | false | dec | 3 |
| 5 | cers5 | 2.592 | false | dec | 2 |
| 6 | crppa | 18.967 | false | inc | 2 |
| 7 | cyp17a1 | 2.731 | false | dec | 1 |
| 8 | cyp27b1 | 3.005 | false | dec | 1 |
| 9 | dhcr7 | 2.405 | false | dec | 1 |
| 10 | fa2h_t3 | 1.666 | false | dec | 3 |

Showing 1 to 10 of 31 entries Previous 1 2 3 4 Next

Table 14: Table of the 4th cluster content

#### Cluster 40 - 4 transcripts (BMD q25 = 3.01 µg/L)

**Driver GO terms** :

**KEGG pathways :**

**Wikipathways :**

Curvesplot BMDplot

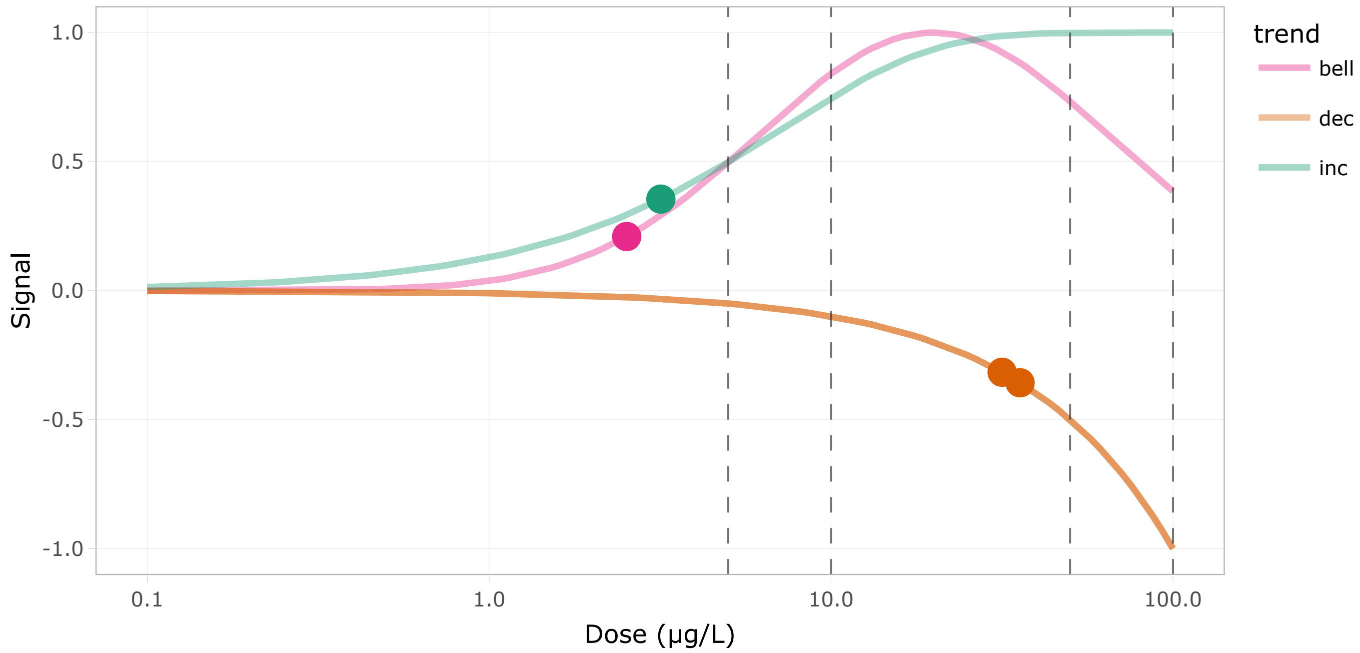


Figure 11: DR curves for the 5th cluster

Show 10 entries Search:

| | id | BMD.zSD | TF | trend | friendliness |
|---|---|---|---|---|---|
| | All | All | All | All | All |
| 1 | cbfa2t3_t2 | 2.525 | false | bell | 1 |
| 2 | ldb1b_t2 | 31.648 | true | dec | 1 |
| 3 | lmo1_t1 | 3.178 | true | inc | 1 |
| 4 | tcf3a_t2 | 35.716 | true | dec | 1 |

Showing 1 to 4 of 4 entries Previous 1 Next

Table 15: Table of the 5th cluster content

### Cluster 29 - 27 transcripts (BMD q25 = 3.07 µg/L)

**Driver GO terms :** myofibril assembly, striated muscle contraction

**KEGG pathways :** Motor proteins

**Wikipathways :**

Curvesplot BMDplot

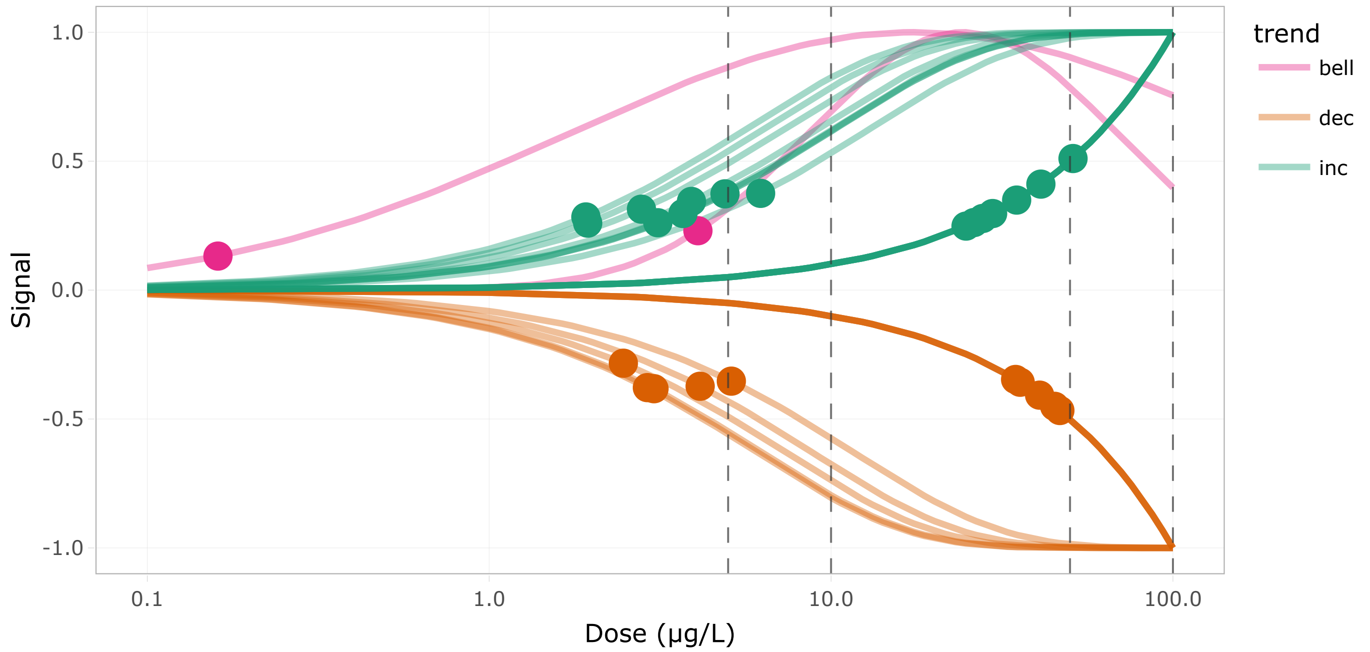

Figure 13: DR curves for the 6th cluster

Show 10 entries Search:

| | id | BMD.zSD | TF | trend | friendliness |
|---|---|---|---|---|---|
| | All | All | All | All | All |
| 1 | MYO1D | 4.141 | false | dec | 1 |
| 2 | chchd10_t2 | 41.092 | false | inc | 1 |
| 3 | cx39.9 | 27.779 | false | inc | 2 |
| 4 | dctn1a_t2 | 35.737 | false | dec | 2 |
| 5 | dctn4_t3 | 4.079 | false | bell | 2 |
| 6 | dynll2b | 3.689 | false | inc | 3 |
| 7 | erp44 | 3.034 | false | dec | 3 |
| 8 | kif1b_t1 | 5.109 | false | dec | 3 |
| 9 | kifc1_t2 | 26.202 | false | inc | 2 |
| 10 | kifc3_t5 | 45.079 | false | dec | 2 |

Showing 1 to 10 of 27 entries Previous 1 2 3 Next

Table 16: Table of the 6th cluster content

#### Cluster 32 - 33 transcripts (BMD q25 = 3.08 µg/L)

**Driver GO terms** : carboxylic acid metabolic process

**KEGG pathways** : Alanine, aspartate and glutamate metabolism, Biosynthesis of amino acids, Cysteine and methionine metabolism

**Wikipathways** :

Curvesplot BMDplot

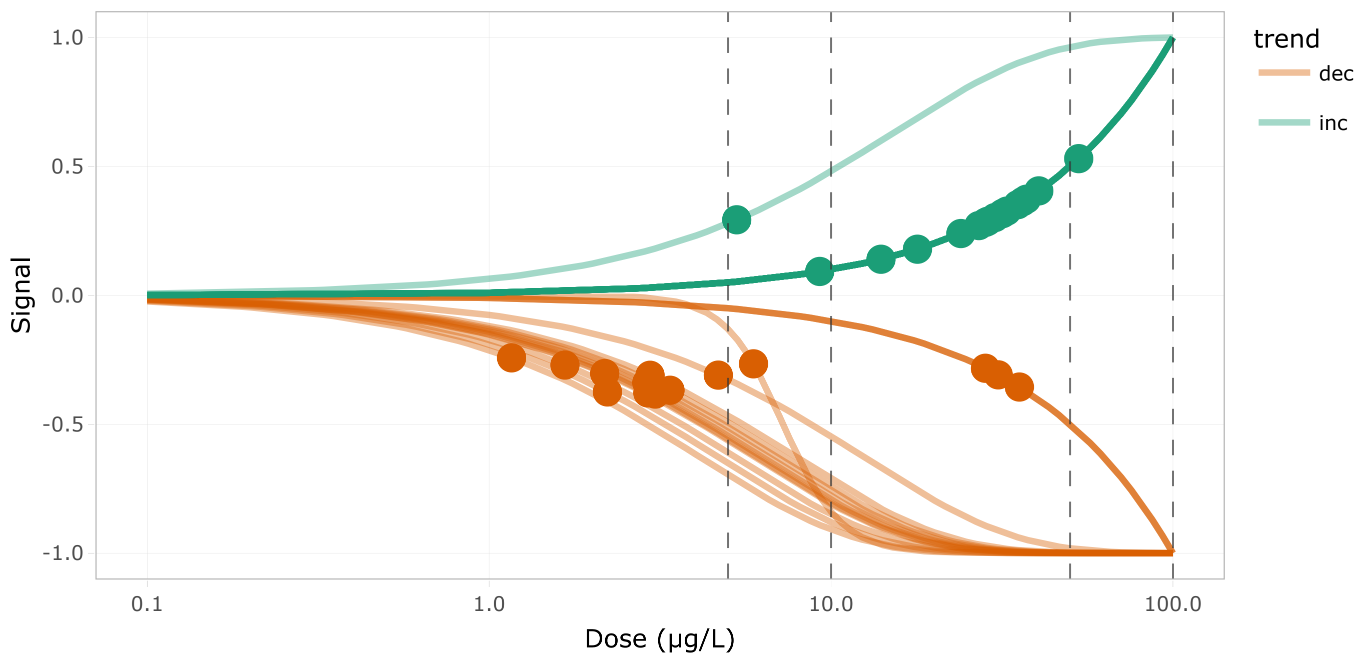


Figure 15: DR curves for the 7th cluster

Show 10 entries Search:

| | id | BMD.zSD | TF | trend | friendliness |
|---|---|---|---|---|---|
| | All | All | All | All | All |
| 1 | HPDL | 28.323 | false | inc | 2 |
| 2 | abcd1_t2 | 4.68 | false | dec | 2 |
| 3 | acox3 | 3.06 | false | dec | 2 |
| 4 | acsl1b | 2.177 | false | dec | 3 |
| 5 | adi1 | 28.749 | false | inc | 3 |
| 6 | aldh1l1_g2t1 | 2.912 | false | dec | 2 |
| 7 | alox12 | 3.381 | false | dec | 3 |
| 8 | apip | 37.287 | false | inc | 2 |
| 9 | asdurf_t2 | 40.519 | false | inc | 2 |
| 10 | aspa_t2 | 28.279 | false | dec | 1 |

Showing 1 to 10 of 33 entries

Previous 1 2 3 4 Next

Table 17: Table of the 7th cluster content

#### Cluster 52 - 37 transcripts (BMD q25 = 3.31 μg/L)

**Driver GO terms** : protein glycosylation

**KEGG pathways** : N-Glycan biosynthesis, Protein processing in endoplasmic reticulum, Various types of N-glycan biosynthesis

**Wikipathways** :

Curvesplot BMDplot

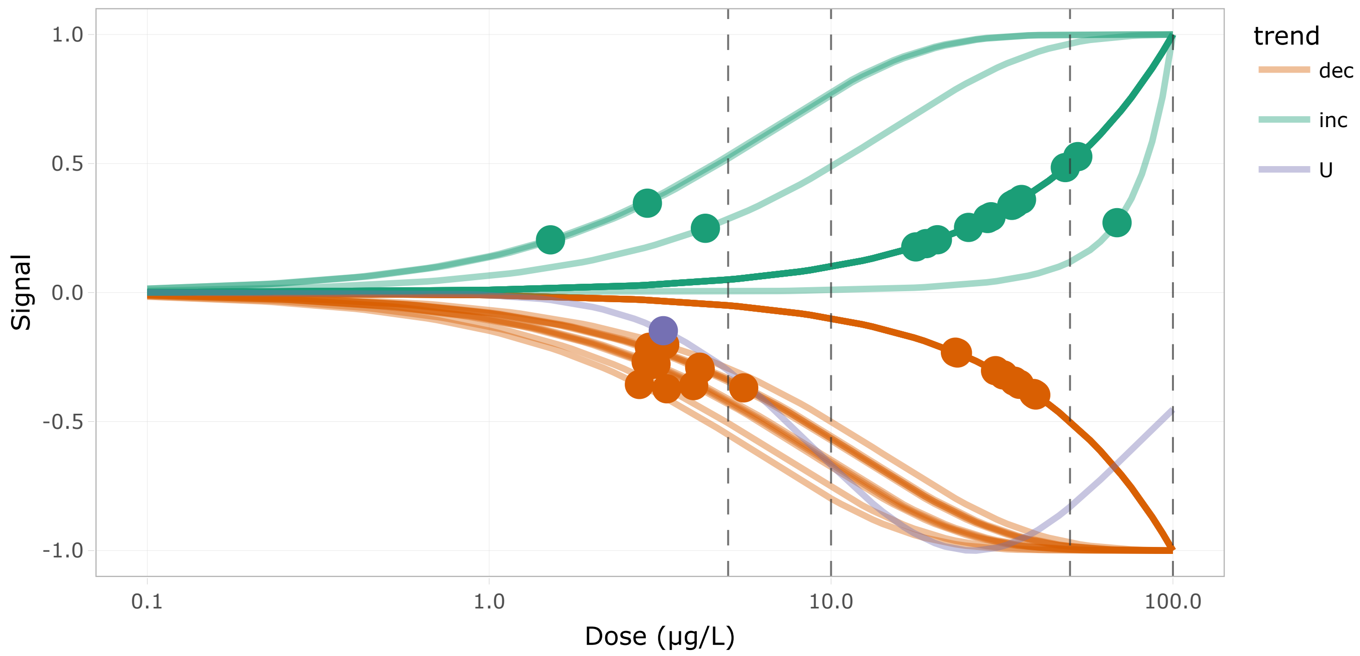


Figure 17: DR curves for the 8th cluster

Show 10 entries

Search:

| | id | BMD.zSD | TF | trend | friendliness |
|---|---|---|---|---|---|
| | All | All | All | All | All |
| 1 | RNF5 | 31.927 | false | dec | 4 |
| 2 | alg11_t2 | 23.422 | false | dec | 1 |
| 3 | atf6 | 30.276 | true | dec | 4 |
| 4 | calr3b | 4.129 | false | dec | 6 |
| 5 | canx | 4.142 | false | dec | 5 |
| 6 | crppa | 18.967 | false | inc | 2 |
| 7 | dnajb11 | 2.879 | false | dec | 5 |
| 8 | dnajc1 | 33.784 | true | inc | 4 |
| 9 | dnajc5aa | 39.493 | false | dec | 5 |
| 10 | eif2s1b | 34.593 | false | inc | 8 |

Showing 1 to 10 of 37 entries

Previous 1 2 3 4 Next

Table 18: Table of the 8th cluster content

#### Cluster 42 - 43 transcripts (BMD q25 = 3.38 µg/L)

**Driver GO terms** : canonical NF-kappaB signal transduction

**KEGG pathways** : Apoptosis, C-type lectin receptor signaling pathway, MAPK signaling pathway

**Wikipathways :**

Curvesplot | BMDplot

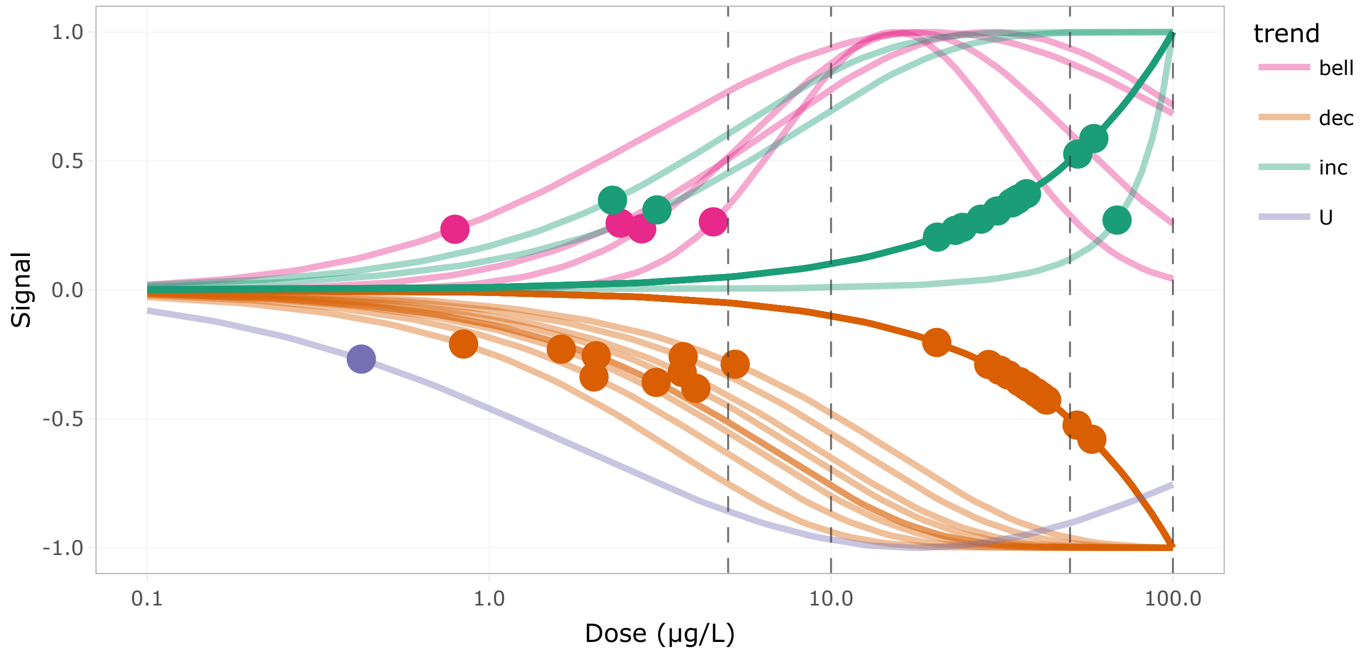


Figure 19: DR curves for the 9th cluster

Show 10 entries

Search:

| | id | BMD.zSD | TF | trend | friendliness |
|---|---|---|---|---|---|
| | All | All | All | All | All |
| 1 | akt3a_t2 | 37.907 | false | dec | 3 |
| 2 | apaf1_t2 | 0.423 | false | U | 2 |
| 3 | arf2b_g1t2 | 3.693 | false | dec | 1 |
| 4 | braf_t2 | 2.794 | false | bell | 4 |
| 5 | cacng1a | 30.612 | false | inc | 2 |
| 6 | cts12 | 52.433 | false | dec | 1 |
| 7 | ctsh | 33.747 | false | inc | 1 |
| 8 | ctss2.2 | 58.704 | false | inc | 1 |
| 9 | cylda_t2 | 2.295 | false | inc | 3 |
| 10 | eif2s1b | 34.593 | false | inc | 8 |

Showing 1 to 10 of 43 entries

Previous 1 2 3 4 5 Next

Table 19: Table of the 9th cluster content

### Cluster 21 - 35 transcripts (BMD q25 = 3.4 µg/L)

**Driver GO terms** : organic acid metabolic process

**KEGG pathways** : 2-Oxocarboxylic acid metabolism, Biosynthesis of amino acids, Carbon metabolism, Citrate cycle (TCA cycle), Glyoxylate and dicarboxylate metabolism

**Wikipathways** : TCA cycle

Curvesplot BMDplot

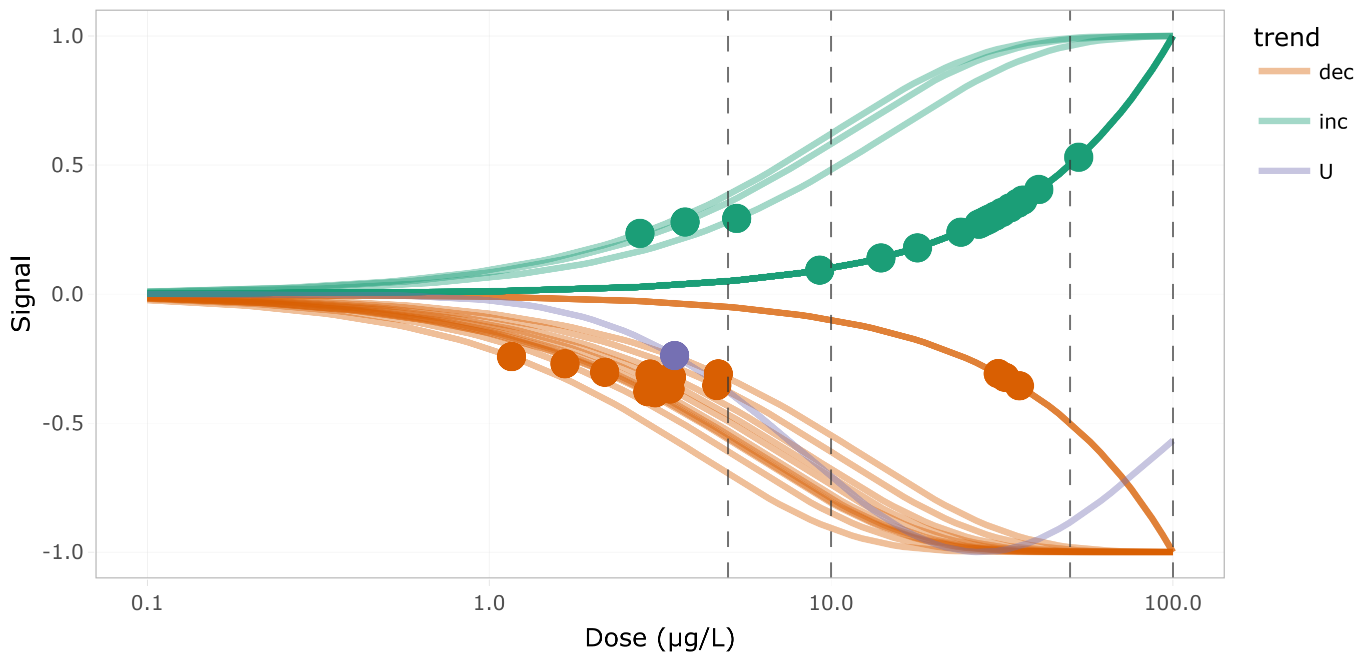


Figure 21: DR curves for the 10th cluster

Show 10 entries Search:

| | id | BMD.zSD | TF | trend | friendliness |
|---|---|---|---|---|---|
| | All | All | All | All | All |
| 1 | HPDL | 28.323 | false | inc | 2 |
| 2 | abcd1_t2 | 4.68 | false | dec | 2 |
| 3 | aco1 | 3.412 | false | dec | 1 |
| 4 | acox3 | 3.06 | false | dec | 2 |
| 5 | acsl1b | 2.177 | false | dec | 3 |
| 6 | acss2l | 27.531 | false | inc | 1 |
| 7 | adi1 | 28.749 | false | inc | 3 |
| 8 | aldh1l1_g2t1 | 2.912 | false | dec | 2 |
| 9 | alox12 | 3.381 | false | dec | 3 |
| 10 | asdurf_t2 | 40.519 | false | inc | 2 |

Showing 1 to 10 of 35 entries Previous 1 2 3 4 Next

Table 20: Table of the 10th cluster content

### Cluster 48 - 50 transcripts (BMD q25 = 3.44 μg/L)

**Driver GO terms** : carbohydrate metabolic process

**KEGG pathways** : N-Glycan biosynthesis, Protein processing in endoplasmic reticulum

**Wikipathways** :

Curvesplot BMDplot

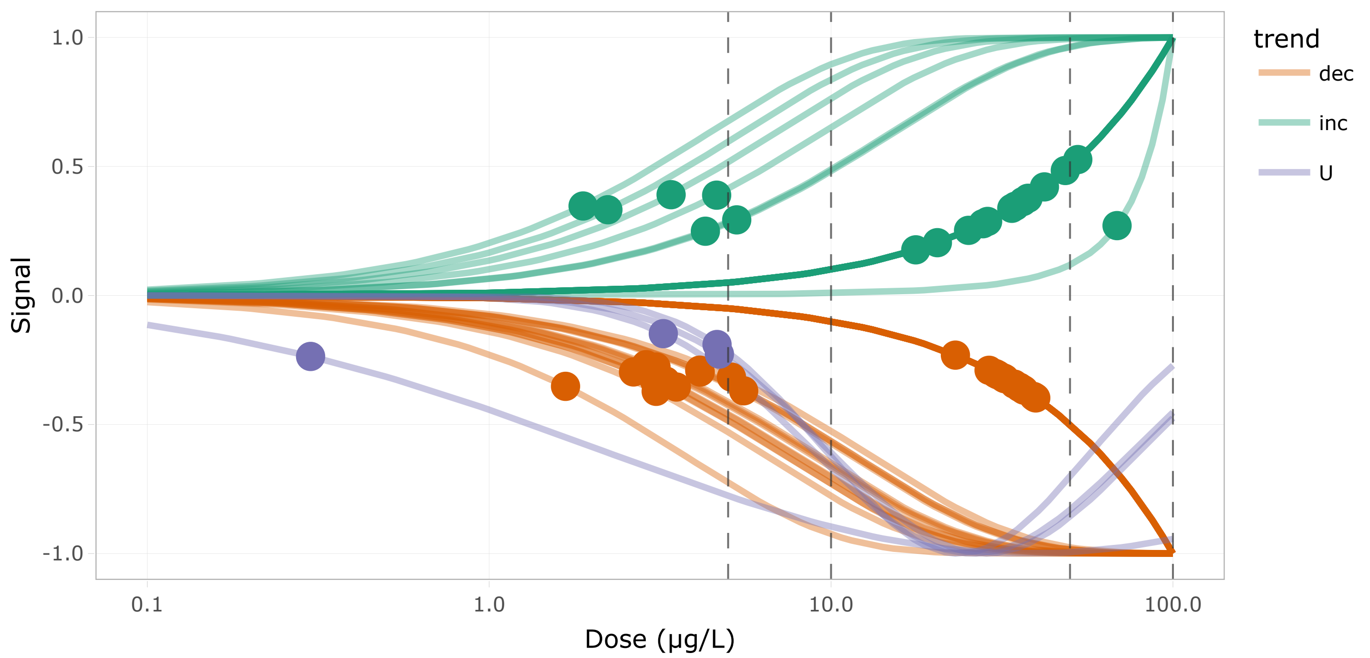


Figure 23: DR curves for the 11th cluster

Show 10 entries Search:

| | id | BMD.zSD | TF | trend | friendliness |
|---|---|---|---|---|---|
| | All | All | All | All | All |
| 1 | CABZ01079192.1 | 1.672 | false | dec | 1 |
| 2 | CR759923.1_t4 | 2.225 | false | inc | 1 |
| 3 | CR774178.2 | 5.113 | false | dec | 1 |
| 4 | GANAB_t2 | 2.643 | false | dec | 1 |
| 5 | MAN1C1_t2 | 28.988 | false | dec | 1 |
| 6 | RNF5 | 31.927 | false | dec | 4 |
| 7 | agla_t1 | 3.046 | false | dec | 2 |
| 8 | atf6 | 30.276 | true | dec | 4 |
| 9 | calr3b | 4.129 | false | dec | 6 |
| 10 | canx | 4.142 | false | dec | 5 |

Showing 1 to 10 of 50 entries

Previous 1 2 3 4 5 Next

Table 21: Table of the 11th cluster content

#### Cluster 47 - 52 transcripts (BMD q25 = 3.57 µg/L)

**Driver GO terms** :

**KEGG pathways** : Regulation of actin cytoskeleton, Salmonella infection, Tight junction

**Wikipathways** :

Curvesplot | BMDplot

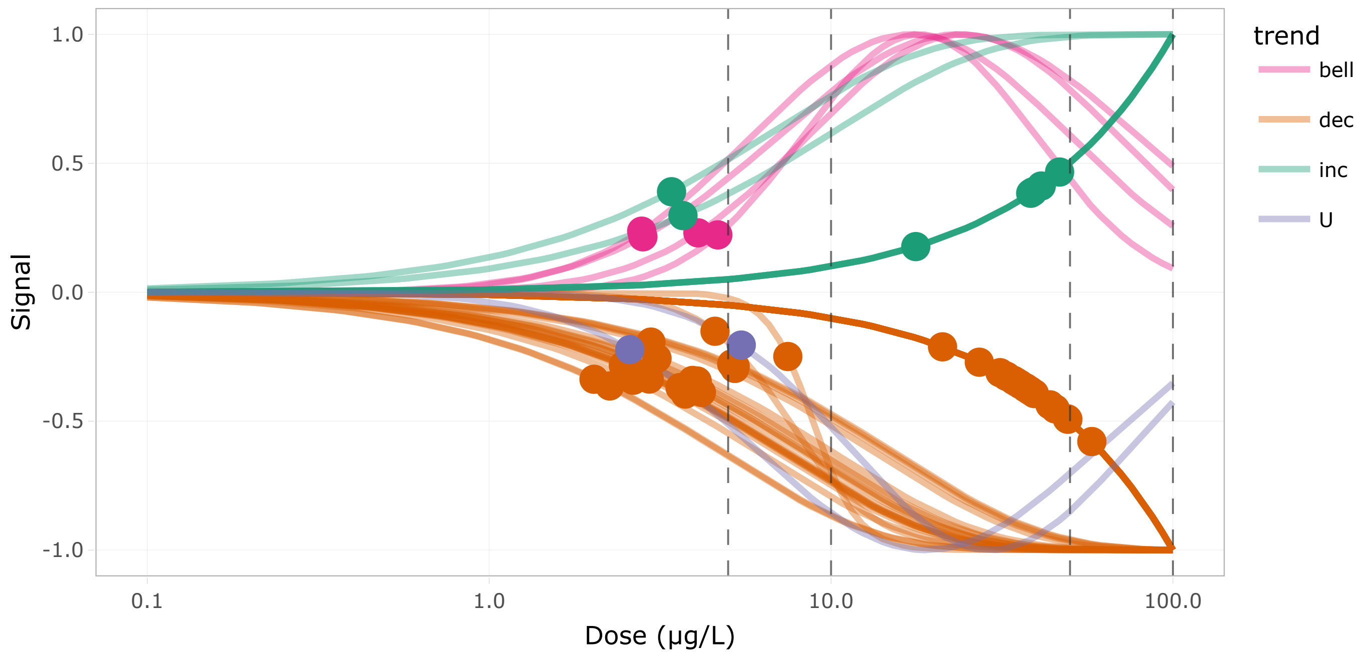


Figure 25: DR curves for the 12th cluster

Show 10 entries

Search:

| | id | BMD.zSD | TF | trend | friendliness |
|---|---|---|---|---|---|
| | All | All | All | All | All |
| 1 | ACBD3 | 2.638 | false | dec | 1 |
| 2 | abi1b_t2 | 49.282 | false | dec | 1 |
| 3 | acbd3 | 27.141 | false | dec | 1 |
| 4 | actb2_t2 | 3.748 | true | dec | 1 |
| 5 | actr2a | 4.577 | false | dec | 1 |
| 6 | actr2b | 5.231 | false | dec | 1 |
| 7 | afdna_t1 | 45.241 | false | dec | 2 |
| 8 | akt3a_t2 | 37.907 | false | dec | 3 |
| 9 | arf2a_t1 | 2.579 | false | U | 1 |
| 10 | arf2a_t2 | 2.973 | false | dec | 1 |

Showing 1 to 10 of 52 entries

Previous 1 2 3 4 5 6 Next

Table 22: Table of the 12th cluster content

### Cluster 23 - 56 transcripts (BMD q25 = 3.96 µg/L)

**Driver GO terms** : RNA splicing, via transesterification reactions, RNA splicing

**KEGG pathways** : Spliceosome

**Wikipathways** :

Curvesplot BMDplot

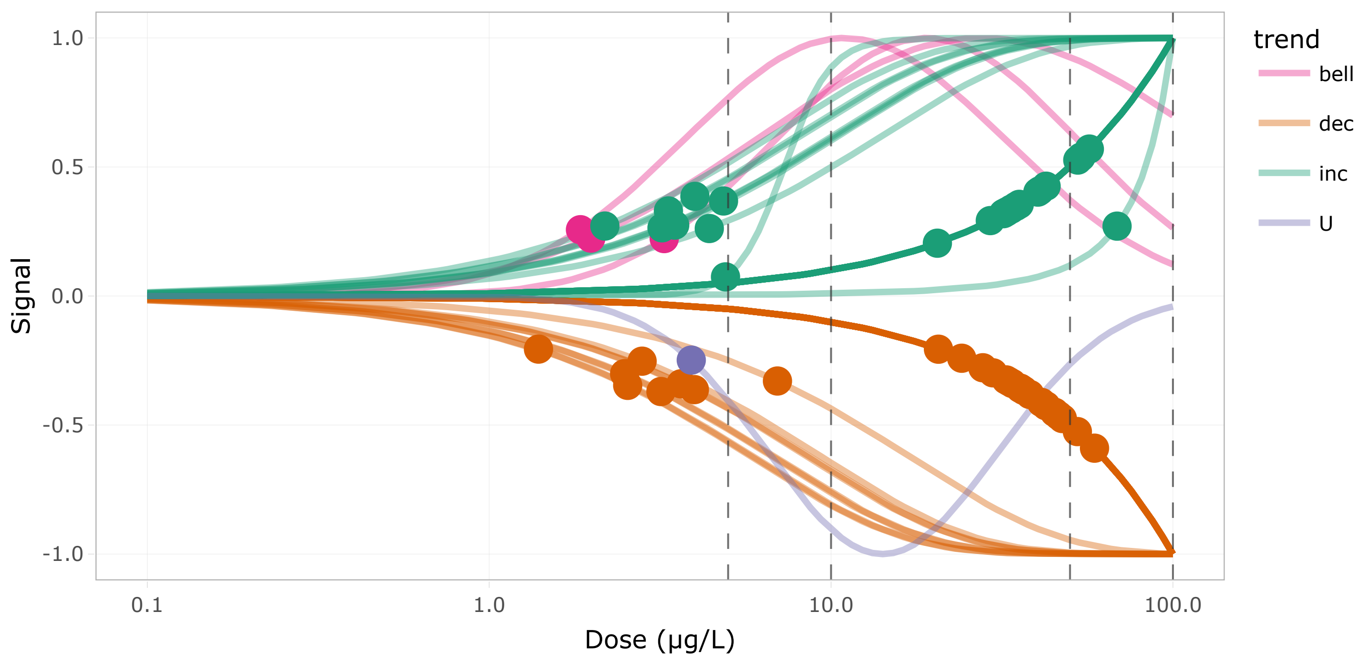


Figure 27: DR curves for the 13th cluster

Show 10 entries Search:

| | id | BMD.zSD | TF | trend | friendliness |
|---|---|---|---|---|---|
| | All | All | All | All | All |
| 1 | LSM2_t4 | 3.18 | false | dec | 1 |
| 2 | clk2a_t1 | 6.97 | false | dec | 2 |
| 3 | ddx5_t1 | 2.49 | true | dec | 1 |
| 4 | ddx5_t4 | 3.998 | true | inc | 1 |
| 5 | gemin2 | 32.258 | false | inc | 3 |
| 6 | gemin8_g2t1 | 42.542 | false | dec | 3 |
| 7 | hmga1a_t1 | 3.64 | true | dec | 1 |
| 8 | hmga1a_t2 | 2.801 | true | dec | 1 |
| 9 | hmga1a_t3 | 4.912 | true | inc | 1 |
| 10 | hnrnpa1b_t2 | 2.179 | false | inc | 2 |

Showing 1 to 10 of 56 entries

Previous 1 2 3 4 5 6 Next

Table 23: Table of the 13th cluster content

#### Cluster 41 - 51 transcripts (BMD q25 = 4.01 µg/L)

**Driver GO terms** : mRNA processing

**KEGG pathways** : mRNA surveillance pathway

**Wikipathways** : mRNA processing

Curvesplot BMDplot

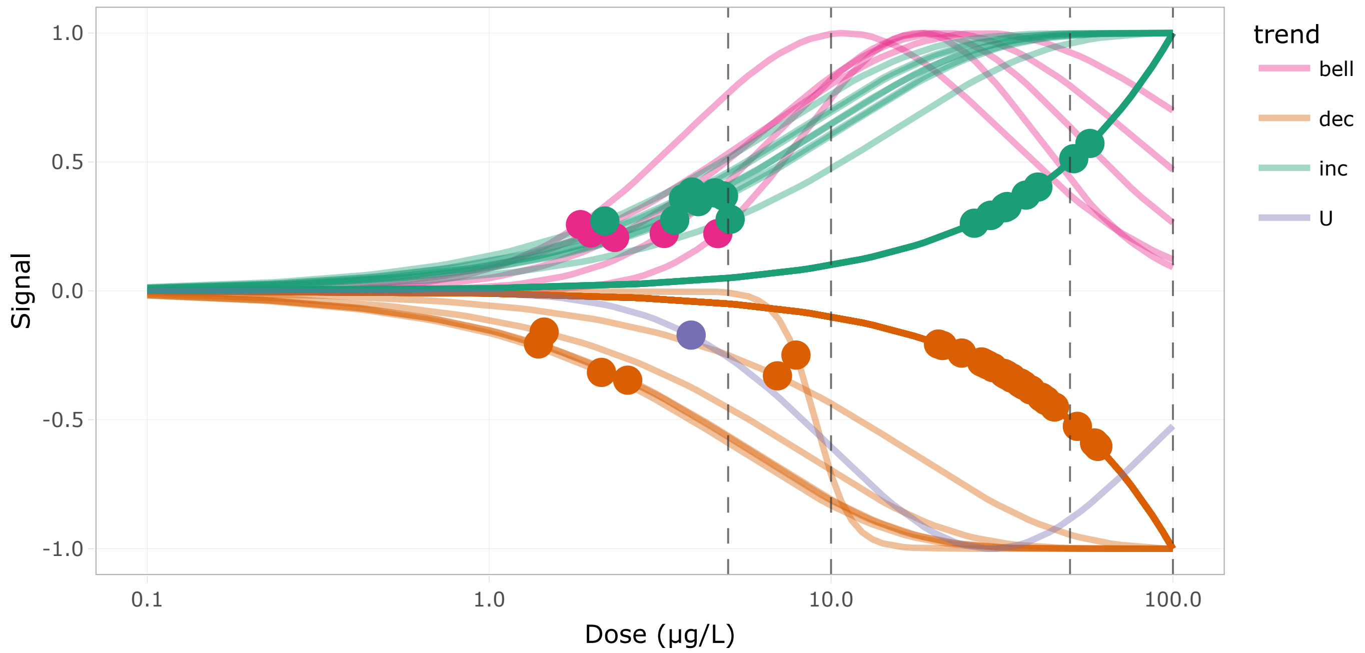


Figure 29: DR curves for the 14th cluster

Show 10 entries Search:

| | id | BMD.zSD | TF | trend | friendliness |
|---|---|---|---|---|---|
| | All | All | All | All | All |
| 1 | cdc73_t2 | 3.897 | true | U | 2 |
| 2 | celf2_t2 | 32.673 | false | dec | 1 |
| 3 | celf2_t4 | 27.548 | false | dec | 1 |
| 4 | clk2a_t1 | 6.97 | false | dec | 2 |
| 5 | cpsf3 | 57.18 | false | inc | 1 |
| 6 | cpsf6_t2 | 2.13 | false | dec | 1 |
| 7 | cstf2_t2 | 3.702 | false | inc | 1 |
| 8 | ern1 | 60.306 | true | dec | 1 |
| 9 | gemin2 | 32.258 | false | inc | 3 |
| 10 | gemin8_g2t1 | 42.542 | false | dec | 3 |

Showing 1 to 10 of 51 entries

Previous 1 2 3 4 5 6 Next

Table 24: Table of the 14th cluster content

#### Cluster 12 - 30 transcripts (BMD q25 = 4.13 µg/L)

**Driver GO terms** : protein folding

**KEGG pathways** :

**Wikipathways** :

Curvesplot BMDplot

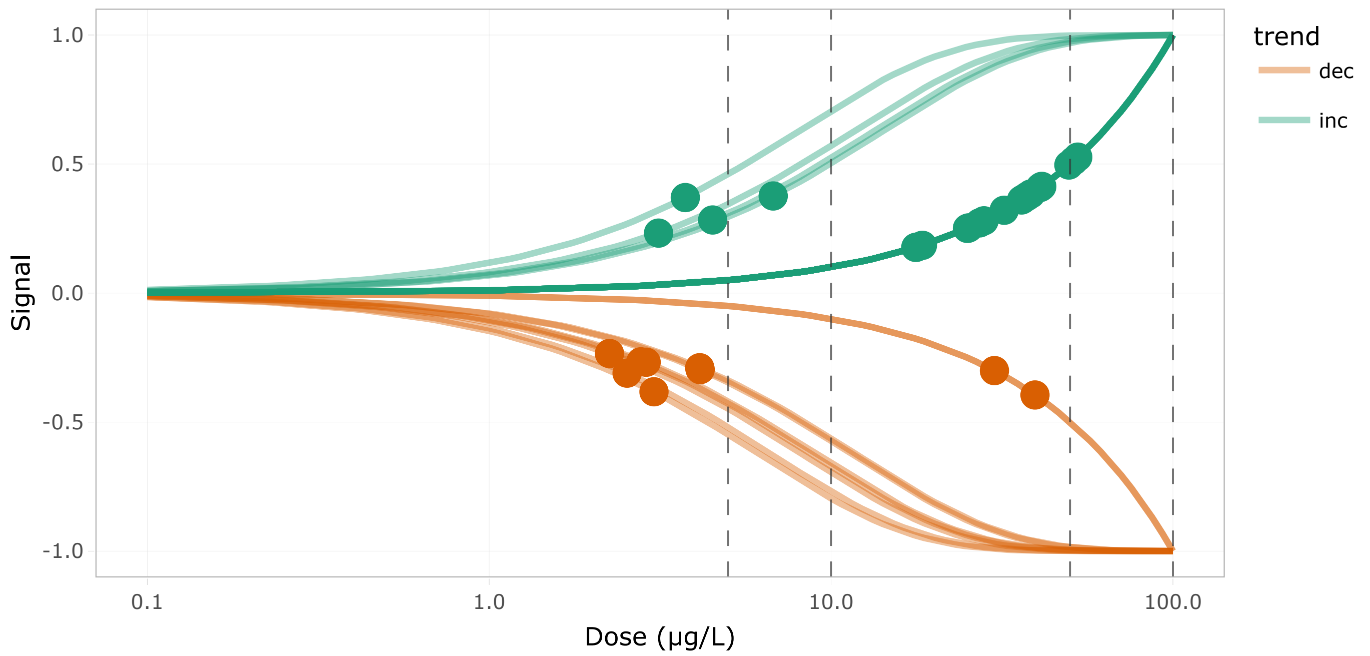


Figure 31: DR curves for the 15th cluster

Show 10 entries

Search:

| | id | BMD.zSD | TF | trend | friendliness |
|---|---|---|---|---|---|
| | All | All | All | All | All |
| 1 | CABZ01080568.1 | 17.799 | false | inc | 1 |
| 2 | ahsa1a_t3 | 4.506 | false | inc | 2 |
| 3 | calr3b | 4.129 | false | dec | 6 |
| 4 | canx | 4.142 | false | dec | 5 |
| 5 | cct2_g1t1 | 6.773 | false | inc | 1 |
| 6 | cct3 | 36.076 | false | inc | 1 |
| 7 | cct4 | 36.427 | false | inc | 1 |
| 8 | cct5 | 38.401 | false | inc | 1 |
| 9 | cct6a_t3 | 37.452 | false | inc | 1 |
| 10 | cct7 | 49.62 | false | inc | 1 |

Showing 1 to 10 of 30 entries

Previous 1 2 3 Next

Table 25: Table of the 15th cluster content

#### Cluster 13 - 28 transcripts (BMD q25 = 4.15 µg/L)

**Driver GO terms** : translational initiation

**KEGG pathways** : Herpes simplex virus 1 infection

**Wikipathways** :

Curvesplot BMDplot

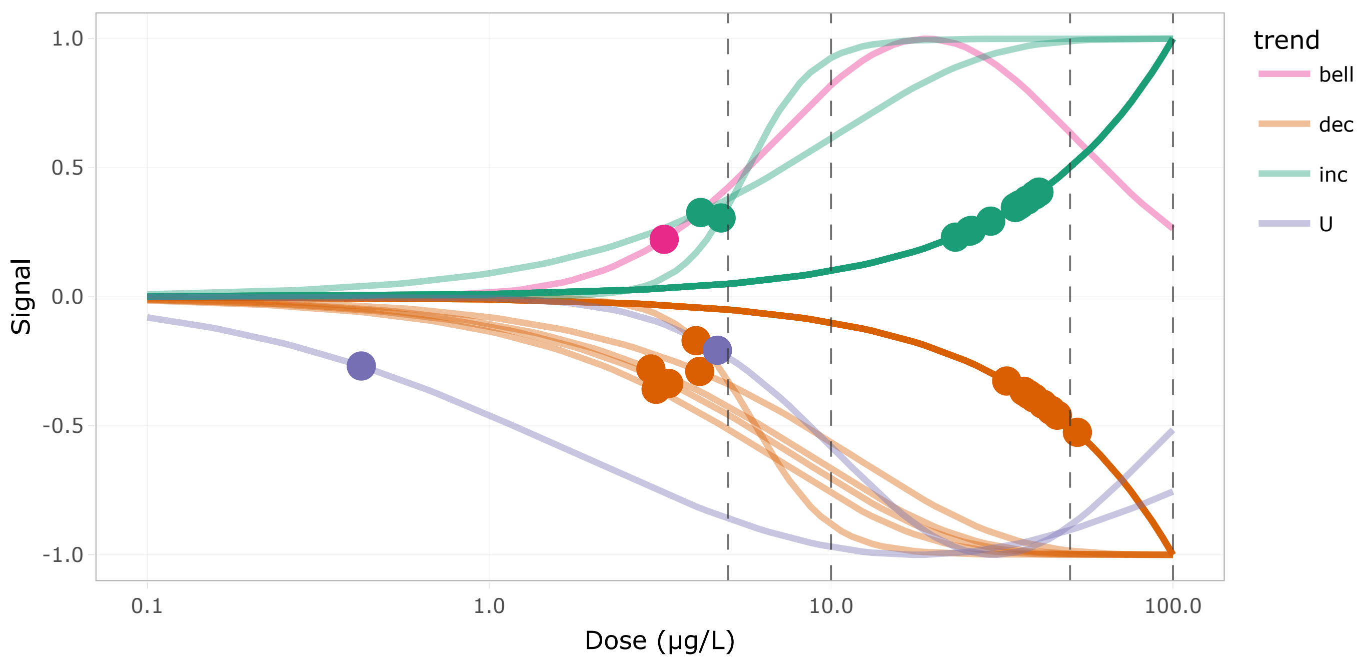


Figure 33: DR curves for the 16th cluster

Show 10 entries

Search:

| | id | BMD.zSD | TF | trend | friendliness |
|---|---|---|---|---|---|
| | All | All | All | All | All |
| 1 | akt3a_t2 | 37.907 | false | dec | 3 |
| 2 | apaf1_t2 | 0.423 | false | U | 2 |
| 3 | calr3b | 4.129 | false | dec | 6 |
| 4 | cdc123_t2 | 34.853 | false | inc | 1 |
| 5 | eif2b1_t2 | 25.738 | false | inc | 1 |
| 6 | eif2b3_t2 | 37.461 | false | inc | 1 |
| 7 | eif2b4_t2 | 25.392 | false | inc | 1 |
| 8 | eif2b5 | 39.412 | false | inc | 1 |
| 9 | eif2s1b | 34.593 | false | inc | 8 |

| | id | BMD.zSD | TF | trend | friendliness |
|---|---|---|---|---|---|
| 10 | eif2s2 | 40.557 | false | inc | 1 |

Showing 1 to 10 of 28 entries

Previous 1 2 3 Next

Table 26: Table of the 16th cluster content

#### Cluster 28 - 28 transcripts (BMD q25 = 4.25 µg/L)

**Driver GO terms :**

**KEGG pathways :** Protein export, Protein processing in endoplasmic reticulum

**Wikipathways :**

Curvesplot BMDplot

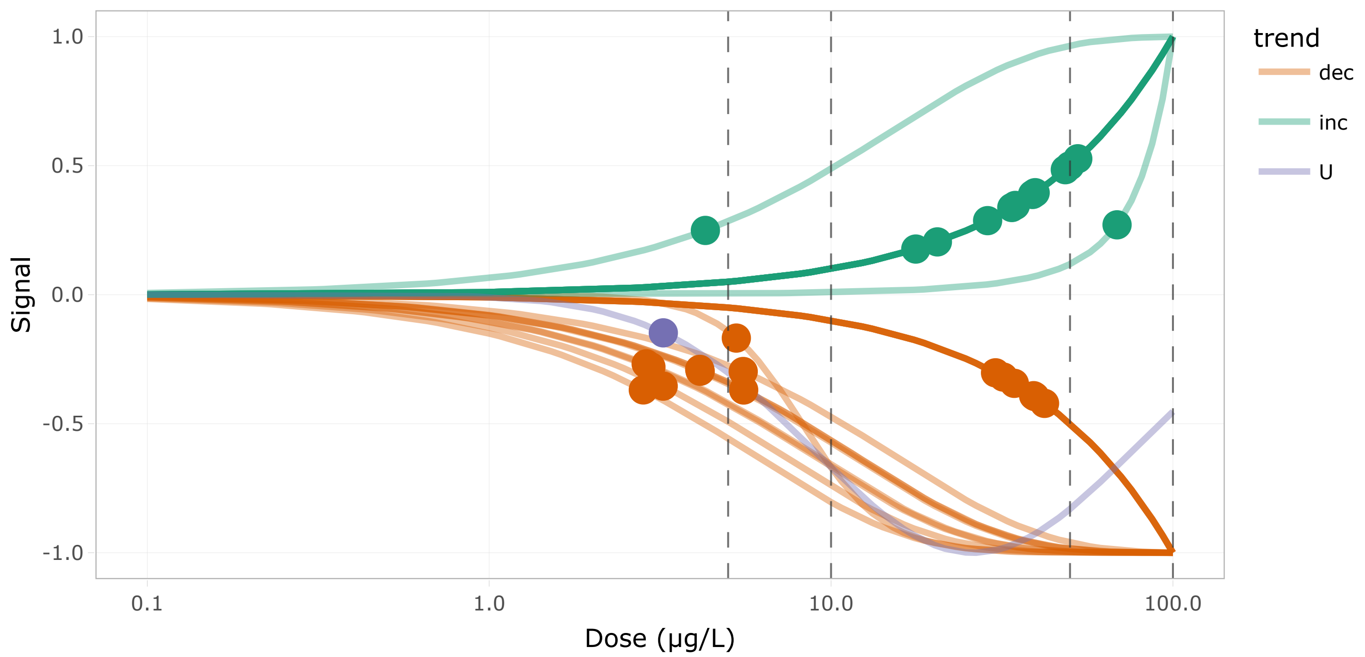


Figure 35: DR curves for the 17th cluster

Show 10 entries

Search:

| | id | BMD.zSD | TF | trend | friendliness |
|---|---|---|---|---|---|
| | All | All | All | All | All |
| 1 | RNF5 | 31.927 | false | dec | 4 |
| 2 | atf6 | 30.276 | true | dec | 4 |
| 3 | calr3b | 4.129 | false | dec | 6 |
| 4 | canx | 4.142 | false | dec | 5 |
| 5 | dnajb11 | 2.879 | false | dec | 5 |
| 6 | dnajc1 | 33.784 | true | inc | 4 |

| | id | BMD.zSD | TF | trend | friendliness |
|---|---|---|---|---|---|
| 7 | dnajc5aa | 39.493 | false | dec | 5 |
| 8 | eif2s1b | 34.593 | false | inc | 8 |
| 9 | ganab | 34.356 | false | dec | 4 |
| 10 | hsp70.1_t2 | 52.681 | false | inc | 7 |

Showing 1 to 10 of 28 entries

Previous 1 2 3 Next

Table 27: Table of the 17th cluster content

#### Cluster 50 - 24 transcripts (BMD q25 = 4.25 µg/L)

**Driver GO terms** : COPII-coated vesicle budding

**KEGG pathways** : Protein processing in endoplasmic reticulum

**Wikipathways** :

Curvesplot BMDplot

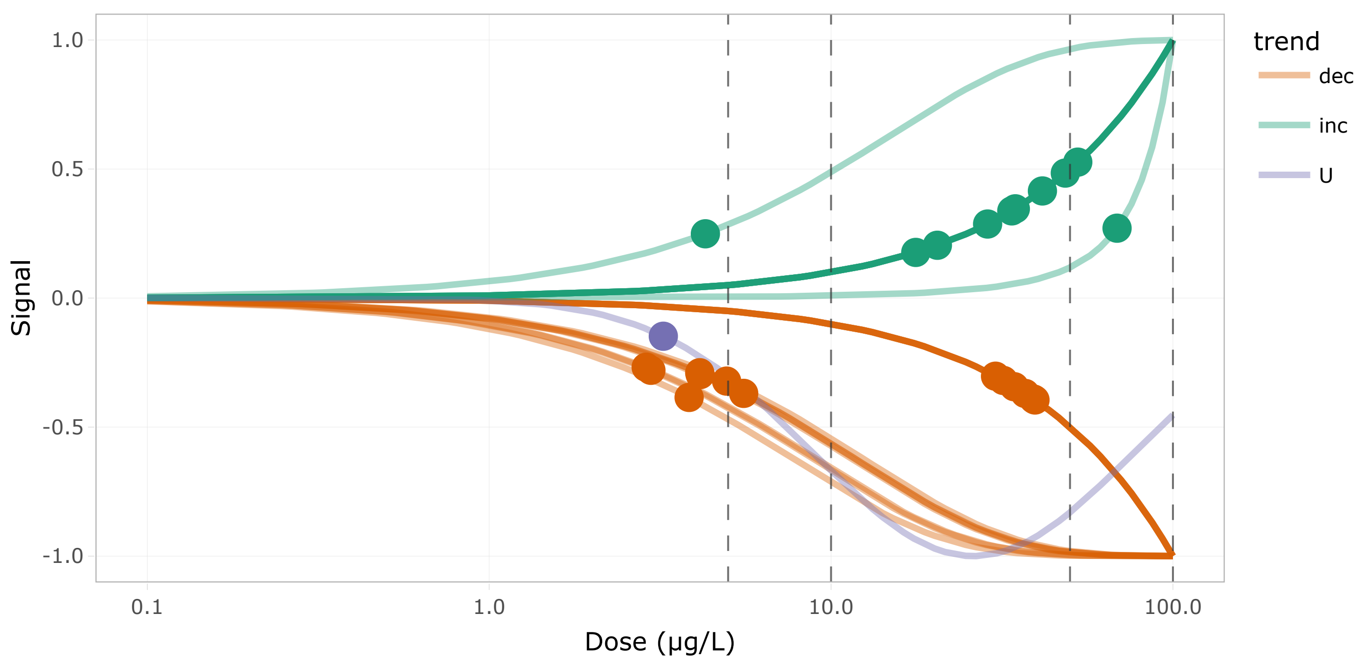


Figure 37: DR curves for the 18th cluster

Show 10 entries

Search:

| | id | BMD.zSD | TF | trend | friendliness |
|---|---|---|---|---|---|
| | All | All | All | All | All |
| 1 | RNF5 | 31.927 | false | dec | 4 |
| 2 | atf6 | 30.276 | true | dec | 4 |
| 3 | calr3b | 4.129 | false | dec | 6 |

| | id | BMD.zSD | TF | trend | friendliness |
|---|---|---|---|---|---|
| 4 | canx | 4.142 | false | dec | 5 |
| 5 | dnajb11 | 2.879 | false | dec | 5 |
| 6 | dnajc1 | 33.784 | true | inc | 4 |
| 7 | dnajc5aa | 39.493 | false | dec | 5 |
| 8 | eif2s1b | 34.593 | false | inc | 8 |
| 9 | ganab | 34.356 | false | dec | 4 |
| 10 | hsp70.1_t2 | 52.681 | false | inc | 7 |

Showing 1 to 10 of 24 entries

Previous 1 2 3 Next

Table 28: Table of the 18th cluster content

#### Cluster 31 - 14 transcripts (BMD q25 = 4.33 µg/L)

**Driver GO terms** :

**KEGG pathways** : ATP-dependent chromatin remodeling

**Wikipathways** :

Curvesplot BMDplot

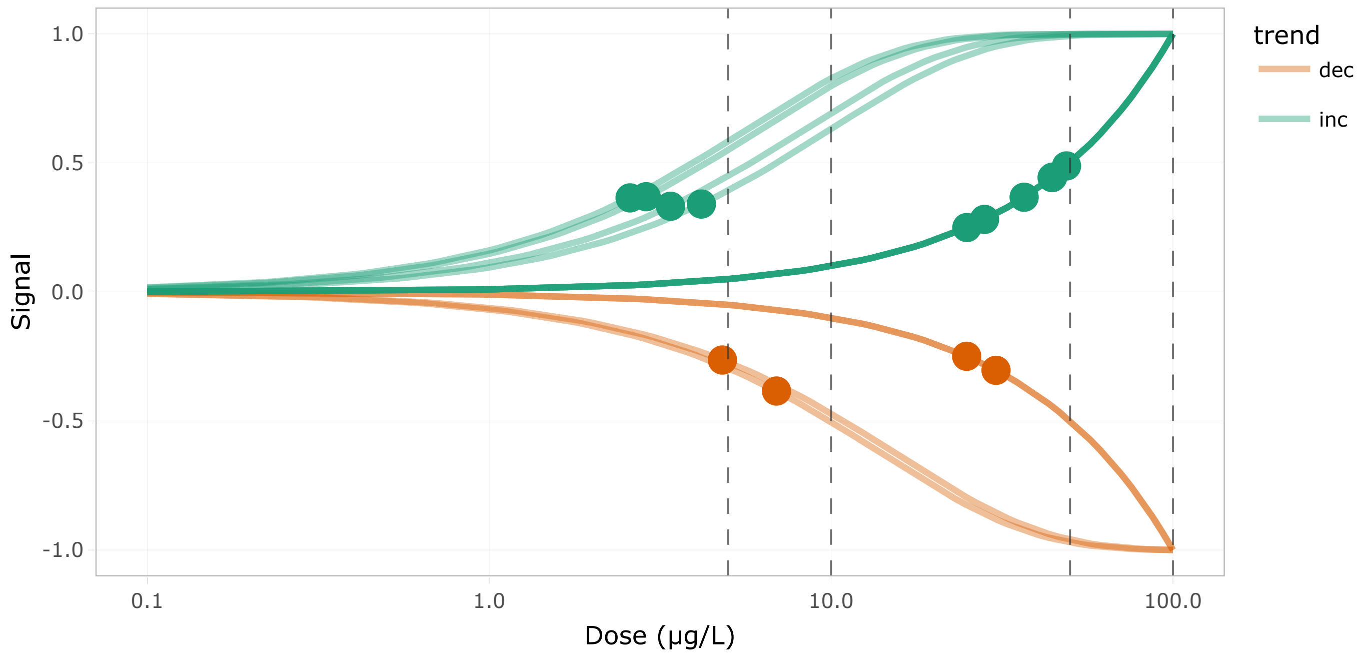


Figure 39: DR curves for the 19th cluster

Show 10 entries

Search:

| | id | BMD.zSD | TF | trend | friendliness |
|---|---|---|---|---|---|
| | All | All | All | All | All |
| 1 | actr8 | 44.489 | true | inc | 1 |
| 2 | brd8_t3 | 6.925 | false | dec | 1 |
| 3 | brd8_t4 | 4.175 | false | inc | 1 |
| 4 | chaf1a_g2t2 | 4.812 | false | dec | 1 |
| 5 | chd3_g2t2 | 30.379 | true | dec | 1 |
| 6 | chrac1_t2 | 24.966 | true | inc | 1 |
| 7 | ep400_t2 | 3.392 | true | inc | 1 |
| 8 | gatad2b | 24.931 | true | dec | 1 |
| 9 | h2ax | 28.11 | false | inc | 1 |
| 10 | lin52_t2 | 44.316 | false | inc | 1 |

Showing 1 to 10 of 14 entries

Previous 1 2 Next

Table 29: Table of the 19th cluster content

#### Cluster 4 - 54 transcripts (BMD q25 = 4.39 µg/L)

**Driver GO terms** : ATP synthesis coupled electron transport, generation of precursor metabolites and energy

**KEGG pathways** : Oxidative phosphorylation, Cardiac muscle contraction

**Wikipathways** : Electron transport chain, Oxidative phosphorylation

Curvesplot BMDplot

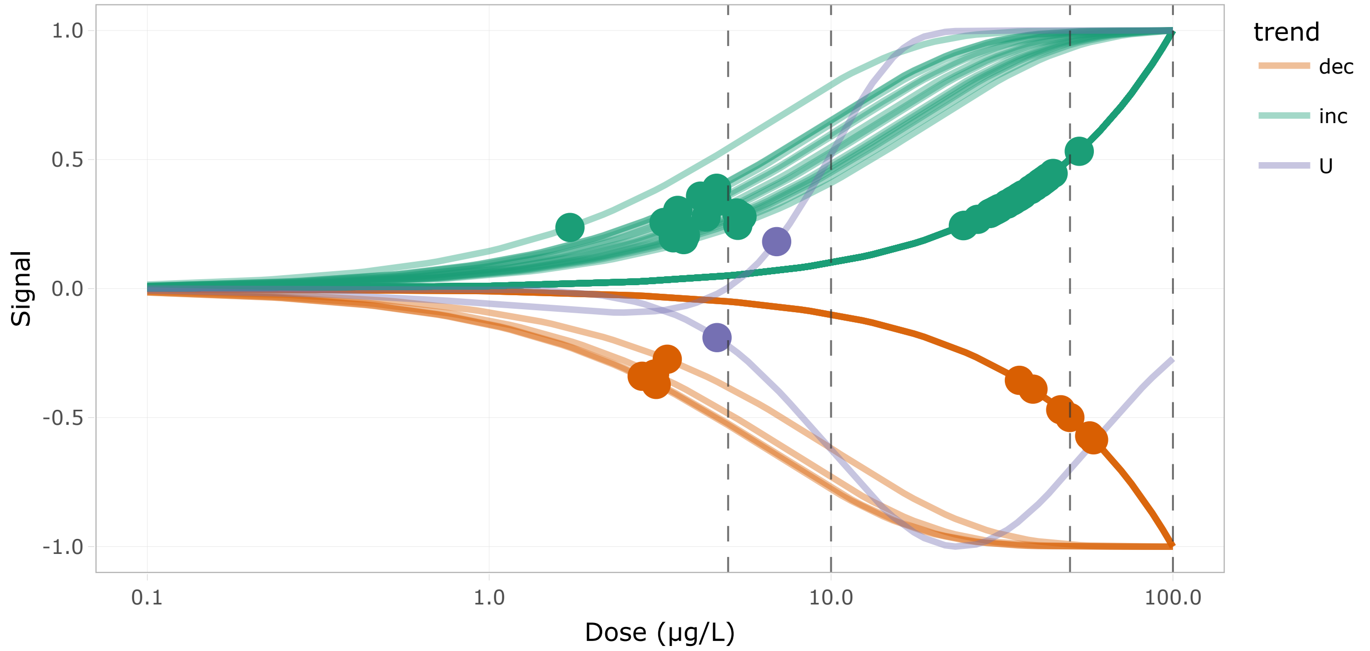


Figure 41: DR curves for the 20th cluster

Show 10 entries

Search:

| | id | BMD.zSD | TF | trend | friendliness |
|---|---|---|---|---|---|
| | All | All | All | All | All |
| 1 | CABZ01102240.1_t1 | 4.152 | false | inc | 1 |
| 2 | agla_t1 | 3.046 | false | dec | 2 |
| 3 | atp1a3a | 58.597 | false | dec | 1 |
| 4 | atp2a1_t1 | 42.978 | false | inc | 2 |
| 5 | atp2a2b | 3.318 | false | dec | 1 |
| 6 | atp2a3_t3 | 2.802 | false | dec | 1 |
| 7 | atp5f1c | 24.517 | false | inc | 1 |
| 8 | cacng1a | 30.612 | false | inc | 2 |
| 9 | coa6 | 4.642 | false | inc | 1 |
| 10 | cox11_t2 | 32.759 | false | inc | 1 |

Showing 1 to 10 of 54 entries

Previous 1 2 3 4 5 6 Next

Table 30: Table of the 20th cluster content

#### Cluster 14 - 30 transcripts (BMD q25 = 4.4 µg/L)

**Driver GO terms** : muscle system process

**KEGG pathways** : Vascular smooth muscle contraction

**Wikipathways** :

Curvesplot BMDplot

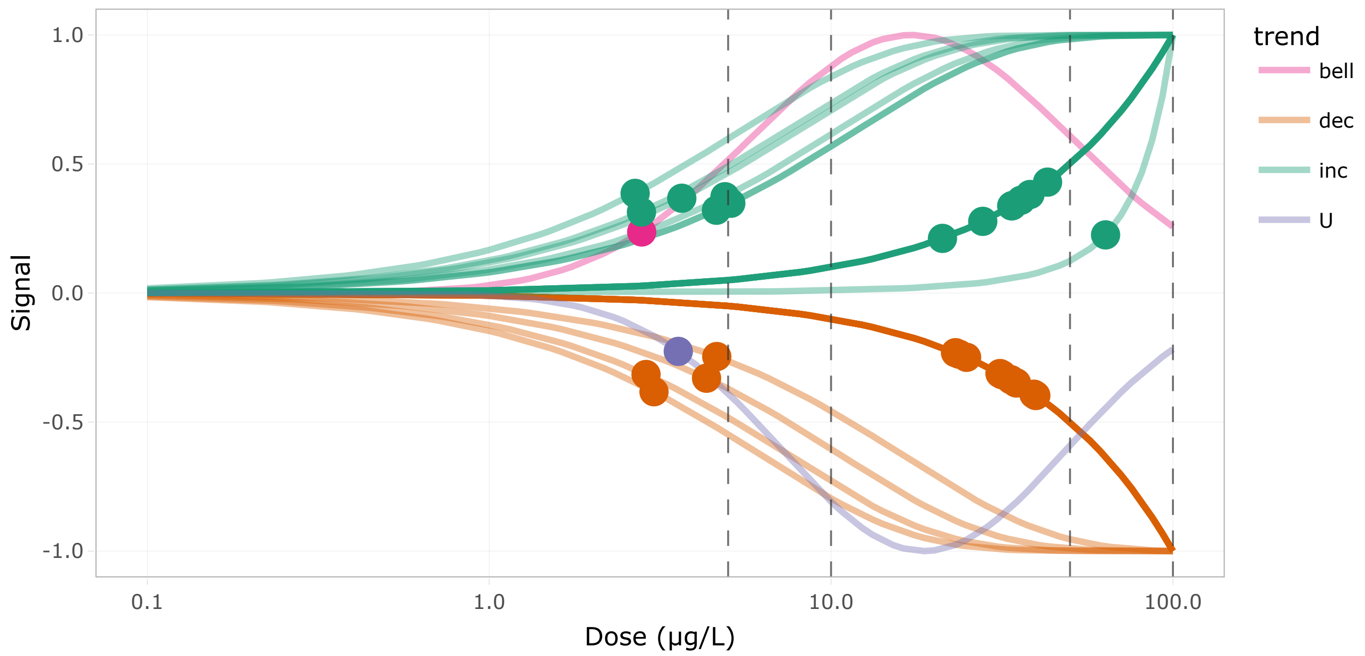


Figure 43: DR curves for the 21th cluster

Show 10 entries

Search:

| | id | BMD.zSD | TF | trend | friendliness |
|---|---|---|---|---|---|
| | All | All | All | All | All |
| 1 | BX936363.1 | 33.564 | false | dec | 1 |
| 2 | adma | 38.185 | false | inc | 1 |
| 3 | atp2a1_t1 | 42.978 | false | inc | 2 |
| 4 | braf_t2 | 2.794 | false | bell | 4 |
| 5 | cald1a | 4.632 | false | dec | 1 |
| 6 | cald1b_t2 | 23.697 | false | dec | 1 |
| 7 | cx39.9 | 27.779 | false | inc | 2 |
| 8 | erp44 | 3.034 | false | dec | 3 |
| 9 | foxo3b | 39.206 | true | dec | 1 |
| 10 | gna11b | 39.71 | false | dec | 1 |

Showing 1 to 10 of 30 entries

Previous 1 2 3 Next

Table 31: Table of the 21th cluster content

#### Cluster 11 - 52 transcripts (BMD q25 = 4.69 µg/L)

**Driver GO terms** :

**KEGG pathways** : Adherens junction, Mitophagy - animal, Ribosome biogenesis in eukaryotes, Wnt signaling pathway

**Wikipathways** :

Curvesplot BMDplot

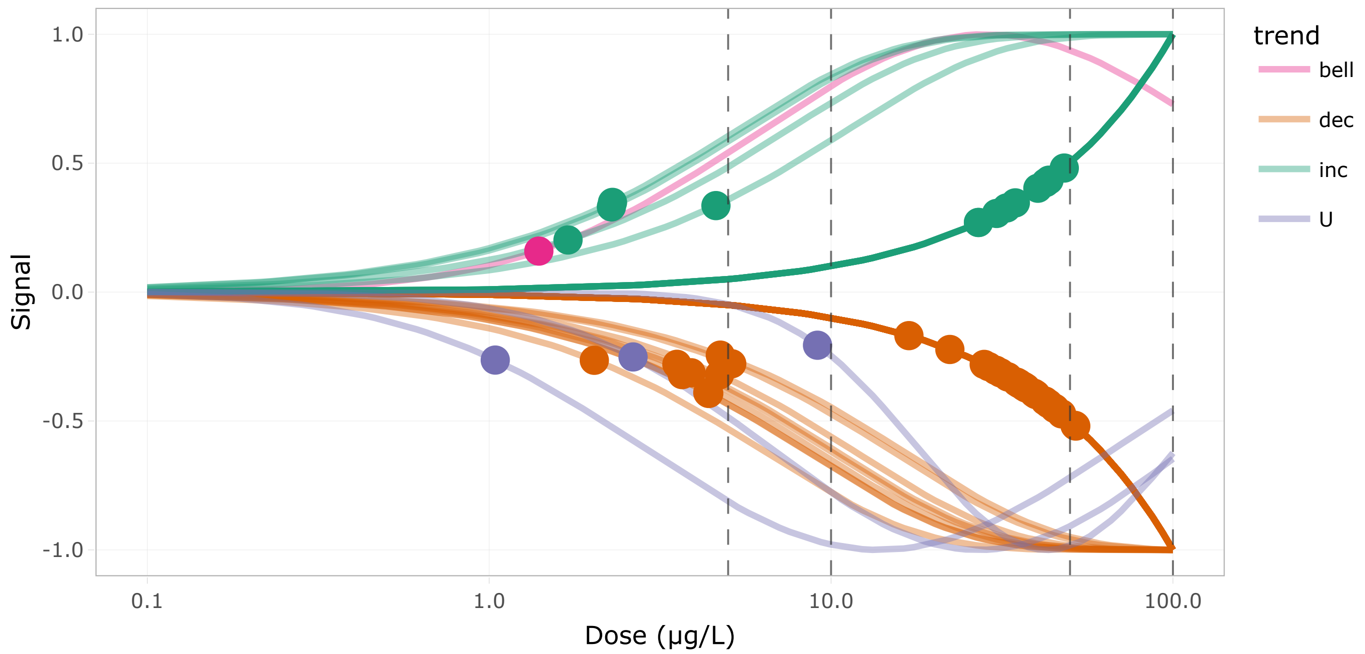


Figure 45: DR curves for the 22th cluster

Show 10 entries Search:

| | id | BMD.zSD | TF | trend | friendliness |
|---|---|---|---|---|---|
| | All | All | All | All | All |
| 1 | UTP14C | 51.973 | false | dec | 3 |
| 2 | acp1_t1 | 32.62 | false | inc | 1 |
| 3 | afdna_t1 | 45.241 | false | dec | 2 |
| 4 | atg9a_t4 | 43.399 | false | inc | 1 |
| 5 | bcl9_t2 | 44.947 | true | dec | 2 |
| 6 | cacybp | 26.991 | false | inc | 1 |
| 7 | cdh1_t2 | 4.36 | false | dec | 2 |
| 8 | clul1_g2t1 | 39.766 | false | dec | 1 |
| 9 | csnk1g2a_t2 | 30.313 | false | dec | 1 |
| 10 | csnk2a2a | 4.606 | false | inc | 1 |

Showing 1 to 10 of 52 entries

Previous 1 2 3 4 5 6 Next

Table 32: Table of the 22th cluster content

#### Cluster 22 - 15 transcripts (BMD q25 = 5.23 µg/L)

**Driver GO terms** : polyamine metabolic process

**KEGG pathways** : Arginine and proline metabolism

**Wikipathways** :

Curvesplot BMDplot

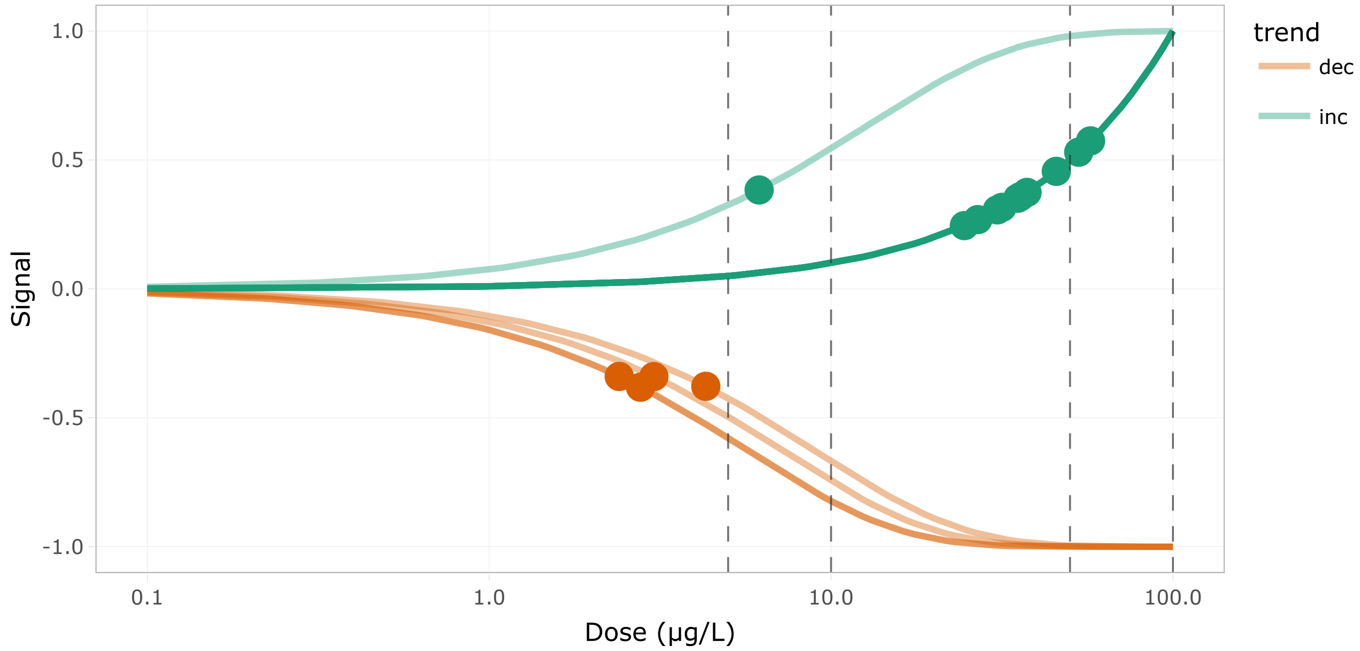


Figure 47: DR curves for the 23th cluster

Show 10 entries Search:

| | id | BMD.zSD | TF | trend | friendliness |
|---|---|---|---|---|---|
| | All | All | All | All | All |
| 1 | agmat | 2.771 | false | dec | 1 |
| 2 | aldh2.1 | 57.408 | false | inc | 1 |
| 3 | aldh3a2b | 3.032 | false | dec | 1 |
| 4 | arg2 | 24.542 | false | inc | 1 |
| 5 | azin1b | 53.019 | false | inc | 4 |
| 6 | ckmb | 30.64 | false | inc | 1 |
| 7 | mao | 2.401 | false | dec | 1 |
| 8 | oat_t2 | 35.164 | false | inc | 3 |
| 9 | oaz2a | 4.302 | false | dec | 1 |
| 10 | oaz2b | 45.589 | false | inc | 1 |

Showing 1 to 10 of 15 entries Previous 1 2 Next

Table 33: Table of the 23th cluster content

#### Cluster 5 - 45 transcripts (BMD q25 = 5.27 µg/L)

**Driver GO terms** : protein catabolic process

**KEGG pathways** : Proteasome

**Wikipathways :**

Curvesplot | BMDplot

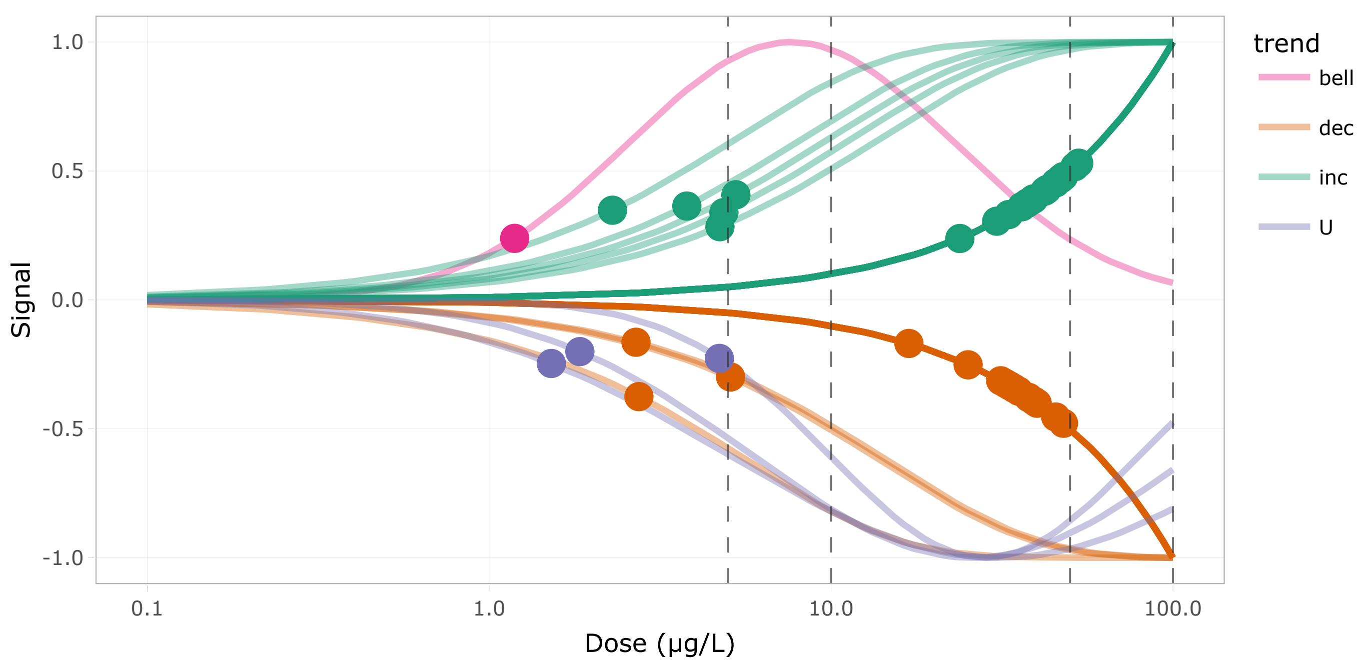


Figure 49: DR curves for the 24th cluster

Show 10 entries Search:

| | id | BMD.zSD | TF | trend | friendliness |
|---|---|---|---|---|---|
| | All | All | All | All | All |
| 1 | CR788324.2 | 33.242 | false | dec | 1 |
| 2 | anapc4_t2 | 4.732 | false | inc | 3 |
| 3 | appbp2 | 40.076 | false | dec | 1 |
| 4 | arih2_t1 | 2.689 | false | dec | 1 |
| 5 | azin1b | 53.019 | false | inc | 4 |
| 6 | cul2 | 30.52 | false | inc | 2 |
| 7 | cylda_t2 | 2.295 | false | inc | 3 |
| 8 | desi1a | 5.268 | false | inc | 1 |
| 9 | fbxo33 | 37.759 | false | dec | 1 |
| 10 | hectd3_t2 | 2.742 | false | dec | 1 |

Showing 1 to 10 of 45 entries

Previous 1 2 3 4 5 Next

Table 34: Table of the 24th cluster content

### Cluster 33 - 32 transcripts (BMD q25 = 6.28 µg/L)

**Driver GO terms** : microtubule-based process

**KEGG pathways** :

**Wikipathways :**

Curvesplot | BMDplot

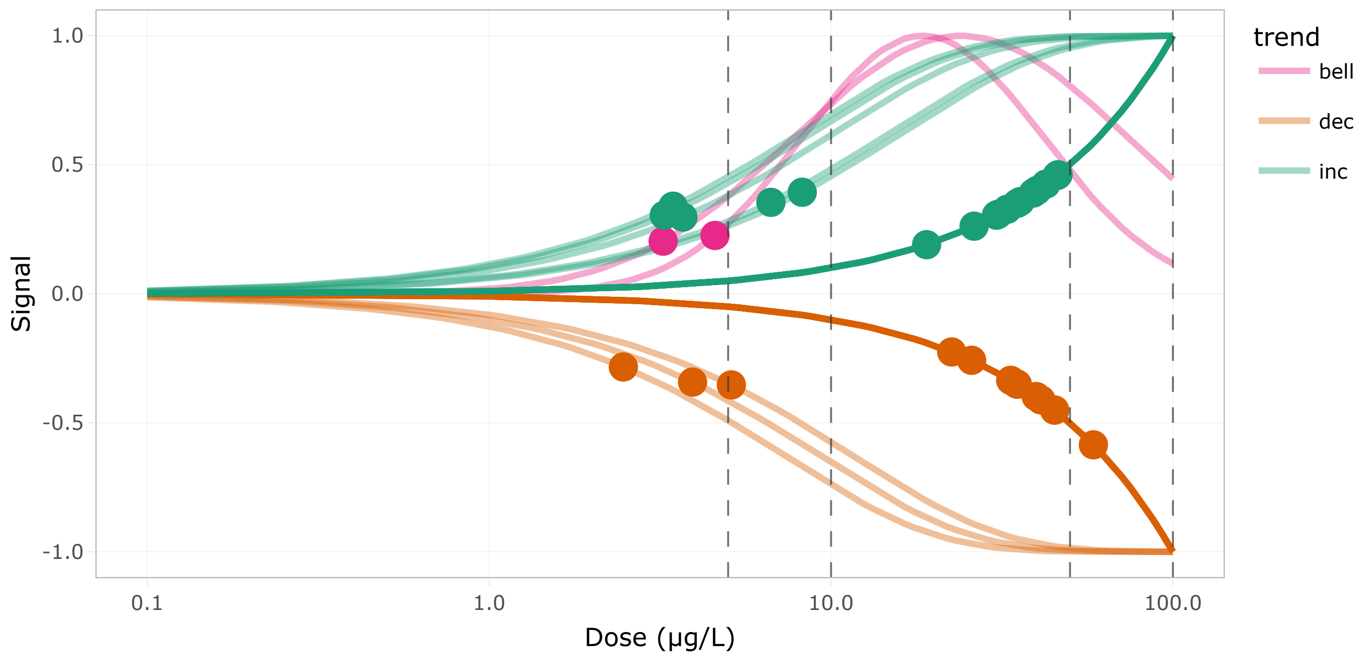


Figure 51: DR curves for the 25th cluster

Show 10 entries

Search:

| | id | BMD.zSD | TF | trend | friendliness |
|---|---|---|---|---|---|
| | All | All | All | All | All |
| 1 | CU929259.1 | 22.537 | false | dec | 1 |
| 2 | cep192_t2 | 40.049 | false | inc | 1 |
| 3 | cep57 | 3.253 | false | inc | 1 |
| 4 | cfap298 | 19.032 | false | inc | 1 |
| 5 | ckap5_t2 | 8.236 | false | inc | 1 |
| 6 | cpeb2_t1 | 3.228 | false | bell | 1 |
| 7 | cpeb3 | 34.885 | false | dec | 1 |
| 8 | dynll2b | 3.689 | false | inc | 3 |
| 9 | fyco1a_t2 | 3.935 | false | dec | 2 |
| 10 | gas2l1_t2 | 25.772 | false | dec | 1 |

Showing 1 to 10 of 32 entries

Previous 1 2 3 4 Next

Table 35: Table of the 25th cluster content

### Cluster 17 - 22 transcripts (BMD q25 = 6.37 µg/L)

**Driver GO terms :**

**KEGG pathways :** Ubiquitin mediated proteolysis

**Wikipathways :**

Curvesplot | BMDplot

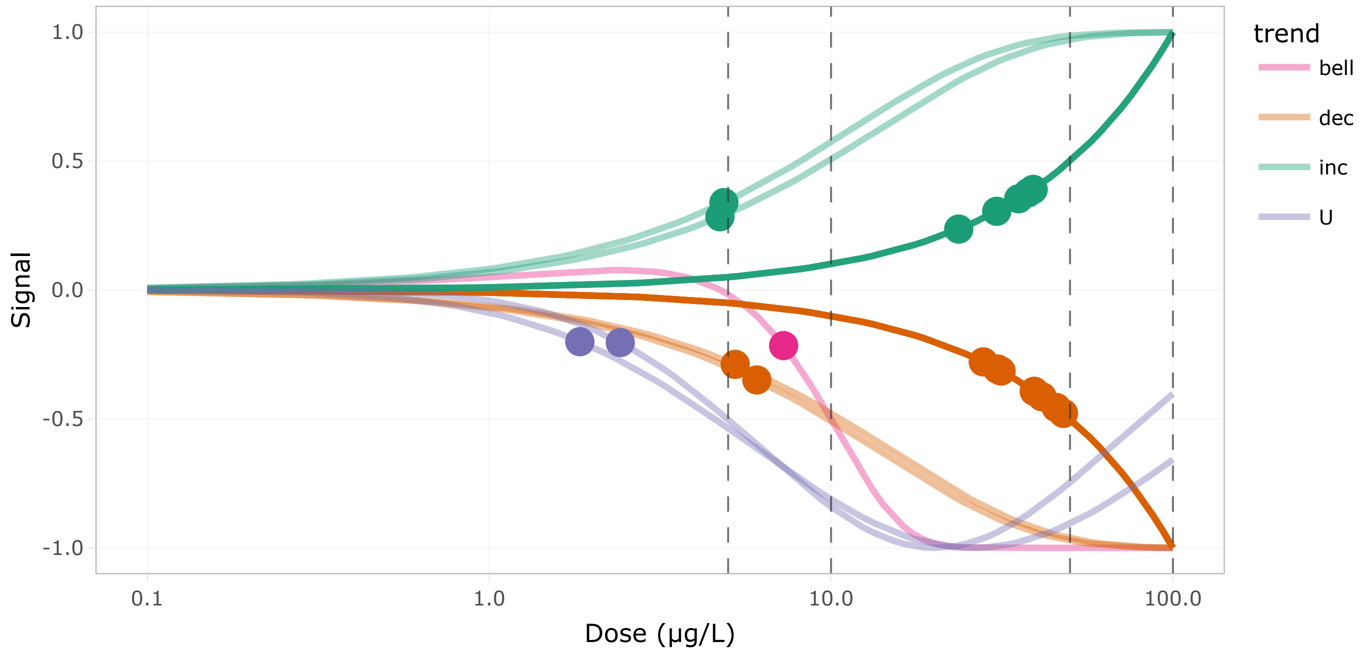


Figure 53: DR curves for the 26th cluster

Show 10 entries

Search:

| | id | BMD.zSD | TF | trend | friendliness |
|---|---|---|---|---|---|
| | All | All | All | All | All |
| 1 | UBE2M_t4 | 35.376 | false | inc | 1 |
| 2 | UBE2M_t5 | 30.787 | false | dec | 1 |
| 3 | anapc4_t2 | 4.732 | false | inc | 3 |
| 4 | anapc5 | 23.627 | false | inc | 2 |
| 5 | cbl | 41.388 | true | dec | 1 |
| 6 | cdc34b | 31.425 | false | dec | 1 |
| 7 | cul2 | 30.52 | false | inc | 2 |
| 8 | itcha | 47.842 | false | dec | 2 |
| 9 | itchb | 39.285 | true | dec | 2 |
| 10 | map3k1 | 5.243 | false | dec | 3 |

Showing 1 to 10 of 22 entries

Previous 1 2 3 Next

Table 36: Table of the 26th cluster content

### Cluster 1 - 161 transcripts (BMD q25 = 6.79 µg/L)

**Driver GO terms** : embryo development ending in birth or egg hatching, myeloid cell differentiation, ribonucleoprotein complex biogenesis, translation, ribosomal small subunit assembly

**KEGG pathways** : Ribosome

**Wikipathways** : Cytoplasmic ribosomal proteins

Curvesplot BMDplot

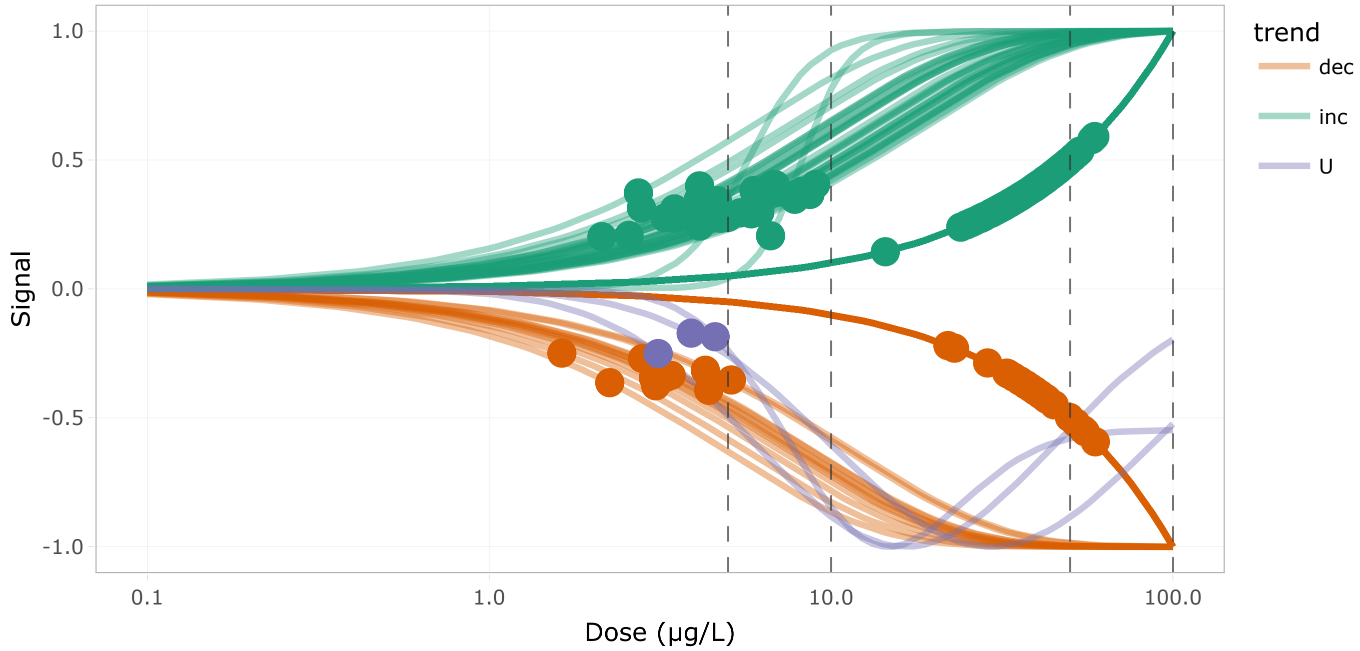


Figure 55: DR curves for the 27th cluster

Show 10 entries Search:

| | id | BMD.zSD | TF | trend | friendliness |
|---|---|---|---|---|---|
| | All | All | All | All | All |
| 1 | UTP14C | 51.973 | false | dec | 3 |
| 2 | aars1_g2t2 | 14.411 | false | inc | 2 |
| 3 | ahsa1a_t3 | 4.506 | false | inc | 2 |
| 4 | bcl9_t2 | 44.947 | true | dec | 2 |
| 5 | btf3 | 27.546 | false | inc | 1 |
| 6 | btf3l4 | 25.572 | false | inc | 1 |
| 7 | cdc73_t2 | 3.897 | true | U | 2 |
| 8 | chd7_t3 | 54.656 | true | dec | 1 |
| 9 | cnbpa_t1 | 47.732 | false | inc | 1 |
| 10 | cnot1_g2t1 | 39.652 | false | dec | 1 |

Showing 1 to 10 of 161 entries Previous 1 2 3 4 5 … 17 Next

Table 37: Table of the 27th cluster content

### Cluster 25 - 6 transcripts (BMD q25 = 7.85 µg/L)

**Driver GO terms :**

**KEGG pathways :**

**Wikipathways :**

Curvesplot BMDplot

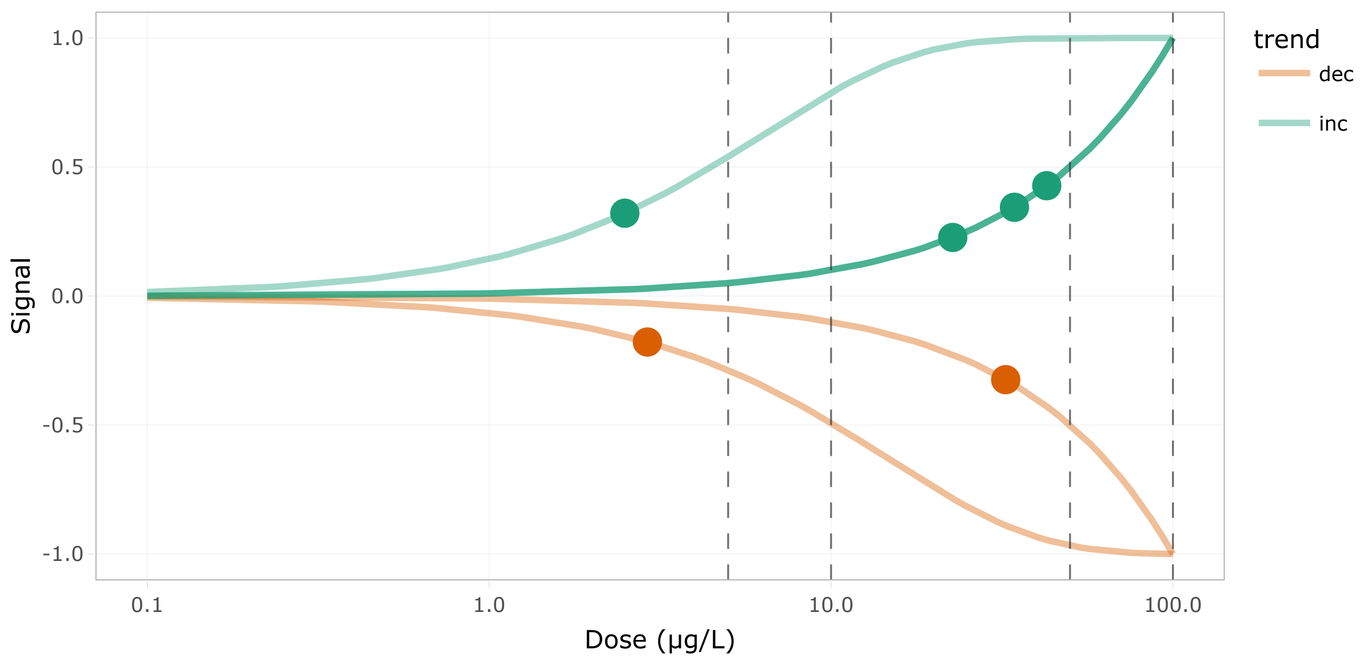


Figure 57: DR curves for the 28th cluster

Show 10 entries Search:

| | id | BMD.zSD | TF | trend | friendliness |
|---|---|---|---|---|---|
| | All | All | All | All | All |
| 1 | bms1_g2t1 | 2.903 | false | dec | 1 |
| 2 | glmna | 34.388 | false | inc | 1 |
| 3 | ncoa6_t2 | 2.494 | false | inc | 1 |
| 4 | ncoa6_t3 | 32.422 | false | dec | 1 |
| 5 | ngdn | 42.746 | false | inc | 1 |
| 6 | pno1 | 22.695 | false | inc | 1 |

Showing 1 to 6 of 6 entries

Previous 1 Next

Table 38: Table of the 28th cluster content

### Cluster 3 - 23 transcripts (BMD q25 = 13.65 µg/L)

**Driver GO terms :** tRNA aminoacylation

**KEGG pathways :** Aminoacyl-tRNA biosynthesis

**Wikipathways :**

Curvesplot BMDplot

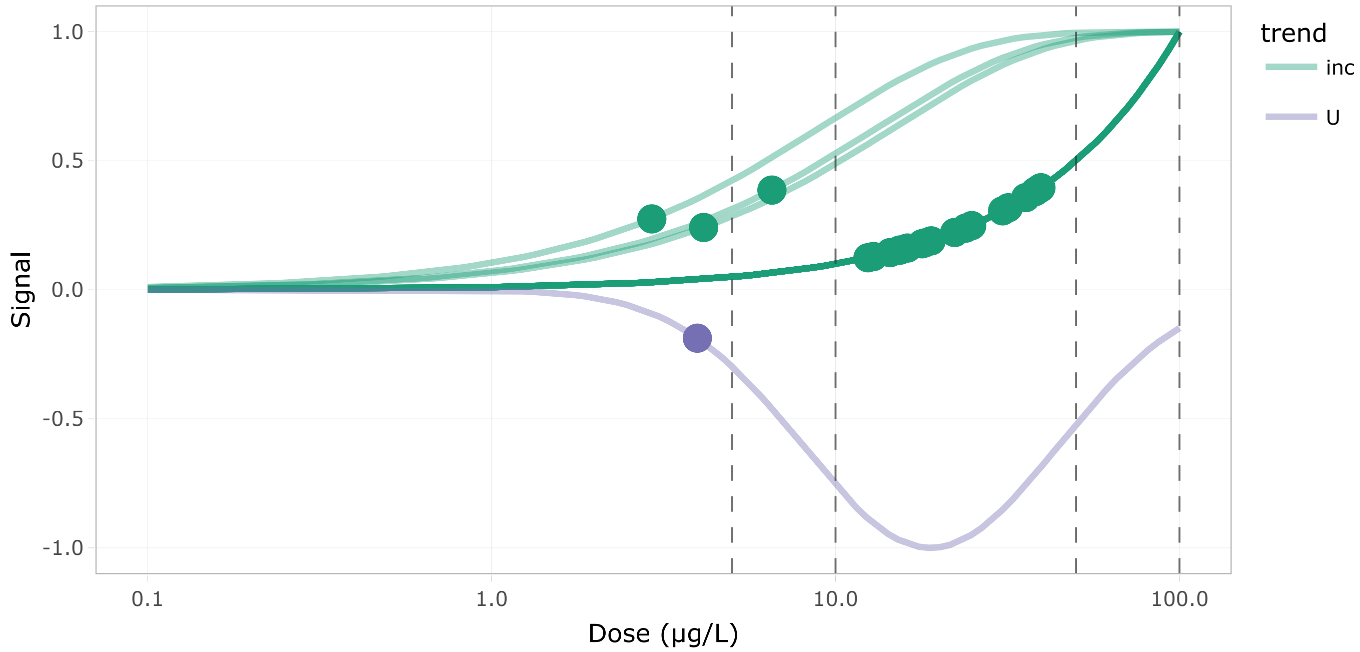


Figure 59: DR curves for the 29th cluster

Show 10 entries Search:

| | id | BMD.zSD | TF | trend | friendliness |
|---|---|---|---|---|---|
| | All | All | All | All | All |
| 1 | LO017852.1 | 6.532 | false | inc | 1 |
| 2 | WARS1 | 39.552 | false | inc | 1 |
| 3 | aars1_g1t2 | 15.339 | false | inc | 1 |
| 4 | aars1_g2t2 | 14.411 | false | inc | 2 |
| 5 | aimp1a | 35.732 | false | inc | 1 |
| 6 | aimp1b_t4 | 4.136 | false | inc | 1 |
| 7 | aimp2_t2 | 38.006 | false | inc | 1 |
| 8 | dars1 | 17.833 | false | inc | 1 |
| 9 | eprs1_t2 | 15.442 | false | inc | 1 |
| 10 | farsa | 16.181 | false | inc | 1 |

Showing 1 to 10 of 23 entries Previous 1 2 3 Next

Table 39: Table of the 29th cluster content

#### Cluster 39 - 7 transcripts (BMD q25 = 17.7 µg/L)

**Driver GO terms** : mitotic chromosome condensation

**KEGG pathways** :

**Wikipathways** :

Curvesplot BMDplot

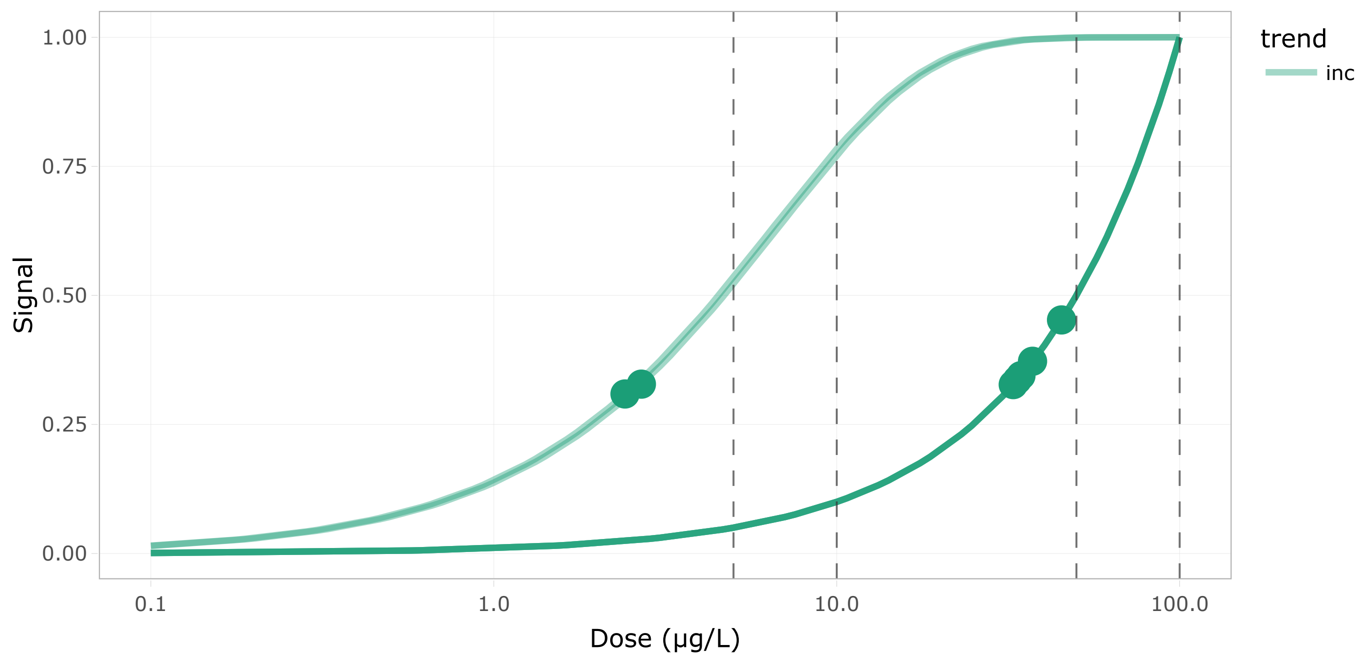


Figure 61: DR curves for the 30th cluster

Show 10 entries Search:

| | id | BMD.zSD | TF | trend | friendliness |
|---|---|---|---|---|---|
| | All | All | All | All | All |
| 1 | ncapd3_g1t1 | 2.693 | false | inc | 1 |
| 2 | ncapd3_g2t1 | 45.242 | false | inc | 1 |
| 3 | ncapg2_t2 | 37.233 | false | inc | 1 |
| 4 | ncapg_g1t2 | 2.413 | false | inc | 1 |
| 5 | ncapg_g3t1 | 32.715 | false | inc | 1 |
| 6 | ncaph_g2t1 | 33.606 | false | inc | 1 |
| 7 | smc2 | 34.447 | false | inc | 1 |

Showing 1 to 7 of 7 entries Previous 1 Next

Table 40: Table of the 30th cluster content

#### Cluster 46 - 21 transcripts (BMD q25 = 17.89 μg/L)

**Driver GO terms** : DNA alkylation, methylation

**KEGG pathways** : Cysteine and methionine metabolism

**Wikipathways :**

Curvesplot | BMDplot

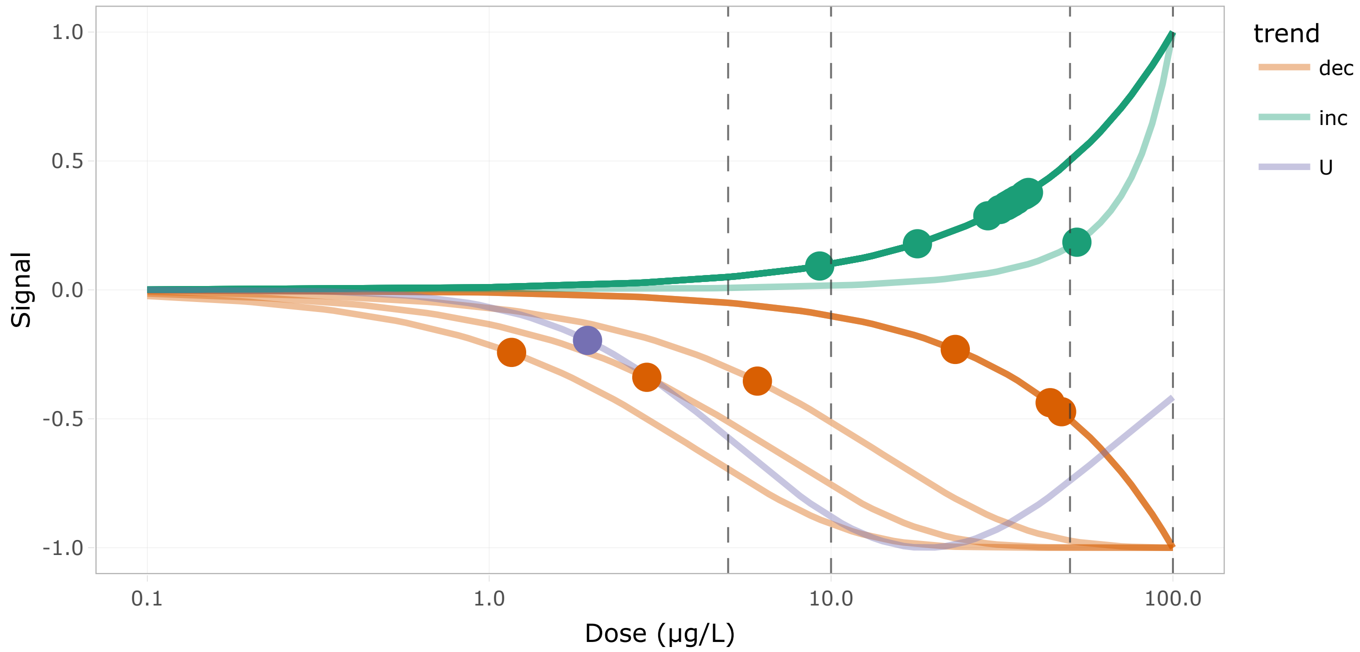


Figure 63: DR curves for the 31th cluster

Show 10 entries

Search:

| | id | BMD.zSD | TF | trend | friendliness |
|---|---|---|---|---|---|
| | All | All | All | All | All |
| 1 | adi1 | 28.749 | false | inc | 3 |
| 2 | apip | 37.287 | false | inc | 2 |
| 3 | dnmt1 | 34.061 | true | inc | 1 |
| 4 | dnmt3aa_t2 | 43.738 | false | dec | 1 |
| 5 | dnmt3ab_t2 | 23.084 | false | dec | 1 |
| 6 | dnmt3bb.1_g2t1 | 6.089 | false | dec | 2 |
| 7 | dnmt3bb.3_g1t2 | 36.719 | false | inc | 2 |
| 8 | dot1l_t3 | 47.243 | false | dec | 3 |
| 9 | gclm | 2.892 | false | dec | 2 |
| 10 | mto1 | 52.371 | false | inc | 1 |

Showing 1 to 10 of 21 entries

Previous 1 2 3 Next

Table 41: Table of the 31th cluster content

#### Cluster 20 - 31 transcripts (BMD q25 = 25.13 µg/L)

**Driver GO terms** : DNA geometric change, DNA replication

**KEGG pathways** : Cell cycle, DNA replication

**Wikipathways** : Cell cycle, DNA replication, G1 to S cell cycle control

Curvesplot BMDplot

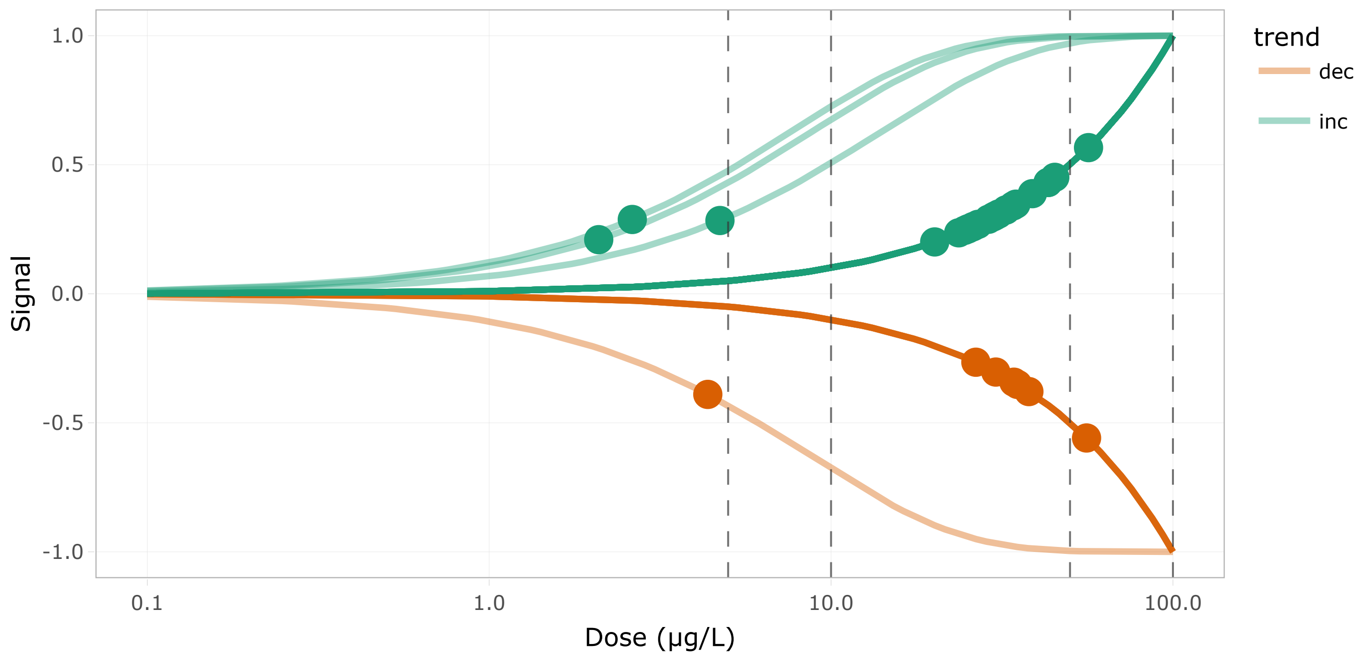


Figure 65: DR curves for the 32th cluster

Show 10 entries Search:

| | id | BMD.zSD | TF | trend | friendliness |
|---|---|---|---|---|---|
| | All | All | All | All | All |
| 1 | anapc4_t2 | 4.732 | false | inc | 3 |
| 2 | anapc5 | 23.627 | false | inc | 2 |
| 3 | atrx_g2t1 | 55.886 | true | dec | 2 |
| 4 | ccnb1_t2 | 24.594 | false | inc | 2 |
| 5 | ccne2_t3 | 32.539 | false | inc | 2 |
| 6 | cdh1_t2 | 4.36 | false | dec | 2 |
| 7 | cdk2 | 25.841 | true | inc | 2 |
| 8 | cdkn1ca | 45.124 | false | inc | 2 |
| 9 | cenps_g2t1 | 34.058 | false | inc | 2 |
| 10 | dna2 | 34.726 | false | inc | 2 |

Showing 1 to 10 of 31 entries Previous 1 2 3 4 Next

Table 42: Table of the 32th cluster content

#### Cluster 16 - 54 transcripts (BMD q25 = 25.18 µg/L)

**Driver GO terms** : DNA metabolic process, DNA recombination

**KEGG pathways** : DNA replication, Base excision repair, Mismatch repair, Nucleotide excision repair, Fanconi anemia pathway, Homologous recombination

**Wikipathways** : DNA replication, G1 to S cell cycle control

Curvesplot BMDplot

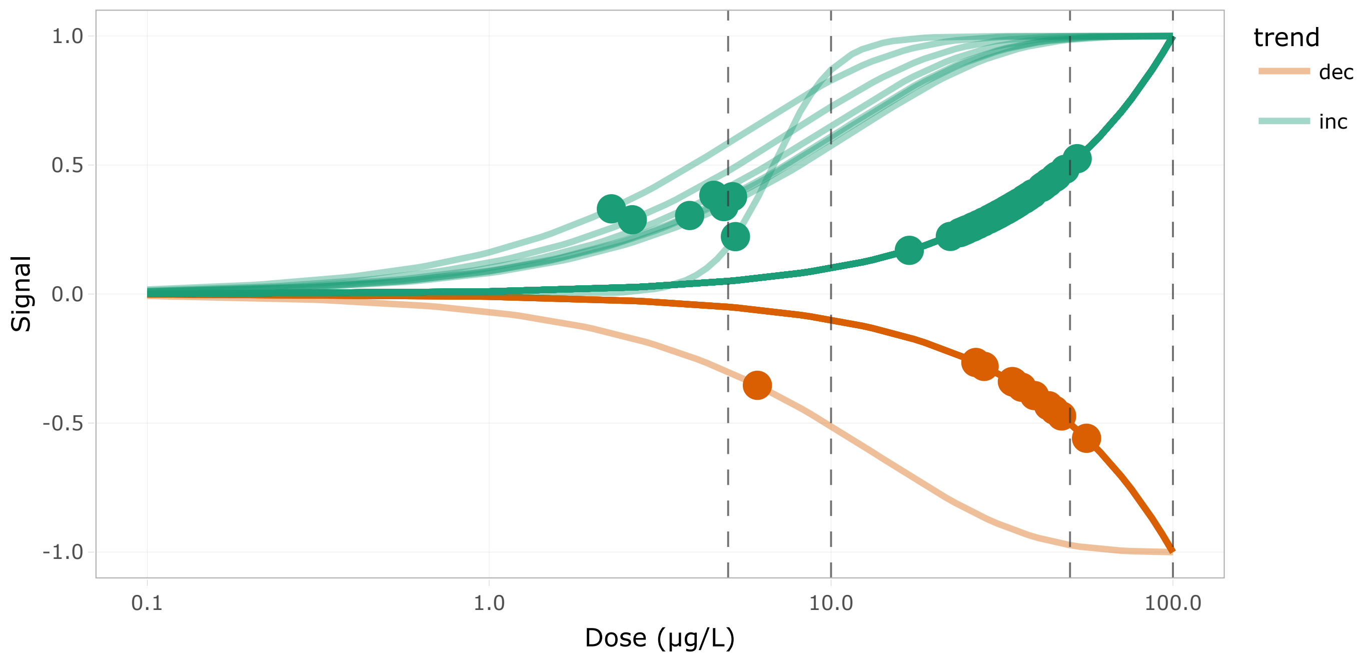


Figure 67: DR curves for the 33th cluster

Show 10 entries Search:

| | id | BMD.zSD | TF | trend | friendliness |
|---|---|---|---|---|---|
| | All | All | All | All | All |
| 1 | FO834799.1 | 5.151 | false | inc | 1 |
| 2 | atrx_g2t1 | 55.886 | true | dec | 2 |
| 3 | brca2 | 16.94 | true | inc | 1 |
| 4 | ccdc36 | 4.542 | false | inc | 1 |
| 5 | ccnb1_t2 | 24.594 | false | inc | 2 |
| 6 | ccne2_t3 | 32.539 | false | inc | 2 |
| 7 | cdca7a | 31.664 | false | inc | 1 |
| 8 | cdk2 | 25.841 | true | inc | 2 |
| 9 | cdkn1ca | 45.124 | false | inc | 2 |
| 10 | cenps_g2t1 | 34.058 | false | inc | 2 |

Showing 1 to 10 of 54 entries Previous 1 2 3 4 5 6 Next

Table 43: Table of the 33th cluster content

### Cluster 34 - 15 transcripts (BMD q25 = 25.53 µg/L)

**Driver GO terms :**

**KEGG pathways :** Nucleotide metabolism, Pyrimidine metabolism

**Wikipathways :**

Curvesplot BMDplot

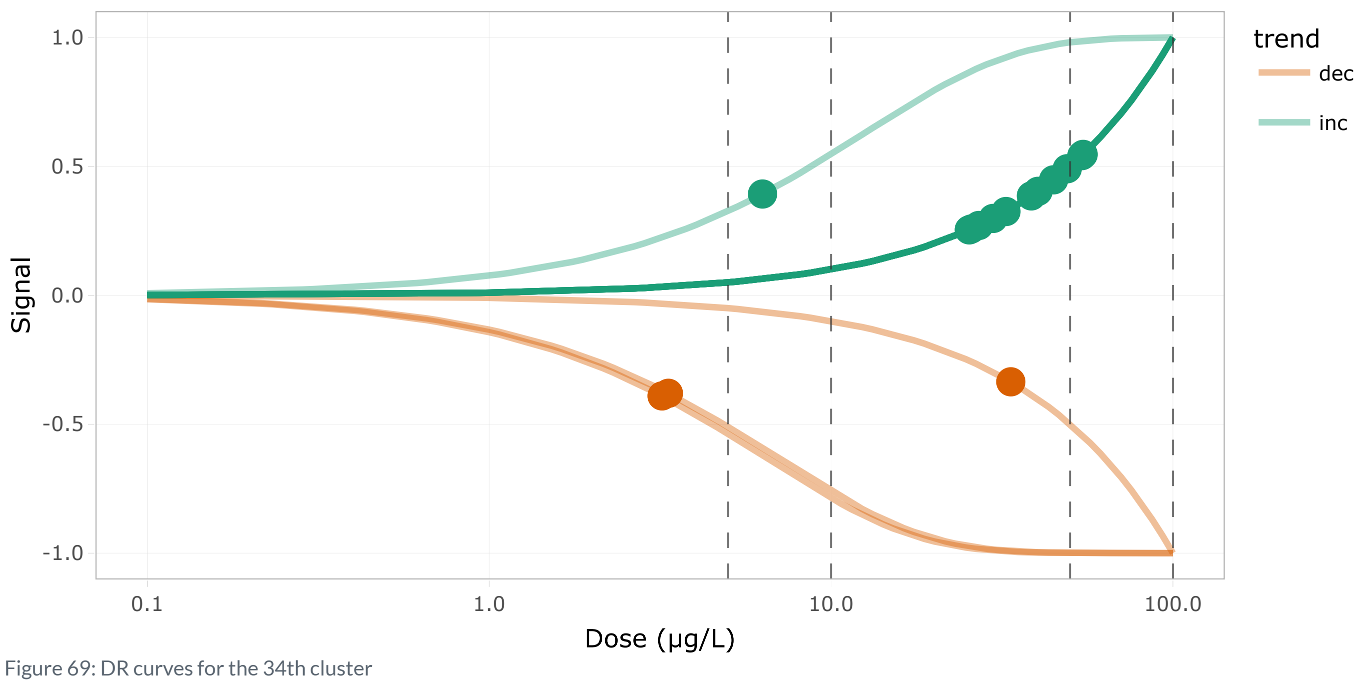


Figure 69: DR curves for the 34th cluster

Show 10 entries Search:

| | id | BMD.zSD | TF | trend | friendliness |
|---|---|---|---|---|---|
| | All | All | All | All | All |
| 1 | adka_t2 | 3.344 | false | dec | 1 |
| 2 | adkb_t2 | 54.629 | false | inc | 1 |
| 3 | ak1 | 6.303 | false | inc | 1 |
| 4 | cad_t2 | 27.057 | false | inc | 3 |
| 5 | dut | 32.481 | false | inc | 1 |
| 6 | gldc | 25.707 | false | inc | 1 |
| 7 | impdh1b_t2 | 33.561 | false | dec | 1 |
| 8 | mthfd1a_t2 | 3.204 | false | dec | 1 |
| 9 | nme6 | 38.532 | false | inc | 1 |
| 10 | nt5c2a | 49.076 | false | inc | 1 |

Showing 1 to 10 of 15 entries

Previous 1 2 Next

Table 44: Table of the 34th cluster content

### Cluster 19 - 26 transcripts (BMD q25 = 25.56 µg/L)

**Driver GO terms** : transmembrane receptor protein serine/threonine kinase signaling pathway

**KEGG pathways** : Cytokine-cytokine receptor interaction, TGF-beta signaling pathway

**Wikipathways** :

Curvesplot | BMDplot

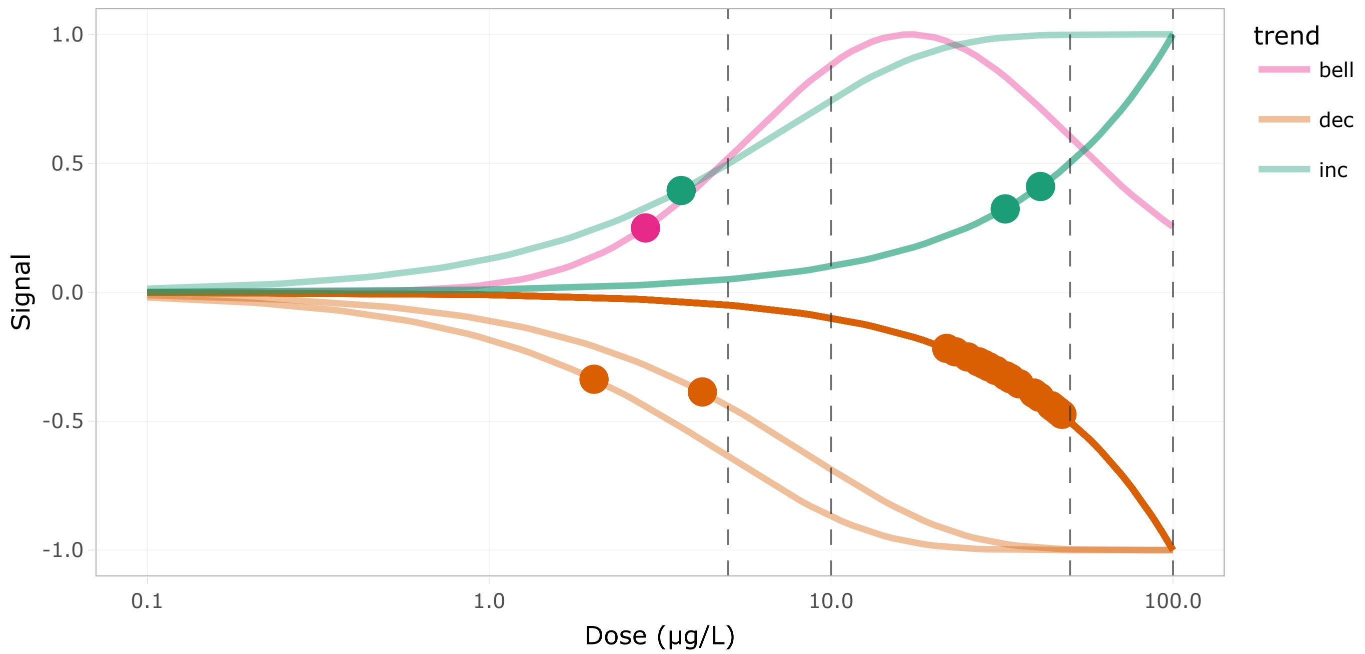


Figure 71: DR curves for the 35th cluster

Show 10 entries

Search:

| | id | BMD.zSD | TF | trend | friendliness |
|---|---|---|---|---|---|
| | All | All | All | All | All |
| 1 | acvr1ba | 39.099 | false | dec | 1 |
| 2 | acvr1bb_t2 | 26.966 | false | dec | 1 |
| 3 | acvr2ba_t3 | 25.085 | false | dec | 1 |
| 4 | acvr2bb | 39.523 | false | dec | 1 |
| 5 | bmpr2a | 46.195 | false | dec | 1 |
| 6 | bmpr2b | 39.152 | false | dec | 1 |
| 7 | flcn_t2 | 3.644 | false | inc | 1 |
| 8 | hdr_t2 | 2.026 | false | dec | 3 |
| 9 | hjv_t2 | 32.319 | false | inc | 1 |
| 10 | il10rb_t2 | 40.99 | false | inc | 1 |

Showing 1 to 10 of 26 entries

Previous 1 2 3 Next

Table 45: Table of the 35th cluster content

#### Cluster 18 - 6 transcripts (BMD q25 = 25.82 µg/L)

**Driver GO terms** :

**KEGG pathways** :

**Wikipathways** :

Curvesplot BMDplot

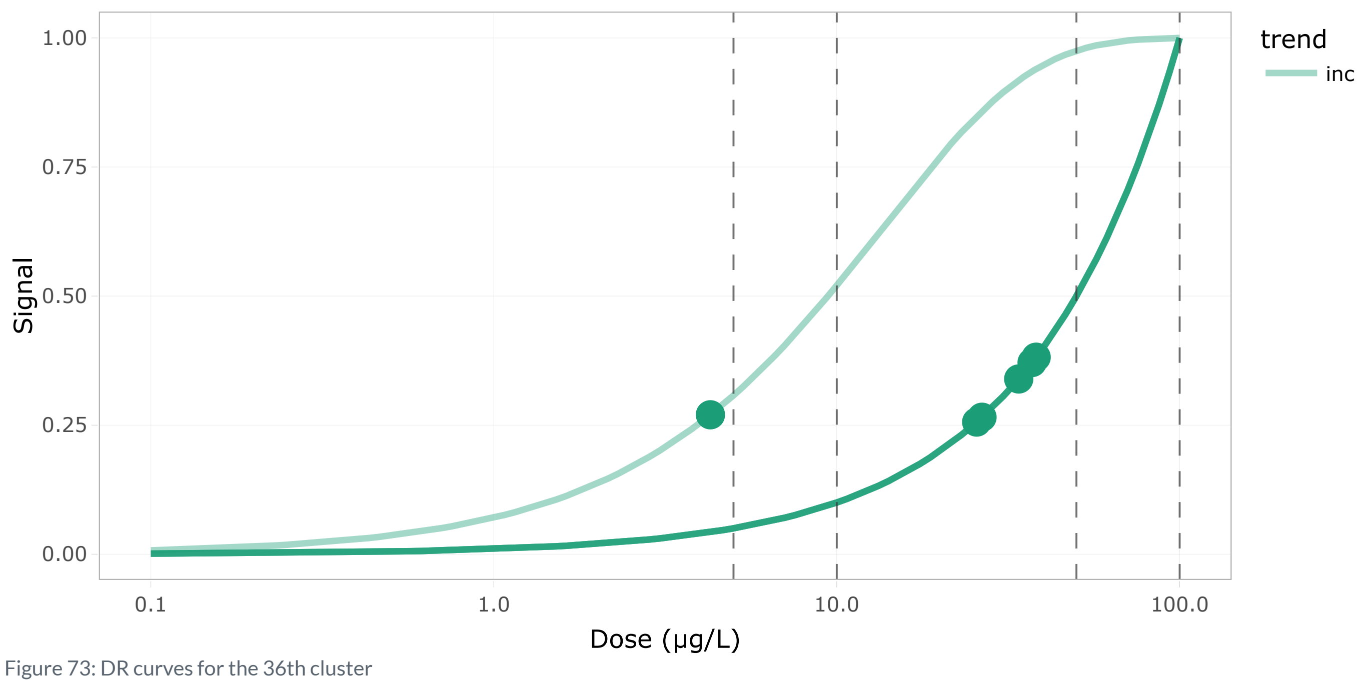


Figure 73: DR curves for the 36th cluster

Show 10 entries Search:

| | id | BMD.zSD | TF | trend | friendliness |
|---|---|---|---|---|---|
| | All | All | All | All | All |
| 1 | HTRA2_g1t2 | 4.282 | false | inc | 1 |
| 2 | birc5a | 25.592 | true | inc | 1 |
| 3 | cdca8 | 33.936 | false | inc | 1 |
| 4 | melk | 26.522 | false | inc | 1 |
| 5 | nusap1 | 37.128 | false | inc | 1 |
| 6 | ube2c_g2t1 | 38.154 | false | inc | 1 |

Showing 1 to 6 of 6 entries Previous 1 Next

Table 46: Table of the 36th cluster content

#### Cluster 36 - 10 transcripts (BMD q25 = 26.62 µg/L)

**Driver GO terms** : regulatory ncRNA-mediated gene silencing

**KEGG pathways :**

**Wikipathways :**

Curvesplot BMDplot

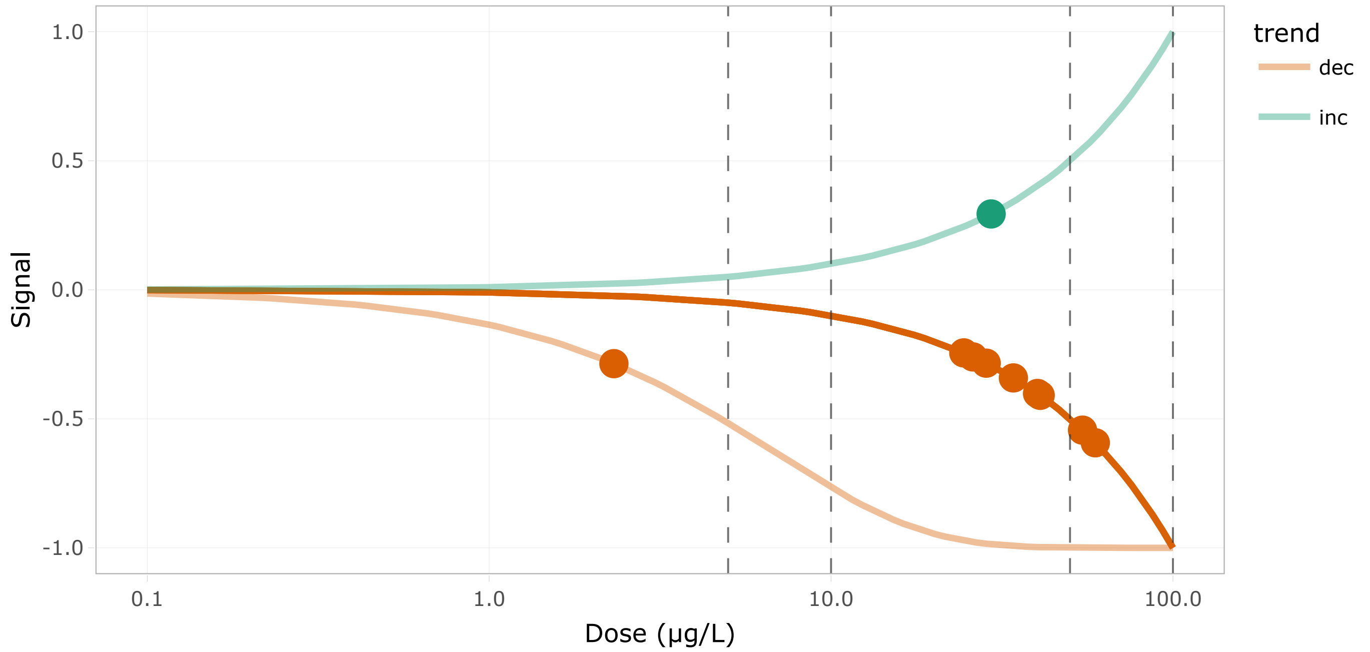


Figure 75: DR curves for the 37th cluster

Show 10 entries Search:

| | id | BMD.zSD | TF | trend | friendliness |
|---|---|---|---|---|---|
| | All | All | All | All | All |
| 1 | ago1 | 40.176 | true | dec | 1 |
| 2 | ago2 | 28.433 | true | dec | 1 |
| 3 | ago4 | 54.387 | false | dec | 1 |
| 4 | dicer1_g1t1 | 26.013 | false | dec | 1 |
| 5 | eri1_g2t1 | 29.392 | false | inc | 3 |
| 6 | pum1_t2 | 34.146 | false | dec | 1 |
| 7 | si:dkey-46g23.1_t1 | 40.9 | false | dec | 2 |
| 8 | snd1_t2 | 2.317 | true | dec | 1 |
| 9 | tarbp2 | 24.448 | false | dec | 1 |
| 10 | tnrc6c1_t4 | 59.285 | false | dec | 2 |

Showing 1 to 10 of 10 entries Previous 1 Next

Table 47: Table of the 37th cluster content

#### Cluster 9 - 10 transcripts (BMD q25 = 27.62 μg/L)

**Driver GO terms** : transcription by RNA polymerase I, transcription by RNA polymerase III

**KEGG pathways** : Cytosolic DNA-sensing pathway, RNA polymerase

**Wikipathways** :

Curvesplot BMDplot

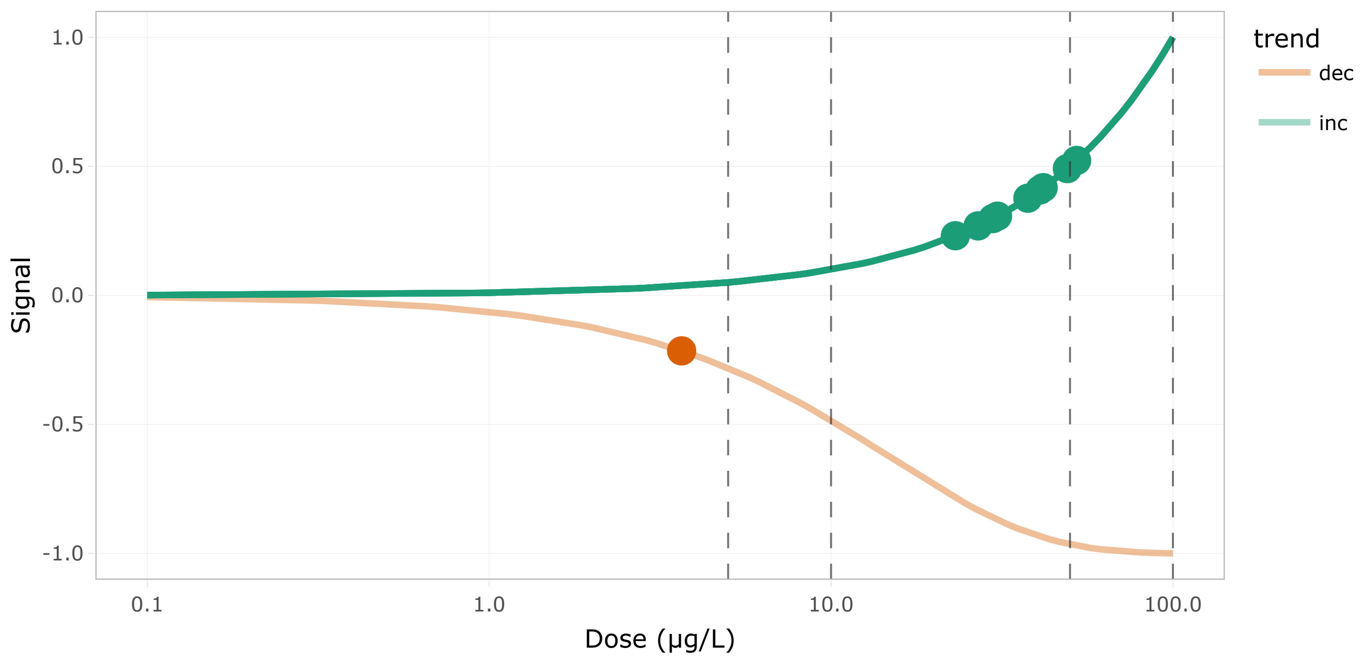


Figure 77: DR curves for the 38th cluster

Show 10 entries Search:

| | id | BMD.zSD | TF | trend | friendliness |
|---|---|---|---|---|---|
| | All | All | All | All | All |
| 1 | irf7_t2 | 23.102 | true | inc | 3 |
| 2 | polr1c_t2 | 40.827 | false | inc | 1 |
| 3 | polr2c_g2t1 | 30.679 | false | inc | 1 |
| 4 | polr2eb | 52.268 | false | inc | 1 |
| 5 | polr2f_g1t1 | 3.656 | false | dec | 1 |
| 6 | polr2h | 29.745 | false | inc | 1 |
| 7 | polr2l | 41.724 | false | inc | 1 |
| 8 | polr3k | 26.916 | false | inc | 1 |
| 9 | rrn3_t2 | 37.641 | true | inc | 1 |
| 10 | snapc1b | 49.114 | false | inc | 1 |

Showing 1 to 10 of 10 entries Previous 1 Next

Table 48: Table of the 38th cluster content

### Cluster 7 - 23 transcripts (BMD q25 = 27.65 µg/L)

**Driver GO terms** : nucleotide-excision repair

**KEGG pathways** : Nucleotide excision repair

**Wikipathways** : Estrogen signaling

Curvesplot BMDplot

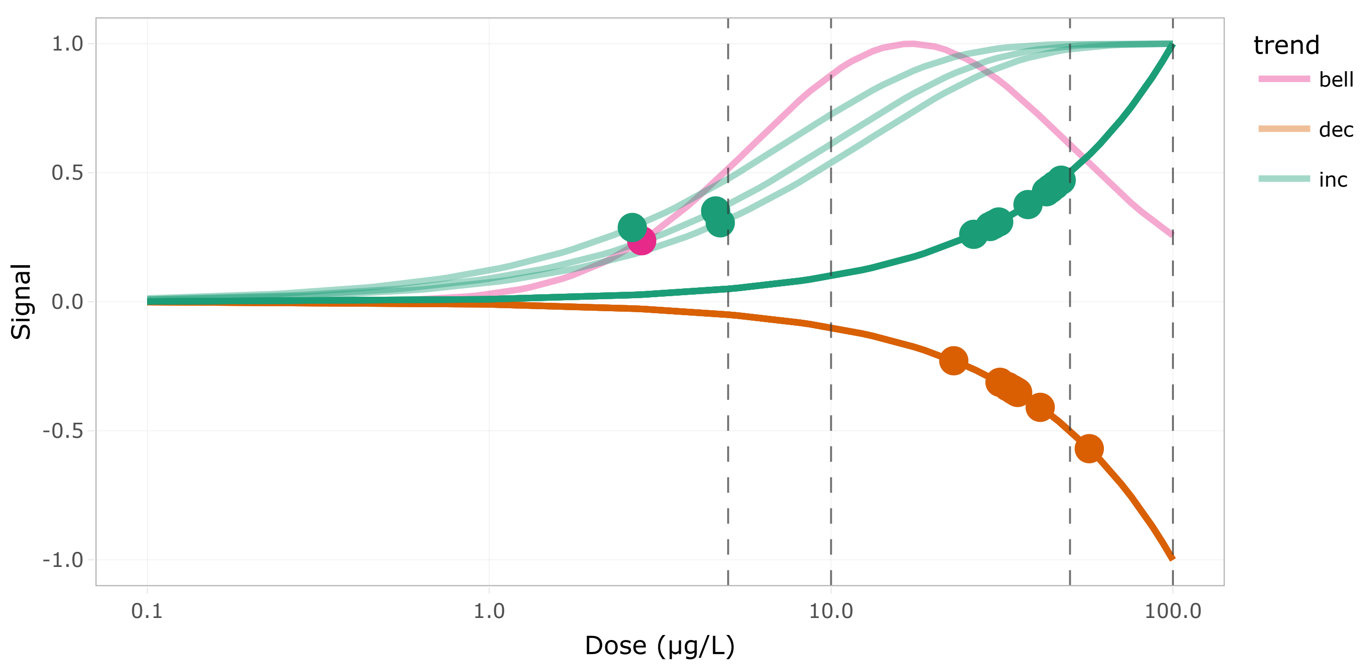


Figure 79: DR curves for the 39th cluster

Show 10 entries Search:

| | id | BMD.zSD | TF | trend | friendliness |
|---|---|---|---|---|---|
| | All | All | All | All | All |
| 1 | braf_t2 | 2.794 | false | bell | 4 |
| 2 | cdk12_t3 | 22.853 | false | dec | 1 |
| 3 | cdk8_g2t1 | 40.907 | true | dec | 1 |
| 4 | ercc3 | 37.679 | true | inc | 1 |
| 5 | ercc8_g1t2 | 4.596 | false | inc | 1 |
| 6 | gnas_t2 | 34.857 | false | dec | 2 |
| 7 | gnb1a | 56.928 | false | dec | 1 |
| 8 | gtf2e2_g2t3 | 43.667 | false | inc | 1 |
| 9 | gtf2h3 | 29.163 | true | inc | 1 |
| 10 | gtf2h4_t1 | 33.911 | false | dec | 2 |

Showing 1 to 10 of 23 entries

Previous 1 2 3 Next

Table 49: Table of the 39th cluster content

#### Cluster 24 - 5 transcripts (BMD q25 = 28.76 µg/L)

**Driver GO terms :**

**KEGG pathways :**

**Wikipathways :**

Curvesplot BMDplot

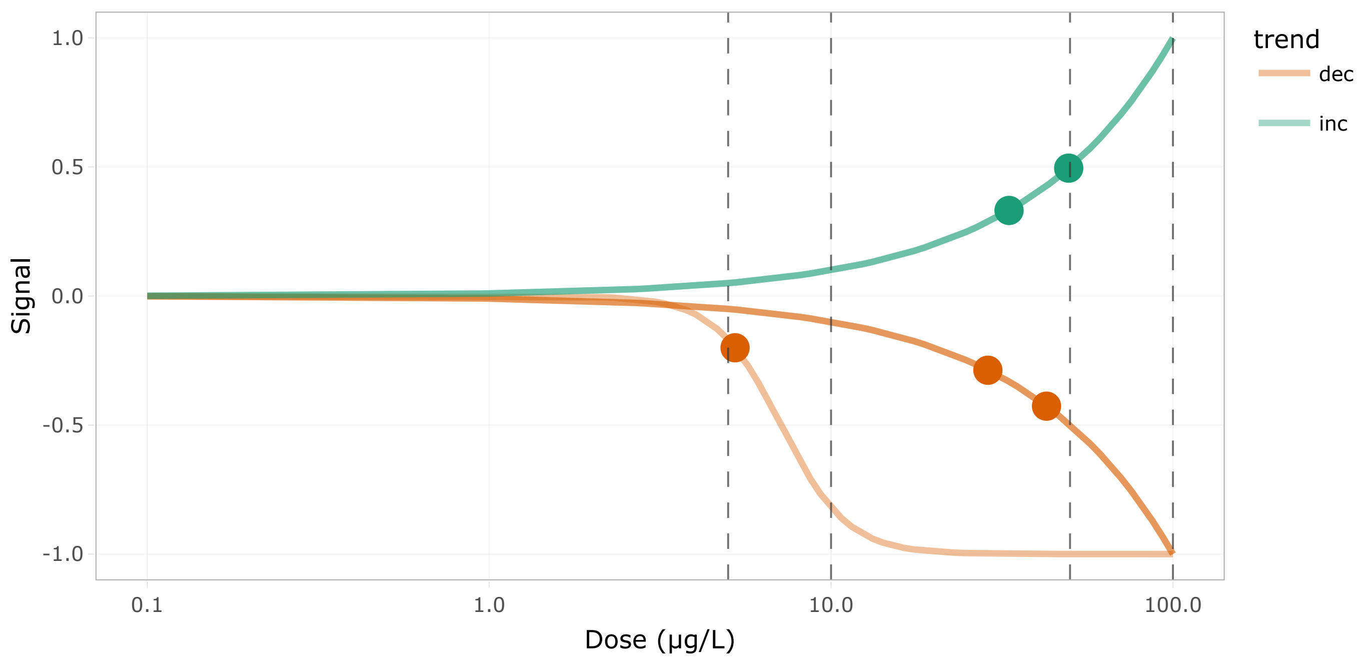


Figure 81: DR curves for the 40th cluster

Show 10 entries Search:

| | id | BMD.zSD | TF | trend | friendliness |
|---|---|---|---|---|---|
| | All | All | All | All | All |
| 1 | acta1b | 28.758 | false | dec | 1 |
| 2 | cttn_t2 | 5.238 | false | dec | 1 |
| 3 | pleca_t5 | 42.677 | false | dec | 1 |
| 4 | scinla | 33.157 | false | inc | 1 |
| 5 | tpma_t3 | 49.555 | false | inc | 1 |

Showing 1 to 5 of 5 entries Previous 1 Next

Table 50: Table of the 40th cluster content

#### Cluster 38 - 17 transcripts (BMD q25 = 29.77 µg/L)

**Driver GO terms :**

**KEGG pathways :** Nucleocytoplasmic transport

**Wikipathways :**

Curvesplot | BMDplot

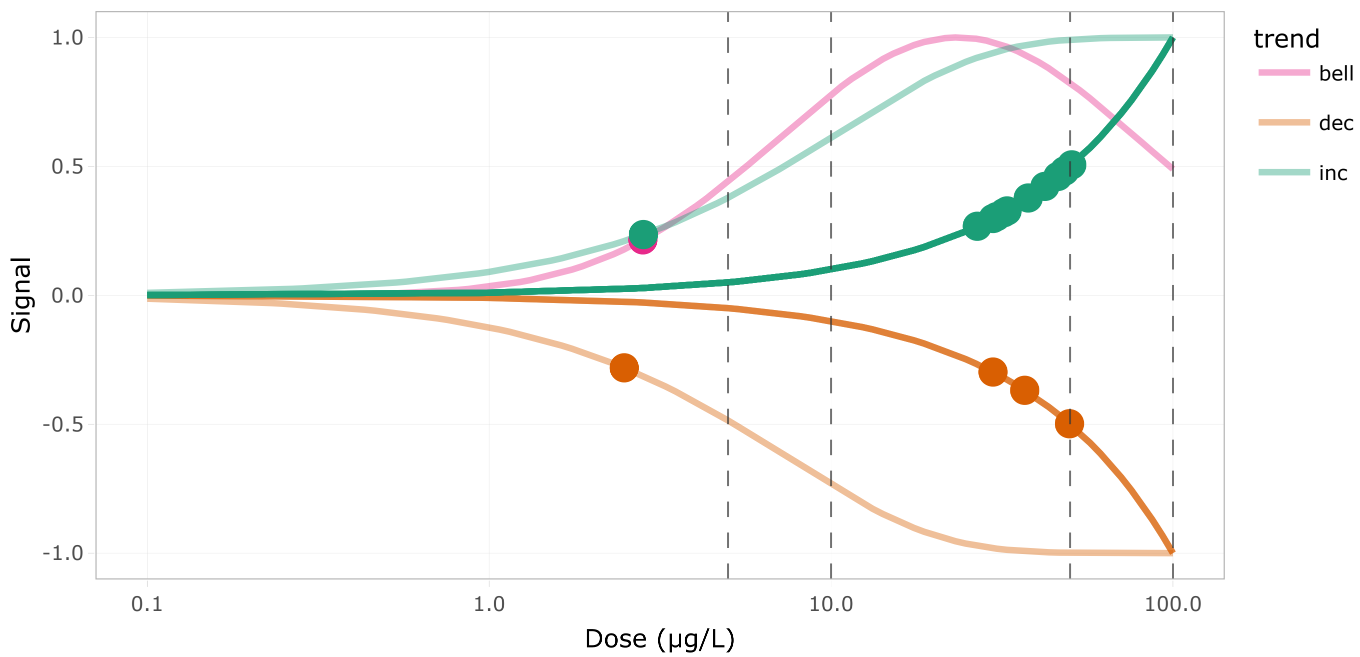


Figure 83: DR curves for the 41th cluster

Show 10 entries Search:

| | id | BMD.zSD | TF | trend | friendliness |
|---|---|---|---|---|---|
| | All | All | All | All | All |
| 1 | ahctf1_t2 | 48.302 | true | inc | 1 |
| 2 | eef1a1l1_t2 | 29.782 | false | inc | 2 |
| 3 | eef1a1l1_t3 | 49.818 | false | dec | 2 |
| 4 | kpna1 | 2.816 | false | bell | 2 |
| 5 | kpna3 | 2.483 | false | dec | 2 |
| 6 | kpna6_t2 | 36.855 | false | dec | 1 |
| 7 | nmd3 | 48.099 | false | inc | 3 |
| 8 | nup205_g1t3 | 42.231 | false | inc | 1 |
| 9 | nup37_t2 | 2.825 | false | inc | 1 |
| 10 | nup43_t3 | 37.734 | false | inc | 1 |

Showing 1 to 10 of 17 entries

Previous 1 2 Next

Table 51: Table of the 41th cluster content

#### Cluster 35 - 15 transcripts (BMD q25 = 31.17 µg/L)

**Driver GO terms :** ribosome biogenesis

**KEGG pathways :** Ribosome biogenesis in eukaryotes

**Wikipathways :**

Curvesplot BMDplot

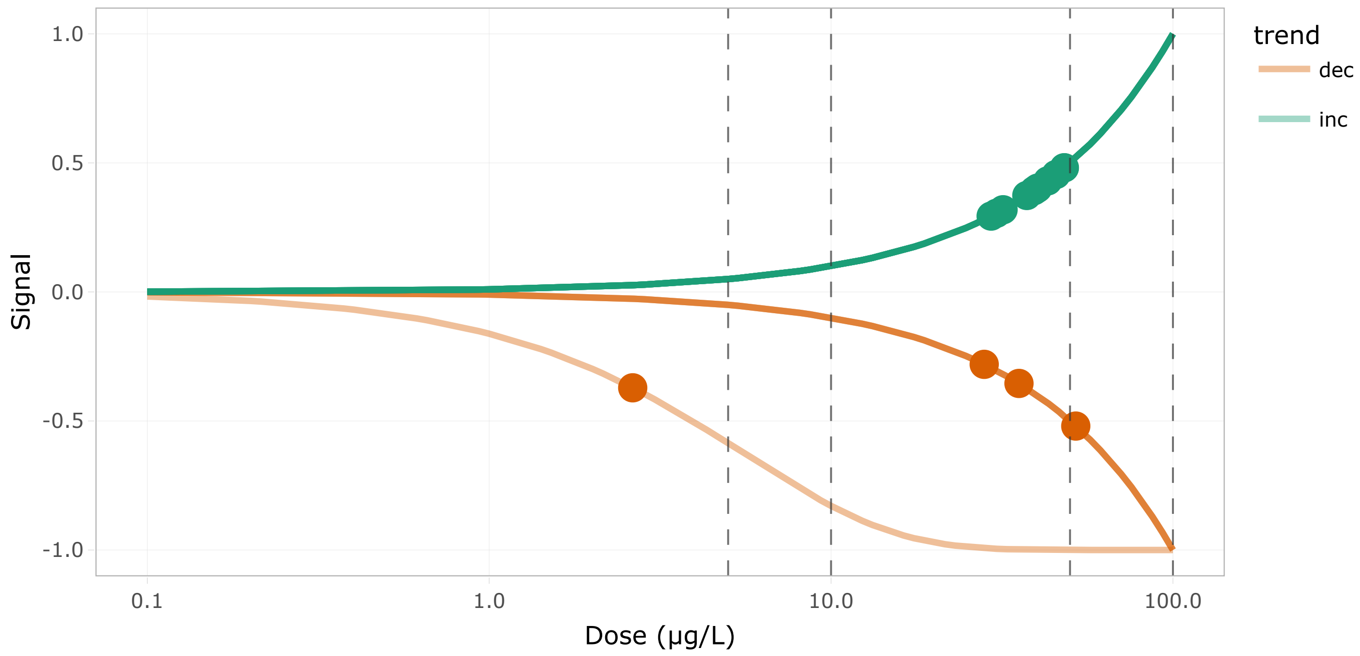


Figure 85: DR curves for the 42th cluster

Show 10 entries Search:

| | id | BMD.zSD | TF | trend | friendliness |
|---|---|---|---|---|---|
| | All | All | All | All | All |
| 1 | UTP14C | 51.973 | false | dec | 3 |
| 2 | emg1 | 48.122 | false | inc | 1 |
| 3 | eri1_g2t1 | 29.392 | false | inc | 3 |
| 4 | nip7 | 45.505 | false | inc | 2 |
| 5 | nmd3 | 48.099 | false | inc | 3 |
| 6 | noc4l | 43.046 | false | inc | 1 |
| 7 | nop58 | 2.629 | false | dec | 1 |
| 8 | pin4 | 37.377 | false | inc | 2 |
| 9 | ran | 30.488 | true | inc | 3 |
| 10 | rrp7a | 39.912 | false | inc | 1 |

Showing 1 to 10 of 15 entries Previous 1 2 Next

Table 52: Table of the 42th cluster content

#### Cluster 27 - 5 transcripts (BMD q25 = 31.66 µg/L)

**Driver GO terms :** mismatch repair

**KEGG pathways :**

**Wikipathways :**

Curvesplot BMDplot

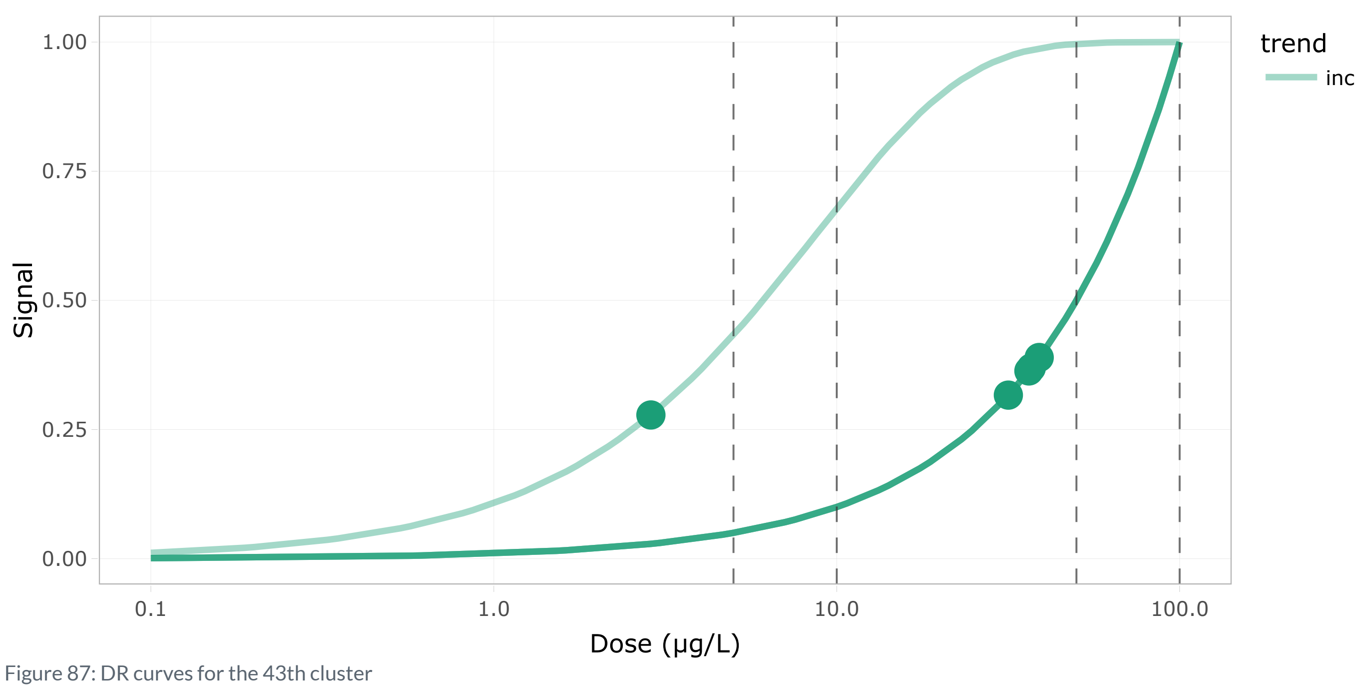


Figure 87: DR curves for the 43th cluster

Show 10 entries Search:

| | id | BMD.zSD | TF | trend | friendliness |
|---|---|---|---|---|---|
| | All | All | All | All | All |
| 1 | mlh1_t2 | 2.871 | false | inc | 1 |
| 2 | msh2 | 36.922 | false | inc | 1 |
| 3 | msh6 | 38.941 | false | inc | 1 |
| 4 | pms1_t2 | 36.337 | true | inc | 1 |
| 5 | recql4 | 31.657 | false | inc | 1 |

Showing 1 to 5 of 5 entries Previous 1 Next

Table 53: Table of the 43th cluster content

#### Cluster 51 - 4 transcripts (BMD q25 = 33.6 µg/L)

**Driver GO terms :**

**KEGG pathways :**

**Wikipathways :**

Curvesplot BMDplot

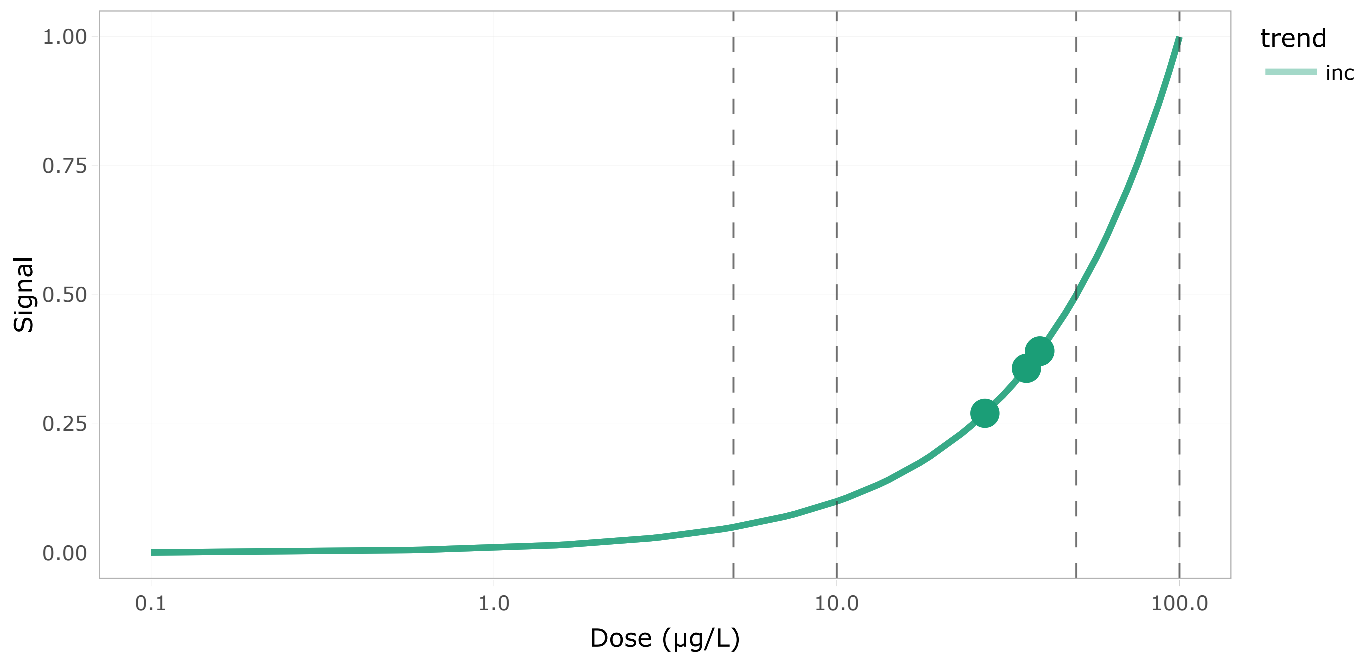

Figure 89: DR curves for the 44th cluster

Show 10 entries Search:

| | id | BMD.zSD | TF | trend | friendliness |
|---|---|---|---|---|---|
| | All | All | All | All | All |
| 1 | tomm22 | 39.032 | false | inc | 1 |
| 2 | tomm5 | 27.065 | false | inc | 1 |
| 3 | tomm6_t1 | 35.774 | false | inc | 1 |
| 4 | tomm7_t2 | 39.152 | false | inc | 1 |

Showing 1 to 4 of 4 entries

Previous 1 Next

Table 54: Table of the 44th cluster content
